# Longitudinal trajectories of brain development from infancy to school age and their relationship to literacy development

**DOI:** 10.1101/2024.06.29.601366

**Authors:** Ted K. Turesky, Elizabeth Escalante, Megan Loh, Nadine Gaab

## Abstract

Reading is one of the most complex skills that we utilize daily, and it involves the early development and interaction of various lower-level subskills, including phonological processing and oral language. These subskills recruit brain structures, which begin to develop long before the skill manifests and exhibit rapid development during infancy. However, how longitudinal trajectories of early brain development in these structures support long-term acquisition of literacy subskills and subsequent reading is unclear. Children underwent structural and diffusion MRI scanning at multiple timepoints between infancy and second grade and were tested for literacy subskills in preschool and decoding and word reading in early elementary school. We developed and implemented a reproducible pipeline to generate longitudinal trajectories of early brain development to examine associations between these trajectories and literacy (sub)skills. Furthermore, we examined whether familial risk of reading difficulty and children’s home literacy environments, two common literacy-related covariates, influenced those trajectories. Results showed that individual differences in curve features (e.g., intercepts and slopes) for longitudinal trajectories of volumetric, surface-based, and white matter organization measures were linked directly to phonological processing and indirectly to first-grade decoding and word reading skills via phonological processing. Altogether, these findings suggest that the brain bases of phonological processing, previously identified as the strongest behavioral predictor of reading and decoding skills, may already begin to develop by birth but undergo further refinement between infancy and preschool. The present study underscores the importance of considering academic skill acquisition from the very beginning of life.

**Significance Statement:** Reading is crucial for academic, vocational, and health outcomes, but acquiring proficient reading skills is a protracted developmental process involving lower-level subskills and brain structures that undergo rapid development starting in infancy. We examined how longitudinal trajectories of early brain development support long-term acquisition of reading using a reproducible pipeline we developed specifically for infant-to-school-age longitudinal MRI data. Findings suggest that the brain bases of reading-related skills begin to develop by birth but continue building between infancy and preschool. This study emphasizes the importance of considering academic skill acquisition as a dynamic process preceding the emergence of the skill, and it offers a roadmap for future studies to examine relationships between early brain development and academic skill acquisition.

## 1. Introduction

“…the best way to determine how a child learns is to follow them closely while they are learning.” (1, 2).

Reading acquisition is a multifactorial, developmental process that begins long before the skill manifests. Behaviorally, the acquisition of reading necessitates the acquisition and complex interplay of lower-level literacy subskills. In turn, these subskills represent waypoint products of brain development that began in utero. Therefore, understanding how reading skill emerges requires examining its behavioral subskills and the developmental trajectories of the brain areas subserving it starting from the very beginnings of life.

Literacy development represents a model process through which to examine academic skill acquisition, because literacy develops hierarchically, with lower-level “subskills” interacting and driving the emergence of higher-level academic skills (e.g., 3). For instance, phonological processing, which refers to the ability to detect, understand, and manipulate speech sounds (4–7), is the most consistent predictor of subsequent decoding and word reading (4, 8–10). Meanwhile, various studies have shown that both word reading and oral language skills, which encompass abilities supporting listening comprehension (11); e.g., vocabulary and syntactic knowledge), are important subskills for reading comprehension (12). However, these subskills themselves have protracted development and the brain structures that support them begin developing long before they manifest, with the most rapid development transpiring perinatally (13, 14).

Several cross-sectional and prospective studies have been conducted linking performance of these key literacy subskills to brain architecture. For instance, cross-sectional studies in preschoolers and kindergarteners have shown that phonological processing is associated with brain structure and function in left temporoparietal, occipitotemporal, and inferior frontal regions and tracts (15–20). This set of regions has also been linked with preschool and kindergarten oral language skills in some studies (21–23), but other studies have not shown associations (17, 24). Looking to early development, infant brain function and white matter organization have been shown to relate prospectively to (pre)school-age literacy outcomes (25–29), including phonological processing and oral language skills (30–32). Taken together, these studies suggest that certain brain areas may be important for academic skill *performance* at a particular time during development or as ‘neural scaffolding’ that supports subsequent language development (33). However, recent work in school-age children using large-scale, multi-site cross-sectional datasets showed little evidence for stable associations between individual differences in white matter organization and reading performance. Rather, using a separate longitudinal dataset, investigators found associations between slopes of white matter growth and reading gains, suggesting a dynamic interplay between brain development and learning to read (34). Moreover, the brain itself is a dynamic system that undergoes its most rapid development during infancy and early childhood (13, 14), making cross-sectional and prospective designs suboptimal to capture individual differences in early skill *acquisition* (2) As such, individual longitudinal trajectories, which capture heterogenous rates of brain and skill development between infancy and second grade, are needed to examine the acquisition of (sub)skills important for literacy.

Therefore, the overall goal of the current study is to examine the relationship between longitudinal trajectories of early brain development and acquisition of reading-related subskills, which we undertook in four objectives. The first objective was to generate individual longitudinal trajectories of early brain development spanning infancy to school age. Prior longitudinal studies have examined trajectories of early brain development but only up to the second year of life (e.g., 35), while others have examined developmental trajectories of brain structure spanning infancy to school age but only by harmonizing cross-sectional and longitudinal datasets acquired and processed with varying methods (e.g., 36). However, to our knowledge, no purely longitudinal studies have mapped the development of brain structure, including white matter organization, from infancy to school age, using a consistent processing pipeline and one that is appropriate for early development. Indeed, methodological challenges associated with infant MRI data have restricted the number and breadth of longitudinal studies in the early developmental period. This is not only because acquiring and processing infant MRI data requires specialized procedures and tools to generate accurate brain estimates (37–42), but also because the use of different procedures and tools for different developmental stages can introduce bias in developmental analyses. However, the alternative—to use identical procedures and tools for all developmental stages—would likely cause estimates to vary in accuracy across ages and could introduce spurious effects and/or obscure true effects in developmental comparisons (43). Some infant-specific tools offer age-specific adjustments (e.g., different sets of templates) within the first two years of life, and the small number of studies that have examined longitudinal trajectories of brain development in the first two years have opted for a balanced approach, with age-appropriate adjustments to limited processing steps in a way that does not require fundamentally different techniques for different ages (e.g., additional sequences or different smoothing kernels; (35, 44). However, methods to accommodate a wider early developmental range are lacking.

In the current study, we extend this work by developing reproducible pipelines for generating longitudinal trajectories of volumetric, surface-based, and white matter organization brain measures from infancy to school age. To do this, we leveraged both infant-specific and standard MRI procedures and tools. We then compared among several candidate linear and nonlinear mixed effects models, varying according to function (e.g., linear, logarithmic) and random parameters (e.g., intercepts alone versus intercepts and slopes), to identify the model with the most parsimonious fit (45). In general, prior longitudinal studies have reported rapid growth at birth that tapers with age (35, 46–48), except for cortical thickness, which peaks between ages one and two years (49), suggesting logarithmic functions might provide a better fit compared with other functions for most measures.

Our second objective was to examine the association of individual differences in early brain development to long-term literacy development. As such, we extracted curve features (e.g., intercepts and slopes) from individual longitudinal trajectories for brain areas and tracts previously shown to relate to reading-related skill development in longitudinal studies in older children (34, 50–59) or prospective studies in infants (30–32). We then tested these curve features for correlations with preschool/early kindergarten phonological processing, a key literacy subskill. Based on converging evidence from prior neuroimaging (30–32) and genetics studies (60, 61), we hypothesized that phonological processing skill would relate to curve intercepts (i.e., brain estimates at birth) and slopes (i.e., rate of brain development). If the foundations of literacy development are largely present at birth and stable across early development, then we expect most of these brain-behavior associations to be with curve intercepts, whereas if the brain development subserving literacy subskills is more protracted or dynamic, as suggested by Roy and colleagues (34), then we expect brain-behavior associations to be with curve slopes.

Our third objective was to examine the roles of risk factors related to literacy skills in shaping longitudinal trajectories of early brain development. Reading difficulty is heritable, as 40-60% of children with a familial risk (e.g., first-degree relative with a history) of reading difficulty (FHD+) themselves develop reading difficulty (62, 63) Brain imaging studies of FHD+ preschoolers show reduced gray matter volume, activation, and fractional anisotropy in left occipito-temporal and temporo-parietal regions (16–18) and tracts (57) compared with FHD-preschoolers; these reductions overlap with those observed in children with reading difficulty (20, 64–70), suggesting that the phenotypes characteristic of reading difficulty manifest in some children before the start of formal reading instruction. Concordantly, similar alterations have been observed earlier in development, where FHD+ infants exhibited lower fractional anisotropy in left arcuate fasciculus compared with FHD-infants (71), distinguishable patterns of functional connectivity in left fusiform gyrus as shown by a support vector machine classifier (72), and alterations in neural responses to basic speech sounds as measured by event-related potentials (73–75). Possibly underlying this heritability, work in genetics, including recent large-scale genome-wide studies, has shown that reading-related skills (60) and reading difficulty (76, 77) are associated with variation in genes involved in early developmental processes, including neurogenesis and axon guidance (78).

Another risk factor repeatedly shown to affect children’s literacy skills is the home literacy environment, which includes caregiver–child shared reading and reading-related resources, in preschool literacy skills (79–84) and in infancy/toddlerhood (83, 85–90). Recent work has also identified links between the home literacy environment and brain architecture in preschoolers/kindergarteners (21, 24, 91–94) and infants (95, 96). Overall, these studies offer strong evidence that genetic and environmental factors related to the development of reading-related skills are likely to affect longitudinal trajectories of brain development starting perinatally. Therefore, we hypothesized that the FHD status and home literacy environment would significantly contribute, as covariates, to longitudinal trajectories of brain development in left hemisphere temporo-parietal, occipito-temporal, and inferior frontal regions and tracts.

Our fourth objective was to broaden the scope of our examination of early brain-literacy associations to other literacy-related subskills and subsequent reading skills. Reading acquisition is a hierarchical process in which multiple distinct but interacting subskills (including but not limited to phonological processing) converge to effect higher-order skills. First, we tested the specificity of associations with phonological processing by examining another subskill important for reading comprehension, oral language skill. Second, we tested whether phonological processing mediated relationships between curve features of early brain development and subsequent measures of decoding and word reading skill. Taken altogether, findings from this study will inform our understanding of how early brain development contributes to later reading-related skill acquisition.

## 2. Methods

### 2.1. Participants

Children examined in this study were obtained from two longitudinal cohorts: the New England dataset and the Calgary open-source dataset (https://osf.io/axz5r/). Only children with high-quality (i.e., post-QC, please see below) MRI data from at least two timepoints between birth and late childhood were included. Due to the multi-modal nature of the study, varying numbers of longitudinal datasets were used for structural (n = 98 with 276 observations) versus diffusion analyses (n = 128 with 396 observations). For the New England cohort, family history (i.e., first-degree relative with a history) of reading difficulty was also examined and present in 40 of 80 children. For children ages ≤ 24 months, demographic information was provided by caregivers. Please see Table 1 for overall demographic details, Supplementary Table 1 for demographic details by modality, and Supplementary Methods for participant information, including race demographics, unique to each cohort. This study was approved by the Institutional Review Boards of Boston Children’s Hospital (IRB-P00023182), Harvard University (IRB21-0916), and the University of Calgary Conjoint Health Research Ethics Board (REB13-0020). Participants’ parents gave informed consent, and children gave verbal assent if over 25 months old. Data examined here partially overlap with brain images analyzed in previous studies examining infant brain development (30–32, 36, 48, 54, 71, 72, 96–102).

**Table 1.**
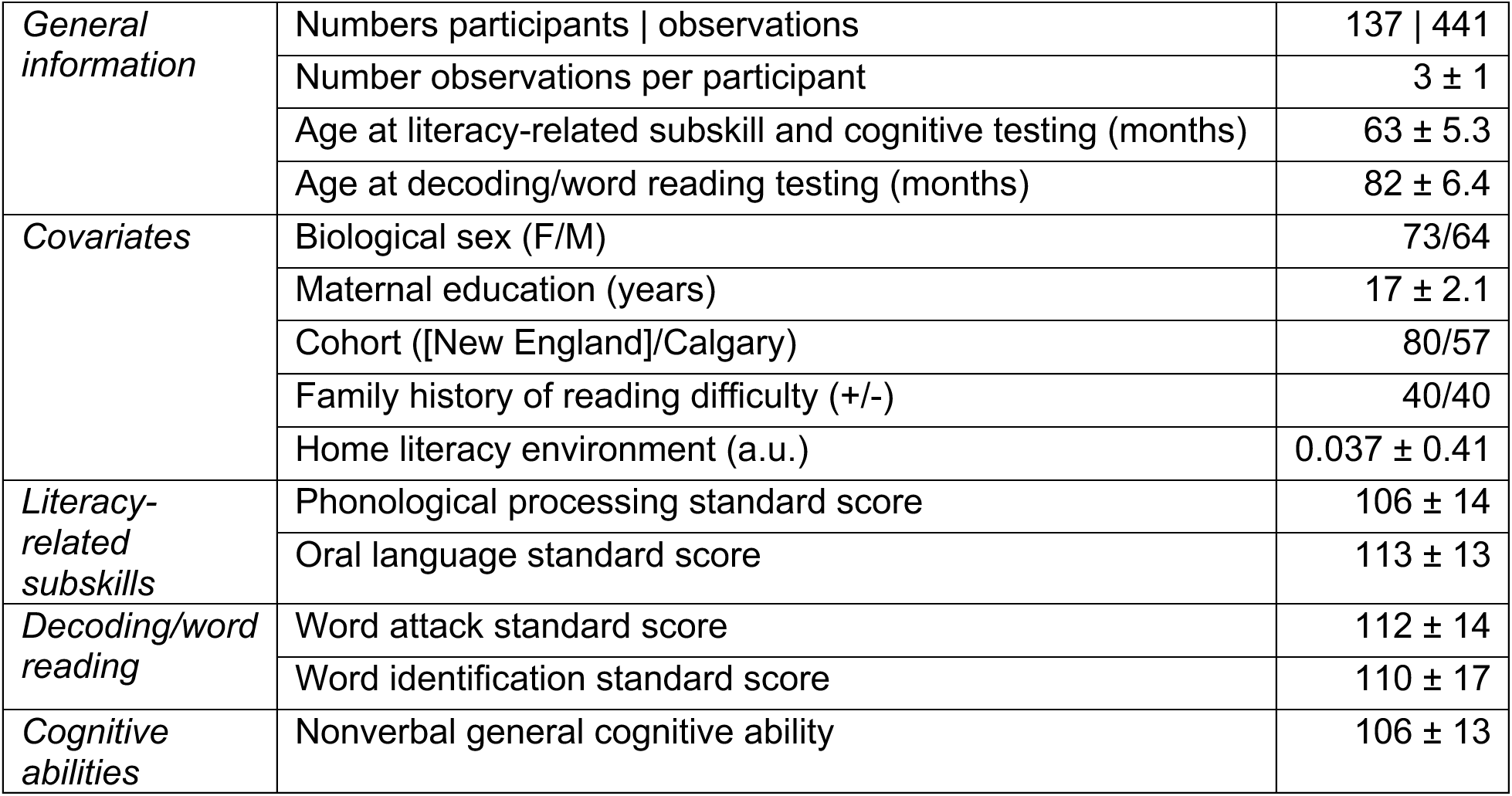
Participant Demographics.

### 2.2. Environmental variables

Socioeconomic status was measured with maternal education, consistent with prior brain imaging studies on socioeconomic status (103–108). Maternal education measures were collected during each timepoint, although inter-time-point variability was low, and these were averaged to generate one socioeconomic status measure across the developmental window. Rather than using ordinal coding, years of education as a continuous measure ranging from 12 to 20 years were used.

For the New England cohort, parents also completed at each time point questionnaires relating to children’s home literacy environments home literacy environment, which includes the extent of parent-child shared reading and access to reading-related resources (109). Responses were indicated using ordinal scales ranging from 1 to 6. As responses were non-normally distributed (p < 0.05 according to the Shapiro-Wilk normality test; Supplementary Figure 1), except for “Time read to per week” for all timepoints and “Frequency with which family members share rhymes or jokes with the child” for one timepoint, they were normalized and then averaged at each timepoint according to procedure used previously (30). As with maternal education, home literacy environment estimates, which exhibited low inter-timepoint variability, were averaged to generate one estimate per individual across the developmental window. These overall home literacy environment estimates, which were normally distributed (Shapiro-Wilk W = 0.99, p > 0.05; Supplementary Figure 2), were used in later statistical analyses.

### 2.3. Literacy and cognitive measures

Two literacy subskills were administered to children prior to the beginning of formal reading instruction: phonological processing and oral language. These constructs were selected as representative subskills supporting literacy development (3, 4, 8–10, 12, 110); however, they constitute a small subset of literacy-related measures collected for this cohort. No outliers were detected using the *isoutlier* function in MATLAB, which sets an outlier threshold at three scaled median absolute deviations from the median.

Phonological processing was measured in both New England and Calgary cohorts. For the New England cohort, the *phonological processing* composite was estimated from three subtests from the WJ-IV Tests of Cognitive Abilities: word access, word fluency, and substitution (111). Word access measures phonetic coding by asking children to identify words containing certain sounds. Word fluency measures speed of lexical access by asking children to name as many words as possible beginning with a certain sound in one minute. Substitution measures children’s ability to produce a new word by replacing one sound from a provided word with another sound. Importantly, while the word access and word fluency subtests require some level of lexical access, these subtests, along with the substitution subtest, are well established measures of phonological/phonemic processing/awareness. For additional details and item examples from the technical manual, please see the Supplementary Methods. Children in the Calgary cohort were administered the phonological processing subtest of the NEPSY-II, which measures phonemic awareness (112). Unlike the New England cohort, each child from the Calgary cohort completed this subtest multiple times. To harmonize phonological processing scores across the two datasets, we used NEPSY-II scores from when the child was closest in age to the average age at which the New England cohort completed the WJ-IV (64 months) and not earlier than 50 months of age.

The composite oral language, which was only measured in the New England cohort, was estimated from two subtests from the WJ-IV Tests of Oral Language: picture vocabulary and oral comprehension (113). Picture vocabulary measures lexical knowledge by asking children to specify a picture corresponding to a given word or naming an object. Oral comprehension measures oral listening, vocabulary, and reasoning by asking children to identify missing words from short passages. All assessments were administered and double-scored by testers trained by a clinical psychologist and then raw scores were converted to standard scores.

In addition, we measured decoding and word reading with two untimed subtests from the Woodcock Reading Mastery Tests III (114): word attack and word identification. For word attack, children were presented with pseudowords that they needed to *decode* using phonological abilities. For word identification, children were presented with individual real words that they needed to *read*. Word attack and word identification subtests were administered at the beginning of formal reading instruction.

Lastly, we measured children’s nonverbal general cognitive ability at preschool/early kindergarten-age using the Matrix Reasoning subtest of the Kaufman Brief Intelligence Test: 2nd Edition (KBIT-2, (115)). Herein, children were asked to identify the piece missing from a matrix of visual images.

All raw estimates for each of the WJ-IV, NEPSY-II, and WRMT subtests were non-normally distributed (p < 0.05 according to the Shapiro-Wilk normality test), except for preschool/early kindergarten picture vocabulary and late kindergarten/grade 1 word attack (Supplementary Figure 3). All standardized (composite) estimates used in subsequent analyses were normally distributed (p > 0.05; Supplementary Figure 4).

### 2.4. MRI data acquisition and processing

All data were acquired on a 3.0 T scanner with a 32-channel head coil. Please see Supplementary Methods for cohort-specific acquisition parameters.

The rapid brain growth transpiring immediately after birth posed serious challenges for structural MRI processing because standard methods that are optimal for older children are suboptimal for infants and vice versa (43). To circumvent bias associated with choosing a single pipeline for multiple developmental stages, we implemented pipelines that are age-appropriate but not fundamentally different for each developmental stage.

#### 2.4.1. Structural MRI processing and quality control

Raw magnetization-prepared rapid gradient-echo (MPRAGE) images were visually inspected for artifacts by a trained rater and scored as “fail,” “check,” or “pass” (116). Only images scored as “check” or “pass” underwent image processing. The standard, unmodified FreeSurfer v7.3 (https://surfer.nmr.mgh.harvard.edu/) “recon” pipeline was used for brains > 50 months. Processing procedures for brains < 50 months were similar in concept to those described in (117). Namely, Infant FreeSurfer (for brains ≤ 24 months) or standard FreeSurfer (for brains 25 to 50 months) was used to extract the brain from the skull, correct for intensity inhomogeneity, and segment MPRAGE images by tissue class (gray matter, white matter, cerebrospinal fluid) and subcortical brain region (40); https://surfer.nmr.mgh.harvard.edu/fswiki/infantFS). To improve tissue classification accuracy, MPRAGE images were submitted in parallel to iBEATv2.0 Docker 1.0.0 (118–120); https://github.com/iBEAT-V2/iBEAT-V2.0-Docker), has been validated for birth to age six years (118). Resulting segmentations from each software package were then hybridized using in-house MATLAB code that combined the cortical pial and white matter boundaries labelled by iBEATv2.0 with the subcortical parcellations of Infant FreeSurfer (brains ≤ 24 months) or standard Freesurfer (brains 25 to 50 months), effectively relabeling cortical gray and white matter in the FreeSurfer-style segmentation; for details, please see the Supplementary Methods. Structural processing was finalized by submitting these FreeSurfer-style hybrid segmentations to a modified version of the standard FreeSurfer v7.3 “recon” pipeline, which, for brains ≤ 24 months, incorporated elements from Infant FreeSurfer. For a schematic of the structural processing pipeline, please see Supplementary Figure 5.

Resulting white and pial surfaces were visually inspected by two (brains > 50 months) or three (brains ≤ 24 months) trained raters on a 3-point scale (0, 1, and 2) and datasets with average ratings > 1.5 were retained for subsequent analyses. Finally, parcellations were visualized to ensure accuracy of anatomical labels. For reproducibility purposes, we did not perform manual editing to correct tissue mislabeling in the remaining images; however, structural measures from edited and unedited pediatric brain images processed with FreeSurfer have been shown to be highly correlated (121), including in children (122). Measures of gray and white matter volume, surface area, cortical thickness, and mean curvature, generated in the final FreeSurfer steps, were extracted from 8 a priori left hemisphere regions delineated with the Desikan-Killiany atlas— banks of the superior temporal sulcus, fusiform gyrus, inferior parietal lobule, middle temporal gyrus, pars opercularis, pars triangularis, superior temporal gyrus, and supramarginal gyrus— based on their reported involvement in reading-related subskills (15–23, 30–32, 54, 60; for reviews, please see 123, Table 3 and 124).

#### 2.4.2. Diffusion MRI processing and quality control

Preprocessed and tractography for all diffusion-weighted image (DWI) data, regardless of age, were performed with MRtrix3 based on the pipeline established for the Developing Human Connectome Project (41, 125). DWI data were first denoised using Marchenko–Pastur principal component analysis (126–128) and then corrected for susceptibility distortions, eddy currents, motion, and intensity inhomogeneity using FSL’s topup and eddy (with slice-to-volume correction) functions (129–133), and Advanced Normalization Tools (ANTs) N4 bias correction tool (134). Subsequently, three tissue response functions for spherical deconvolution were estimated using the Dhollander algorithm, a 0.1 fractional anisotropy threshold, and eight maximum harmonic degrees (135). Fiber orientation densities (FODs) were computed with multi-shell, multi-tissue constrained spherical deconvolution (136, 137) and then normalized using multi-tissue informed log-domain intensity normalization.

Two million streamlines were tracked from the normalized FOD maps using the Anatomically Constrained Tractography (ACT) technique. Importantly, ACT has been shown to greatly improve tractography, but it relies on accurate tissue segmentations, which are typically challenging to generate with infant brain data. Using the hybrid segmentations generated for brains ≤ 50 months (please see above) circumvented this challenge. Thus, FreeSurfer-style hybrid segmentations for brains ≤ 50 months and standard FreeSurfer segmentations for brains > 50 months were registered to the preprocessed DWI images using ANTs and then converted to five-tissue-type images for ACT. The remaining whole-brain tractography parameters included seeding at the gray/white matter boundary and tracking with the iFOD1 probabilistic algorithm; step size, minimum and maximum length, and maximum step angle were set to default (138, 139).

Resulting whole-brain tractography was then submitted to the open-source instantiation of Automated Fiber Quantification (AFQ; (140) for waypoint- and probabilistic-atlas-based fiber tract segmentation. The standard pyAFQ pipeline was used for brains > 24 months (141), whereas pyBabyAFQ was used for brains ≤ 24 months (142). Please see Supplementary Methods for a summary of differences between the two AFQ instantiations. Next, tracts were resampled to 100 equidistant nodes, and diffusion properties (fractional anisotropy; mean diffusivity) were quantified for each node for the left hemisphere tracts of interest—arcuate fasciculus, superior longitudinal fasciculus, and inferior longitudinal fasciculus. Mean fractional anisotropy and mean diffusivity values for each tract were used to examine model fits (please see section 2.5.1), whereas a more fine-grained, node-based approach was taken for brain-behavior analyses (please see section 2.5.2). For the latter, to better align tract cores across participants, five nodes on either end were removed, reducing the total number of nodes per tract to 90. Finally, tracts of interest were visually inspected by two trained raters on a 3-point scale (0, 1, and 2) and datasets with average ratings ≥ 1 were retained for subsequent analyses. For a schematic of the diffusion processing pipeline, please see Supplementary Figure 6.

### 2.5. Statistical analyses

An overview of statistical operations, all of which were performed in RStudio version 4.2.2, can be found in Supplementary Figure 7.

#### 2.5.1. Longitudinal trajectory estimation

Prior to modeling, estimates of brain structure and white matter organization underwent a final quality control procedure to remove estimates for brain areas/tracts of interest if, for brains > 25 months, they were preceded or followed by ≥ 10% annual change (positive or negative) in any measure and for brains ≤ 24 months, inter-observation changes were negative (or positive for mean diffusivity). To mitigate data loss, we next identified which observation—earlier or later— was more likely to be inaccurate, using an outlier detection procedure (described in the Supplementary Methods). All remaining participants after these procedures had multiple observations.

To generate longitudinal trajectories (i.e., growth curves), we next submitted cleaned, longitudinal structural and white matter organization estimates to linear mixed effects models using linear, logarithmic, and quadratic functions from the R ‘lme4’ package. Intercepts and slopes were modeled as fixed and random effects, and covariates for biological sex, socioeconomic status, and cohort (New England or Calgary) were entered as fixed effects. Vijayakumar and colleagues recommends comparing model fits quantitatively (45); consequently, we computed Bayesian Information Criterion (BIC) metrics and the model (i.e., function and number of random terms) that provided the best fit (i.e., lowest BIC value) was selected for subsequent analyses.

Correlations between intercepts and slopes were tested to ensure associations between random terms were minimized. Most curve features were normally distributed according to the Shapiro-Wilk normality test (p > 0.05) and histograms depicting their distributions are provided in Supplementary Figures 9-15.

#### 2.5.2. Associations between longitudinal trajectories of early brain development and phonological processing

Next, individual-level curve features (i.e., intercepts and slopes) were extracted and tested for correlations (Pearson) with literacy subskills. The extracted curve features represent individual variation in intercepts or slopes beyond what would be predicted based on the covariates (e.g., biological sex); consequently, we did not include these same covariates in the tests of correlations. To determine whether curve features of volumetric and surface-based measures were associated with phonological processing in the full sample after accounting for multiple brain regions (8 tests), a significance threshold was set to p_FDR_ < 0.05 and applied to measures (e.g., gray matter volume) separately. As diffusion analyses were performed node-wise (90 nodes), we corrected for multiple comparisons at p_FWE_ < 0.05 using a permutation-based, threshold-free cluster enhancement method (143), implemented in the permuco package in R (144); corrections were performed separately for each tract. These methods are similar to those used previously (24, 96, 108). When significant, individual growth curves were separated into three groups according to scores on their behavioral assessment with low < 85, 85 ≤ average ≤ 115, and high >115, averaged by group, and then plotted.

#### 2.5.3. Sensitivity analyses

We conducted four sensitivity analyses to test the reliability of our results. First, we performed a replication analysis on gray/white matter volume, surface area, and mean diffusivity with nonlinear mixed effects models using asymptotic functions from the R ‘nlme’ package (145), similar to that described in Alex and colleagues (36); https://github.com/knickmeyer-lab/ORIGINs_ICV-and-Subcortical-volume-development-in-early-childhood). Intercepts and asymptotes were modeled as fixed and random effects; rate constants were modeled as fixed effects. Second, instead of testing correlations between curve features and outcomes, we entered outcomes as main and interaction terms in linear mixed effects models. Third, adhering to recommendations to report both raw and TIV-corrected results (45), we recomputed brain-behavior associations for volumetric and surface-based measures using semipartial correlations (Pearson) with the random terms from longitudinal modeling with TIV as covariates of no interest. Fourth, we submitted volumetric and surface-based brain-behavior associations to semipartial correlations (Pearson) with average (across timepoint) Euler numbers, which quantifies topological defects (146). For additional details on these sensitivity analyses and Euler quantification, please see the Supplementary Methods.

#### 2.5.4. Specificity analyses

To determine whether brain-behavior effects were specific to certain brain measures, regions, curve features, and behavioral outcomes, we performed three specificity analyses. First, we generated whole-brain maps depicting variability in associations with phonological processing according to brain measures (e.g., gray matter volume), region, and curve feature. Second, we examined whether brain-behavior associations for diffusion measures persisted in right arcuate fasciculus, superior longitudinal fasciculus, and inferior longitudinal fasciculus. Third, limited to the New England children, we tested correlations between curve features of brain development and oral language skill, another reading subskill, and nonverbal general cognitive abilities. For subsequent testing of literacy-related covariates, we also reanalyzed brain-behavior associations with phonological processing in the New England subsample only. For volumetric and surface-based measures, FDR correction accounted for brain regions and behavioral measures (24 tests). For diffusion, cluster-level FWE correction accounted for nodes (90 tests).

#### 2.5.5. Literacy-related factors as fixed effects in models of early brain development

We determined whether literacy-related covariates (i.e., home literacy environment and FHD status) contributed to the fit of the growth curve in two ways. First, we examined each covariate as fixed effects in models of brain development for measures and regions whose curve features related to phonological processing (please see brain-behavior associations in 2.5.2). For brain structure, a significance threshold of p_FDR_ < 0.05 was applied to correct for the multiple brain development models that fit this criterion. Diffusion analyses, which were performed node-wise for each tract separately (90 tests), were corrected for multiple comparisons using a lenient cluster size threshold of 5 nodes, as threshold-free cluster enhancement method was not available for the statistical tests applied to covariates. Second, we compared brain development models with and without each covariate using BIC metrics. For diffusion, BIC metrics were obtained using tract averages (rather than separately on individual nodes). Correction for multiple comparisons was again set to p_FDR_ < 0.05 and applied as described above for brain structure, this time also including statistics for average diffusion measures.

#### 2.5.6. Indirect effects between longitudinal trajectories of early brain development and word reading outcomes via literacy subskills

As with previous work (147), indirect effects were tested when literacy subskills were related to curve features of brain development and decoding and word reading outcomes (after FDR correction). Formal indirect effects modeling was conducted using the Mediation package in R with 20,000 bootstrapped samples, and effects were determined significant when the 95% confidence interval for the average causal mediation effect did not include 0. All mediation models included covariates for maternal education, home literacy environment, and family history of reading difficulty. As in the sensitivity analyses, models were run twice, once without controlling for TIV and once including TIV curve features. Because mediation testing was limited to data with significant (after correction for multiple comparisons) brain-behavior associations, no additional correction for multiple correction was applied; this is consistent with prior literature (36, 147, 148).

2.6. Data availability

All code used to process and analyze has been made openly available at https://github.com/TeddyTuresky/Longitudinal-Trajectories-Early-Brain-Development-Language. The Calgary dataset is also freely available at https://osf.io/axz5r/. All Boston/Cambridge data used in this study will be made publicly available at upon acceptance of the manuscript.

## 3. Results

### 3.1. Longitudinal trajectories of brain structure and white matter organization in left hemisphere literacy-related brain regions and tracts

Our first objective was to generate accurate longitudinal trajectories of early brain development in regions and tracts reportedly involved in reading-related subskills (15–23, 30–32, 54, 60; for reviews, please see 123, Table 3 and 124). Accordingly, growth curves for regional estimates of gray and white matter volume, surface area, cortical thickness, and mean curvature and for tract estimates of fractional anisotropy and mean diffusivity were generated from 137 children (F/M = 73/64) with 441 observations. Every child had at least two observations between birth and school age with 77 having at least one observation in infancy (Figure 1; please see Supplementary Figure 16 for age distributions by modality). Linear mixed effects models included individual-level intercepts and slopes as well as covariates for biological sex, socioeconomic status, and cohort. All structural and diffusion measures for all regions and tracts were best fit with logarithmic functions according to Bayesian Information Criterion (Supplementary Table 2), whereby rates of development were steeper perinatally and then slowed across the first ten years following birth (Figure 2; Supplementary Figures 17-22).

**Figure 1.**
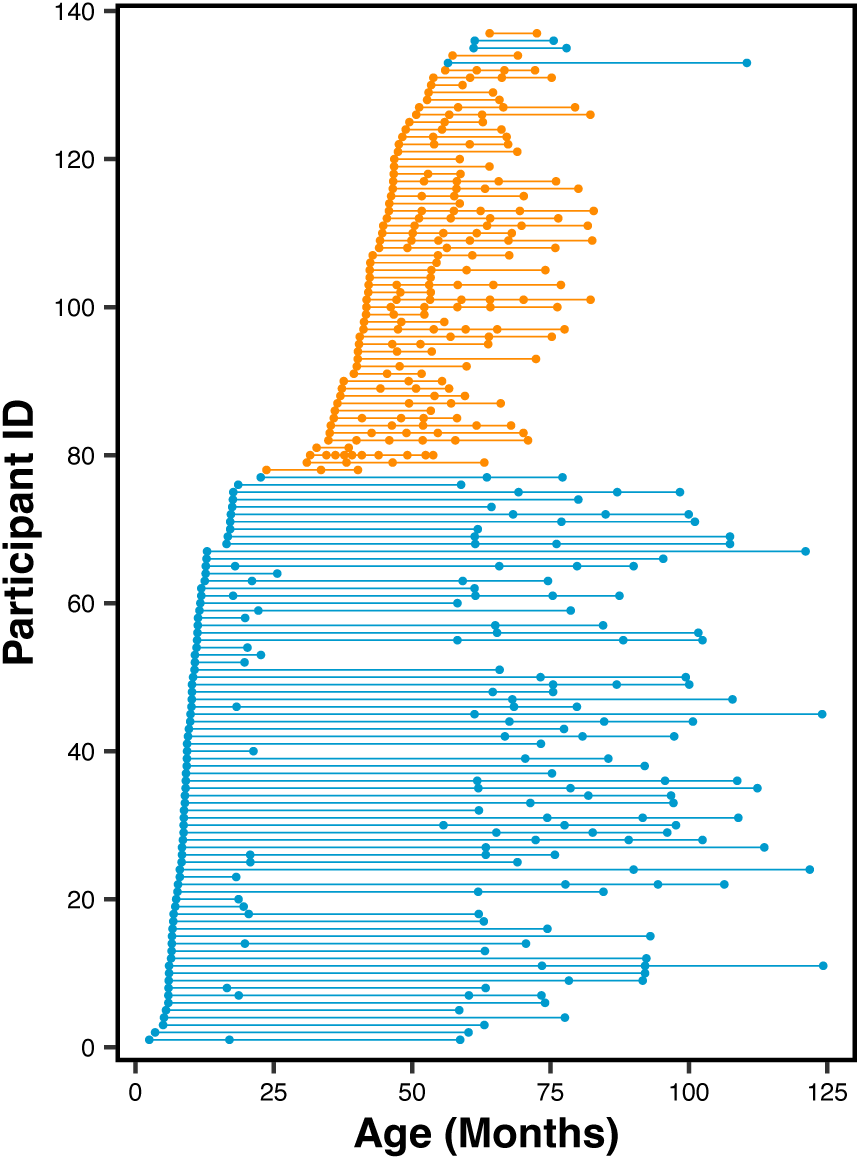
Age distribution of longitudinal dataset from infancy to late childhood. All children had structural and/or diffusion MRI data from at least two observations (dots). Blue, New England cohort; orange, Calgary cohort.

**Figure 2.**
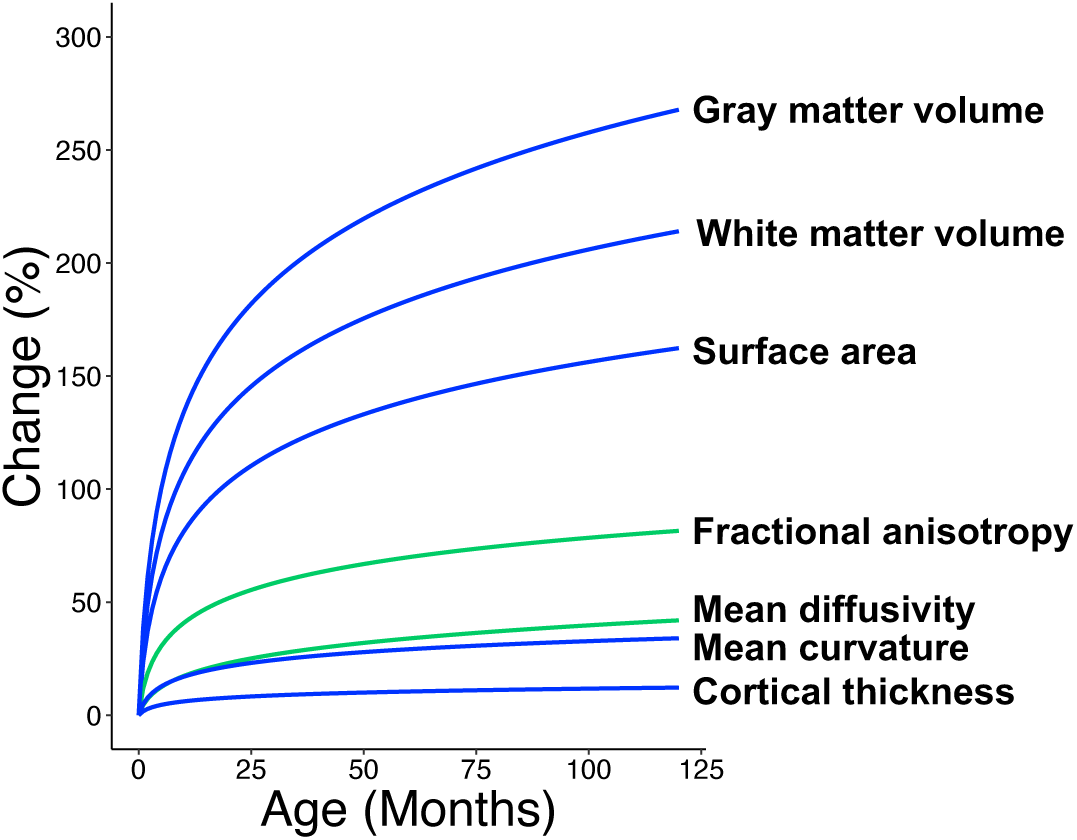
Average longitudinal trajectories from infancy to late childhood by measure. Raw estimates for each brain region examined and each measure were submitted to linear mixed effects models using a logarithmic function. Individual growth curves predicted by this model were averaged to show the overall longitudinal trajectory of the sample for each volumetric/surface-based (blue lines) and diffusion (green lines) measure. Absolute brain estimates were then converted to percent change values to visualize all brain measures along a single axis. For growth curves for separate brain regions and tracts, please see Supplementary Figures 17-22.

### 3.2. Associations between growth curve features and phonological processing

Our second objective was to examine whether brain development affects literacy development. Consequently, curve features (i.e., intercepts and slopes) were extracted from individual-level growth curves and tested for correlations (Pearson) with preschool/early kindergarten phonological processing scores, a subskill critical for literacy (3, 8).

For volumetric and surface-based measures, curve features of gray/white matter volume and surface area in several left hemisphere regions exhibited associations with phonological processing. Specifically, greater phonological processing was associated with (a) greater intercepts of gray matter volume in the left banks of the superior temporal sulcus, (b) greater slopes of white matter volume in left occipitotemporal, temporoparietal, and inferior frontal regions, and (c) greater surface area intercepts and slopes in inferior parietal lobule, pars triangularis, and superior temporal gyrus (Supplementary Figure 23). Next, individual growth curves corresponding to these measures and regions were separated into three groups according to phonological processing scores with low < 85, 85 ≤ average ≤ 115, and high > 115, averaged by group, and then plotted to visualize variation in longitudinal trajectories by phonological processing score. Most notably, children with lower phonological processing scores tended to have less gray matter volume in left banks of the superior temporal sulcus at birth but maintained similar rates of development compared with higher scoring children. At the same time, they had lower rates of white matter volume growth in left occipitotemporal, temporoparietal, and inferior frontal regions (Figure 3).

**Figure 3.**
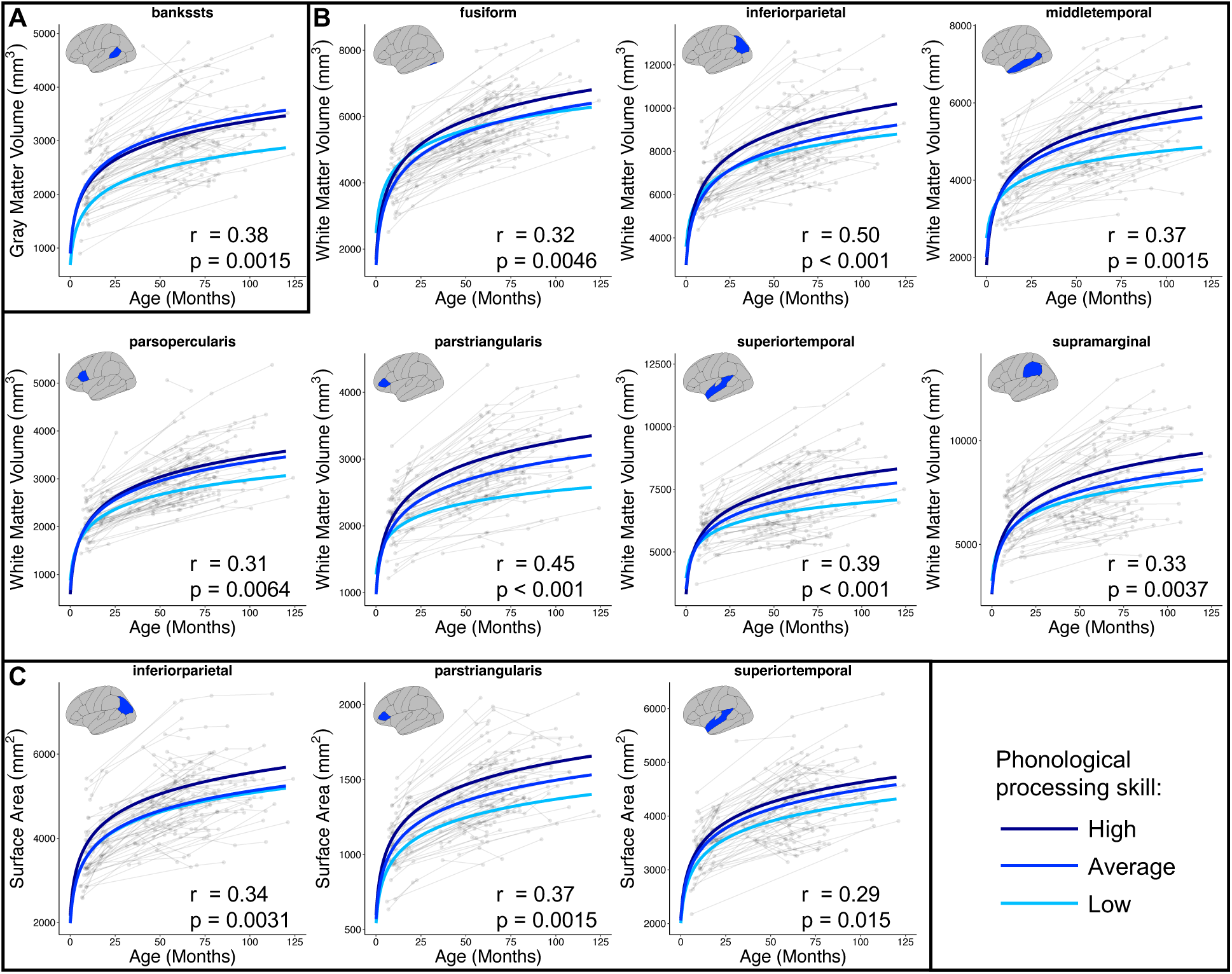
Longitudinal trajectories of brain structure from infancy to late childhood according to phonological processing skill in preschool/early kindergarten. Graphs depict average trajectories for children with low (< 85), average (85–115), and high (> 115) standardized phonological processing scores for measures and regions whose (A) intercepts, (B) slopes, or (C) both intercepts and slopes correlated with phonological processing (p_FDR_ < 0.05). Correlation statistics are reported adjacent to their corresponding plots; intercept and slope statistics for surface area averaged here for visualization purposes but reported separately in Supplemental Figure 23. As a group, children with low phonological processing in preschool/early kindergarten tended to have attenuated longitudinal trajectories, either because they began with lower estimates, as with less gray matter volume in the left banks of the superior temporal gyrus at birth (upper left graph), or because they had slower rates of development, as with white matter volume in other left hemisphere brain regions.

Curve features of white matter organization in the left arcuate fasciculus also correlated with phonological processing outcomes, as greater phonological processing skill was associated with greater slopes of mean diffusivity (Supplementary Figure 24). When distinguishing longitudinal trajectories according to literacy subskills (i.e., low < 85, 85 ≤ average ≤ 115, and high >115), as with volumetric and surface-based measures above, children with low phonological processing scores tended to have higher mean diffusivity at birth but greater rates of mean diffusivity development (i.e., more negative slopes) compared with higher scoring children (Figure 4). No significant associations were observed for the superior or inferior longitudinal fasciculus.

**Figure 4.**
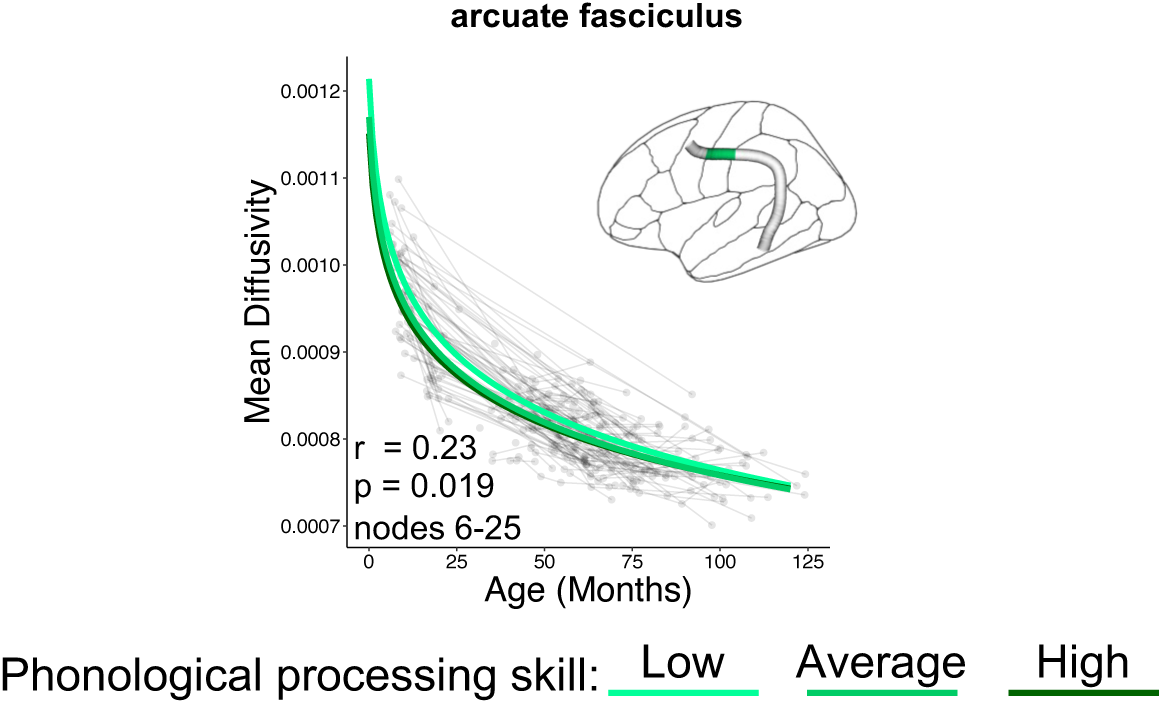
Longitudinal trajectories of mean diffusivity from infancy to late childhood according to phonological processing skill in preschool/early kindergarten. Graph depicts average trajectories for children with low (< 85), average (85–115), and high (> 115) standardized phonological processing scores for the left arcuate fasciculus nodes whose slopes correlated with phonological processing (p_FWE_ < 0.05). Children with low phonological processing tended to exhibit faster development (i.e., more negative slope) in anterior arcuate fasciculus.

### 3.3. Sensitivity Analyses

We performed four sensitivity analyses to test the robustness of the observed brain-behavior relationships. For the first sensitivity analysis, we re-fit our volumetric, surface area, and mean diffusivity estimates with nonlinear mixed effects models using asymptotic functions (Supplementary Figures 25-28). This alternative model to characterize longitudinal trajectories was especially important for mean diffusivity findings, where intercepts and slopes from the main analysis were correlated for many nodes in each tract examined.

We observed that greater phonological processing was still associated with greater intercepts of gray matter volume development in the left banks of the superior temporal sulcus and of surface area in left inferior parietal lobule, pars triangularis, and superior temporal gyrus (Supplementary Table 3A, C). Interestingly, when trajectories were again split according to low (< 85), average (85–115), and high (> 115) standardized phonological processing scores, intercepts were saliently divided, more so than when using linear models (Supplementary Figure 3A,C). Whereas our main analyses allowed us to examine relationships between brain development and literacy subskills directly by using a random slopes term, the nonlinear model used in this sensitivity analysis replaces the random slopes term with a random asymptote term. As such, relationships between brain development and language outcomes may only be inferred where brain measures and regions exhibit asymptote-behavior correlations without corresponding intercept-behavior correlations. Models for middle and superior temporal cortices and supramarginal gyrus white matter volume did not converge. However, other brain regions whose white matter volume and surface area slopes correlated positively with phonological processing in the main analysis also exhibited significant associations between asymptotes and phonological processing. Effect sizes in were generally smaller for white matter volume and larger for surface area compared with the main analysis (Supplementary Tables 3B, C; Supplementary Figure 3B,C). Regarding white matter organization, greater phonological processing was associated with lower mean diffusivity intercepts in left arcuate fasciculus (Supplementary Tables 3D).

Our second sensitivity analysis involved adding phonological processing main and age x phonological processing interaction terms as covariate analogues to brain-behavior correlations with intercepts and slopes, respectively. All results for gray and white matter volume and mean diffusivity present in the main analysis persisted in this sensitivity analysis; however, surface area effects did not (Supplementary Table 4).

For the third sensitivity analysis, we recomputed brain-behavior associations for volumetric and surface-based measures using semipartial correlations, controlling for curve features of the longitudinal trajectory for total intracranial volume (TIV). Relative to the results of the main analysis, effect sizes were reduced when including curve features of TIV. However, associations with phonological processing remained significant for intercepts of the banks of the superior temporal sulcus gray matter and inferior parietal lobule, pars triangularis, and superior temporal gyrus surface area. Slopes of white matter volume development in inferior parietal lobule and pars triangularis also remained significant (Supplementary Table 5).

In the fourth sensitivity analysis, we again recomputed brain-behavior associations, this time controlling for Euler numbers as a reproducible alternative to manual quality control (149). On average, effect sizes showed no drop relative to the main analysis (r_avg_ = 0.35). All brain-behavior associations significant in the main analysis remained significant with the inclusion of Euler numbers, except for the association between phonological processing and slopes of superior temporal gyrus surface area (Supplementary Table 6).

### 3.4. Specificity analyses

We also performed analyses to determine whether brain-behavior associations were limited to specific brain measures, regions, and curve features or contingent upon specific behavioral measures. Whole-brain analyses showed higher brain-behavior effects for white matter volume slopes compared (numerically) to other morphometric measures, especially in left inferior parietal lobule; however, effects did not appear to be specific to the left hemisphere (Supplementary Figure 32). In contrast, brain-behavior associations with mean diffusivity did not replicate in right hemisphere homologue tracts. In a subset of data, we also tested whether associations were specific to phonological processing, to other reading subskills, or to cognitive measures in general. Effects with phonological processing persisted in this subset for all brain-behavior associations except for intercepts of surface area in left inferior parietal and superior temporal cortices. However, similar effects were not observed for oral language skills or nonverbal general cognitive ability (Supplementary Table 7).

### 3.5. Contributions of literacy-related factors to longitudinal trajectories of brain structure and white matter organization

Next, we sought to examine whether the brain-behavior associations, observed in both the full sample and the New England cohort only, are driven by two common literacy-related factors: family history of reading difficulty and the home literacy environment. When modeled as fixed effects, neither constituted significant contributors to the longitudinal trajectories of early brain development that predicted phonological processing (Supplementary Table 8). Furthermore, BIC estimates for models with versus without literacy-related covariates were not significantly different for any brain measures or regions (Supplementary Table 9).

### 3.6. Indirect effects between longitudinal trajectories of early brain development and word reading outcomes via phonological processing

Finally, we examined whether literacy subskills mediated the relationship between brain development curve features (e.g., intercepts and slopes) and decoding and word reading outcomes. As a prerequisite for mediation, the mediator (i.e., phonological processing) must be associated with both the predictor (i.e., brain estimate) and outcome (i.e., word reading). We limited the potential mediator to phonological processing, which related to both decoding (r = 0.48; p < 0.005) and word reading (r = 0.53; p < 0.001), as measured by the Woodcock Reading Mastery Tests III word attack and word identification subtests (114), because brain-behavior associations with oral language were few and inconsistent in sensitivity analyses, and we limited the potential predictors to brain measures and regions/tracts surviving FDR correction in the main analysis. Indirect effects were reported when the 95% confidence intervals, based on 20,000 bootstrapped samples, for the average causal mediation effect did not include 0 (please see Methods).

Phonological processing mediated the relationship between brain and decoding and between brain and word reading for the following measures and regions: intercepts of gray matter volume development in the left banks of the superior temporal sulcus (Figure 5A). Phonological processing skill also mediated associations between decoding/word reading and slopes of white matter volume development and intercepts and slopes of surface area development in left temporo-parietal and inferior frontal regions (Figure 5B, C; Supplementary Tables 10, 11). Indirect effect sizes nominally attenuated when controlling for TIV curve features for decoding (average estimate with TIV = 0.016, average estimate without TIV = 0.020) and word reading (average estimate with TIV = 0.024, average estimate without TIV = 0.031; Supplementary Tables 12, 13). Lastly, phonological processing skill mediated associations between slopes of mean diffusivity development in left arcuate fasciculus and decoding and word reading (Figure 5D).

**Figure 5.**
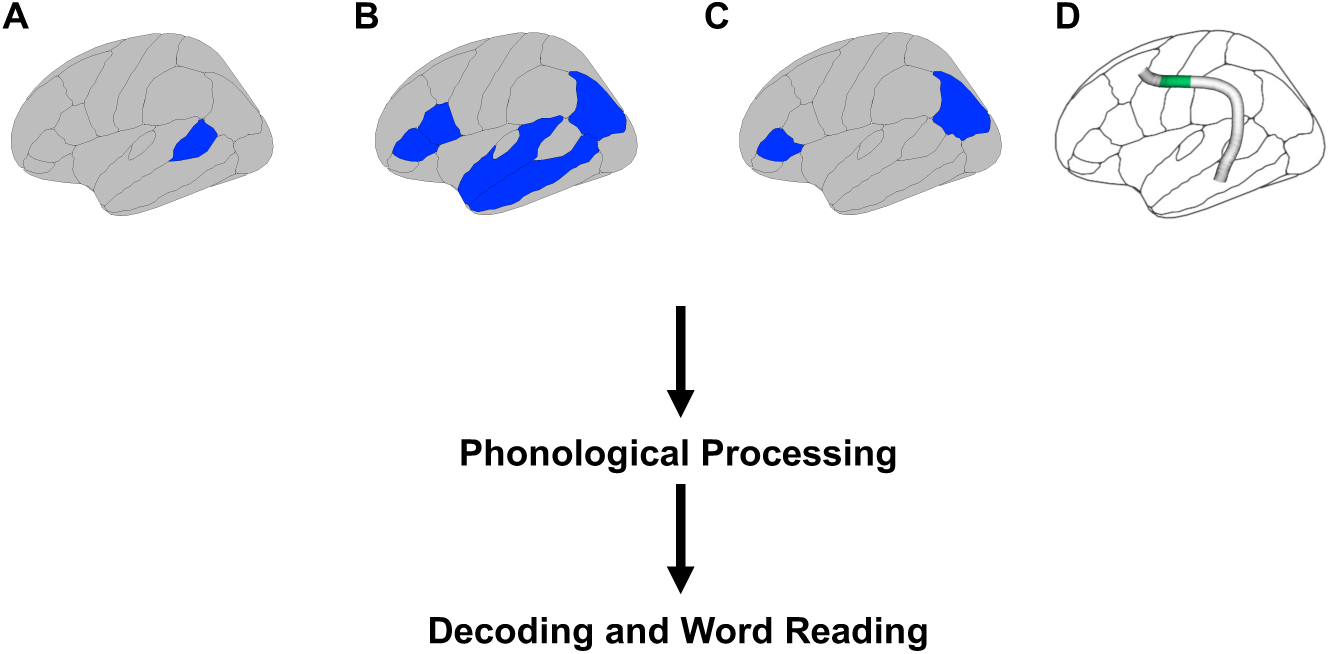
Phonological processing skill mediates the relationship between early brain development and decoding and word reading. Indirect effects (filled arrows) were found for (A) intercepts of gray matter volume in the left banks of the superior temporal sulcus; (B) slopes of white matter volume and (C) intercepts and slopes of surface area in left temporo-parietal and inferior frontal regions; and (D) slopes of mean diffusivity in left arcuate fasciculus (nodes 6-25, green). Note: indirect effects are depicted for surface area slopes only; surface area intercept effects are reported in Supplementary Tables 10, 11.

## 4. Discussion

The brain regions and tracts that eventually support decoding and word reading begin to develop long before the skills themselves emerge. Here, we examined the relationship between longitudinal trajectories of early brain development and acquisition of reading-related (sub)skills in four objectives. First, we generated longitudinal trajectories of early brain structure, including white matter organization, from infancy to school age in regions and tracts previously linked to literacy development (30–32, 34, 50–58) using a novel processing and analysis pipeline appropriate for the early developmental period. Findings showed that longitudinal trajectories were best modeled using logarithmic, compared with linear and quadratic, functions. Second, we examined associations between curve features of longitudinal trajectories and a key literacy subskill: phonological processing. Results showed that curve intercepts (i.e., birth brain estimates) of gray matter volume and surface area and curve slopes (i.e., early brain development) of white matter volume, surface area, and mean diffusivity predicted phonological processing measured in preschool/early kindergarten. While effects were robust in hypothesized left temporo-parietal, occipito-temporal, and inferior frontal regions and tracts, specificity analyses suggest that these brain-behavior associations are not limited to these regions. The predominance and magnitude of slope-outcome associations in comparison with intercept-outcome associations suggests a less stable, more dynamic relationship between brain and literacy development (34). Third, we examined whether familial risk of reading difficulty and home literacy environment, two common literacy-related covariates, influenced those trajectories and found that they did not. Fourth, we expanded the scope of our inquiry to long-term literacy development, showing that phonological processing mediated associations between early brain development and decoding and word reading skills between late kindergarten and second grade. Overall, these findings suggest that the neural foundations for the subsequent development of phonological processing may be partially present at birth but are still forming in the years between birth and preschool and eventually support the development of decoding and word reading skills.

Trajectories of early brain development have been generated from longitudinal studies up to the first two years of life (e.g., 35) and from combined cross-sectional and longitudinal datasets spanning infancy to school age (e.g., 36). However, methodological challenges associated with infant MRI and longitudinal designs in general have created a relative vacuum of longitudinal MRI studies spanning infancy to school age (43). The current study fills this gap as a purely longitudinal examination of early structural brain development in which longitudinal trajectories are generated with a pipeline designed for the early developmental period. Overall, longitudinal trajectories of gray and white matter volume, surface area, fractional anisotropy, and mean diffusivity exhibited rapid postnatal growth that slowed with age, which largely comports with previous early developmental studies using cross-sectional (150–154) and combined longitudinal and cross-sectional designs (13, 36, 46, 48, 155, 156). As shown previously (49, 154), the curves for cortical thickness and mean curvature were generally less steep compared with other brain measures, both initially following birth and into childhood. Nonetheless, empirical comparisons with linear and quadratic functions showed that logarithmic models were more parsimonious for all measures examined.

After generating longitudinal trajectories, we examined how this brain development influenced literacy development by relating curve features of individual trajectories to reading-related subskills measured in preschool/early kindergarten as well as decoding and word reading skills measured between kindergarten and second grade. The finding of direct links to phonological processing and indirect links to decoding and word reading from curve intercepts of gray matter volume (i.e., gray matter volume at birth) specifically in the left banks of the superior temporal sulcus is compelling in the context of a recent large-scale genomics study showing common genetic influences on surface area in this region and reading-related skills (60), including from a gene involved in neurogenesis and axon formation (78). Combined with observed intercept-outcome associations for surface area in other left hemisphere regions and prior work showing an association between reading difficulty and left temporo-parietal sulcal patterns determined in utero (157), these findings offer convergent evidence for a mechanistic pathway through which genetic factors shape the foundations of reading development via prenatal left temporo-parietal brain development. While this intercept-outcome effect remained robust through all four sensitivity analyses, this interpretation should be viewed with caution, as specificity analyses suggest intercept-outcome effects may not be limited to the left banks of the superior temporal sulcus.

The intercept-outcome findings also support the hypothesis that school-age literacy skill builds on an early-developing foundation (33) or ‘neural scaffold’ (30–32). However, the paucity of intercept-outcome associations in contrast to the multitude and magnitude of slope-outcome associations suggests that this neural scaffold develops substantially over the first several years of life. While the mechanisms driving these divergent associations with outcomes will ultimately require further investigation, it is conceivable that they reflect links to distinct early precursors of phonological skills. For instance, the intercept-outcome associations may reflect stable relations with foundational perception skills that develop earlier (e.g., prosody, differentiating phonemes) and remain necessary for the development of more advanced phonological processing skills, whereas slope-outcome associations reflect dynamic associations with more advanced phonological processing skills such as syllable or phoneme deletion or substitution. Overall, the presence of slope-outcome associations offers insights into the early development of a neural scaffold for literacy and in doing so, underscores the importance of examining individual longitudinal trajectories (2), as opposed to cross-sectional and prospective associations.

It was also interesting that most of the slope-outcome findings were with white matter properties, and nearly all of these remained significant in all sensitivity analyses, with some effects becoming even more robust (e.g., please see inferior parietal white matter in second sensitivity analysis). While no prior study has examined the relationship between long-term literacy subskills and longitudinal trajectories of white matter development beginning in infancy (or early development of surface-based measures), there is a small corpus of literature that has linked reading-related skill performance to short-term, school-age developmental changes in left temporo-parietal white matter volume (158) and organization, most consistently in the left arcuate fasciculus (34, 56, 57, 158). This is especially compelling when considering that the slope-outcome relationship we observed with white matter organization was specific to the left arcuate fasciculus and to phonological processing, which is highly predictive of subsequent word reading and decoding performance (4, 8–10). Work from genomics might offer further explanation for the dominance of white matter associations, as genes involved in axon guidance (159), axon formation (78), and oligodendrocyte maturation (160) have also been linked to reading-related skills (60) and reading disability (76, 77). Although white matter volume and organization are not strongly associated and thought to be sensitive to different properties (161), their reliance on myelination, the primary function of oligodendrocytes, and other axonal properties suggests that they may represent another mechanism through which genetic factors may shape early brain development.

Interestingly, specificity analyses showed direct brain-behavior associations with phonological processing but not with oral language skills or nonverbal general cognitive abilities. Prior brain imaging studies in preschoolers/kindergarteners comport with this finding as phonological processing seems to exhibit more consistent associations with brain architecture (15–20), compared with oral language skills (21–23); c.f., (17, 24). While both subskills are considered critical for higher-level reading-related skills, they are thought to function on two distinguishable developmental pathways, whereby phonological processing is more involved in recognizing and decoding printed words and oral language skills are more essential for the development of reading comprehension skills (please see Scarborough’s Reading Rope; (8). It should be noted that this framework also fits with our mediation findings, which show indirect relations between early brain development and decoding and (printed) word reading via phonological processing. Furthermore, whereas phonological processing is limited in its scope, as specifically one’s ability to recognize and manipulate sounds in a word (4–7), oral language was measured in the current study as a composite of vocabulary and oral comprehension subtests, with the latter requiring children to engage with semantic and syntactic cues. As different brain regions may be more specialized for certain component language skills (e.g., pars opercularis may be more involved in syntactic processing while pars triangularis and middle temporal gyrus may be more involved in semantics; (162, 163)), aggregating distinct oral language skills may have obscured brain-behavior associations. Conversely, it is also conceivable that oral language acquisition requires additional, moderating factors that were not modeled in the present study (e.g., social interactions (164) or conversational turn-taking (21)). Nevertheless, the necessity of phonological processing for literacy development and its relation to early brain development reported here underscore the importance of examining reading-related subskills at the very beginnings of life.

Turning to literacy-related covariates, familial history of reading difficulty (FHD) was hypothesized to influence longitudinal trajectories of early brain development, based on previous findings showing FHD-related alterations in brain architecture in infancy (71, 72) and preschool (16, 18, 57). Consequently, the observation that FHD status did not influence longitudinal trajectories in any regions or tracts examined was initially unexpected. However, it is important to recognize that a child who has an older sibling or parent with a reading difficulty (FHD+) does not necessarily have a genetic susceptibility for reading difficulty (165), nor will they necessarily develop reading challenges given the multifactorial nature of reading difficulties. Rather, FHD status should be considered one of multiple risk factors that can contribute to long-term literacy development (166, 167), either through intergenerational transmission of genes or environment (168). Consistent with this, roughly half of FHD+ children develop typical reading skills (62, 63), and those who do develop typical reading skills have, as a group, been shown to recruit right- and inter-hemispheric compensatory pathways (169, 170), a pattern similar to children with reading difficulty who subsequently show improvements (171). Although the specificity analysis examining brain-behavior effects in non-a-priori brain regions did not point to literacy-related effects solely in the left hemisphere, right hemisphere regions and tracts were not thoroughly assayed in the current study. Therefore, it may behoove future studies with larger sample sizes to examine the development of right hemisphere regions and tracts in the context of FHD status and to distinguish FHD+ children who develop typical reading skills from FHD+ children who do not.

The home literacy environment also did not significantly influence longitudinal trajectories in the regions and tracts examined, despite reported links to brain structure and function in infants (95, 96) and preschoolers/kindergarteners (21, 24, 91–94). However, with few exceptions (e.g., common associations to arcuate fasciculus fractional anisotropy in infants (96) and kindergarten (24)), specific brain measures and regions/tracts linked to home literacy environment variables identified at the infant time point were not also identified at the preschool/kindergarten time point, suggesting that brain-home literacy environment relations may vary across this developmental window. Future longitudinal studies with more frequent sampling of observations will be needed to test whether this explanation is accurate. Also, as examination of home literacy was limited to longitudinal trajectories for brain measures and regions/tracts that were associated with phonological processing, it is likely that measures and regions/tracts related to the home literacy environment went untested in the current study.

This study had five main limitations. First, consistent with prior work (36), the models we fit to early brain development generated smooth longitudinal trajectories. In actuality, it is unlikely that early brain development transpires as predictably, especially during sensitive and critical periods (172) or specific learning milestones (2). Although participants in the current study were sampled over three times on average, which is preferred for modeling growth curves (173) and more than most longitudinal imaging studies (174), future studies would benefit from increased sampling, particularly around learning milestones germane to literacy development. Second, the sample size in the current study was relatively small when compared with the sample sizes used in multi-site, combined cross-sectional and longitudinal studies (e.g., 13, 36). While sensitivity analyses for the most part demonstrated the robustness of the results, future studies with larger sample sizes will be needed to confirm the findings presented here. Third, cortical thickness, mean curvature, fractional anisotropy, and mean diffusivity exhibited high correlations between random parameters (i.e., intercept-slope correlations) when modeled with logarithmic functions, which spurred concerns over the accuracy of the estimated random parameters. For the current findings, inaccurate random parameters could have generated false negatives for the former three measures and false positives for mean diffusivity. Consequently, for the former three parameters, even though they were consistently better fit with logarithmic functions, we re-analyzed intercept-outcome and slope-outcome associations using random parameters from models with quadratic functions, which had considerably lower intercept-slope correlations. Despite this, results remained non-significant. Meanwhile, the first sensitivity analysis addressed concerns for mean diffusivity by showing that nonlinear mixed effects models using asymptotic functions decoupled intercepts and slopes while maintaining the significant results observed in the main analysis. It is also important to note that this limitation does not apply to findings for gray and white matter volume or surface area. Fourth, we examined somewhat narrowly literacy-related factors that could contribute to early brain development by only including FHD status and home literacy environment, and it is likely that factors not included also contribute to the longitudinal trajectories examined (e.g., teaching quality, educational opportunities, executive functioning skills (166, 175). Fifth, our longitudinal analysis pipeline identifies and removes brain estimates preceding or following developmental changes that are too steep to occur neuroanatomically and more likely to emerge from region or tract mislabeling, despite our quality control efforts. Consequently, it is possible that estimates without steep developmental changes also suffer mislabeling but remain undetected. Overall, interpretations of findings should be considered in the context of these limitations.

In conclusion, this study examined associations between longitudinal trajectories of early brain development beginning in infancy and long-term reading acquisition, specifically literacy subskills. Longitudinal trajectories were generated using a novel, reproducible pipeline we designed specifically for examining early brain development and included familial risk of a reading difficulty and environmental covariates. Findings indicate that preschool/early kindergarten phonological processing, one of the strongest predictors of subsequent word reading development, relates to gray matter volume and surface area at birth and development of white matter volume, surface area, and mean diffusivity across early development. These results offer further evidence for a neural scaffold for literacy development, which is present at birth and continues forming across the first several years of life. The present study also provides a roadmap for future longitudinal studies to examine the relationship between early brain development and acquisition of other academic skills. Understanding when the foundations for reading emerge can deliver important insights into the development of instructional approaches and preventative, and intervention strategies.

## Acknowledgements

This work was funded by NIH–NICHD R01 HD065762 (2016–2021), the William Hearst Fund (Harvard University), and the Harvard Catalyst/NIH (5UL1RR025758) to N.G. We would like to thank all participating families for their long-term dedication to this study. We are grateful for all additional members of the research team who contributed to data acquisition and quality control, including Victoria Hue, Ja’Kala Barber, Emily Hu, Olivia Baldi, Brooke Jordan, Jade Dunstan, Clarisa Carruthers, Carolyn King, Doroteja Rubez, Kathryn Garrisi, Ally Lee, Omeed Moini, Anaïs Colin, Zumin Chen, Colby Weiss, and Olivia Oh. We also thank Rebecca Petersen for her input on psychometric assessments, Jennifer Zuk for work on infant scanning procedures, Borjan Gagoski and Ross Mair for efforts on scanning sequences, Luke Miratrix and Fernando Aguate for guidance on (non)linear mixed effects models, and Banu Ahtam for feedback.

## Supplementary Methods

### 1. Participants

#### New England cohort

Children participated in a longitudinal investigation of brain and literacy development from infancy to school age (NIH–NICHD R01 HD065762). Families of these children were recruited from the New England Area using the Research Participant Registry provided by the Division of Developmental Medicine at Boston Children’s Hospital, and with flyers and ads disseminated in local schools and newspapers, at community events, and on social media. Children were enrolled as infants with the expectation that participation would continue at subsequent developmental stages across the first decade of life. Children were excluded from the study if at any timepoint they were diagnosed with neurological, sensory, or motor disorders or had contraindications for MRI evaluation (e.g., metal implants). All children included were from English-speaking families and were born at gestational week 37 or later. Race demographics, whose reporting is encouraged to improve equitability in neuroscience (1), are as follows: 70% White/Caucasian, 6% Black/African American, 5% Asian, 16% Multiracial, and 5% Hispanic. Neuroimaging and behavioral testing were conducted at Boston Children’s Hospital prior to 2021 and at the Center for Brain Science at Harvard University from 2021 onward. At Boston Children’s Hospital, children’s anatomical MRI scans (please see parameters below) did not show any potentially malignant brain features, as reviewed by a pediatric neuroradiologist. This study was approved by the Institutional Review Boards of Boston Children’s Hospital (IRB-P00023182) and Harvard University (IRB21-0916). Informed written consent was provided by each participating infant’s parent(s) and children gave verbal assent for participation after 50 months of age.

#### Calgary cohort

Children in the Calgary, Alberta area were recruited from the ongoing study on pregnancy outcomes and nutrition (2). Children in this study were predominantly (roughly 90%) from Caucasian families, but also included Asian/Pacific Islander, African, Filipino, Latino/Hispanic, and Multiracial children. Children were born at gestational week 36 or later and did not have diagnosed genetic, neurological, or neurodevelopmental disorders. The study was approved by the University of Calgary Conjoint Health Research Ethics Board (REB13-0020). Parent and/or guardian consent and child assent were acquired for all participants. For additional information, please see the open-source repository (https://osf.io/axz5r/) and previous publications (3–9).

### 2. Further description for the phonological processing composite in the New England cohort

The phonological processing composite comprises three subtests: word access, word fluency, and substitution. For the word access subtest, the child is asked to provide a word that has a specific phonemic element in a specific location. For example, the child may be shown an image, told what it is, and then shown three additional images and asked to select the one that begins with the same sound as the first image. For the word fluency subtest, the child is asked to name as many words as possible that begin with a specific sound within a 1-minute time frame. For example, the child may be asked to name words that begin with the /b/ sound. Lastly, for the substitution subtest, the child is asked to substitute part of one word to create a new word. For instance, the child may be asked to say the resulting word when they replace the /b/ sound in “bunny” with the /s/ sound. Overall, these assessments are designed to probe phonological and phonemic processing/awareness.

### 3. MRI data acquisition

#### New England cohort

All data were acquired on a 3.0 T Siemens scanner with a 32-channel head coil. Please note, sequence parameters varied to optimize acquisition for the increasing head size and neuroanatomy of the participants across the developmental window. Structural T1-weighted magnetization-prepared rapid gradient-echo (MPRAGE) scans were acquired with the following parameters: TR = 2270-2520 ms, TE = 1.66-1.73 ms, field of view = 192-224 mm, 1 mm^3^ voxels, 144-176 sagittal slices. Diffusion echo planar images were acquired using the following parameters: TR = 3800-8320 ms, TE = 88-89 ms, flip angle = 90°, field of view = 180-256 mm, voxel size = 2 x 2 x 2 mm^3^, 62-78 slices, 30 b = 1000 s/mm^2^ gradient directions, 10-11 b = 0 s/mm^2^ non-diffusion-weighted volumes. Diffusion data were acquired with slice-acceleration (SMS/MB) factor = 2 and one reverse phase encoding (i.e., posterior-to-anterior) volume. Please note, sequence parameters varied to optimize acquisition for the increasing head size and neuroanatomy of the participants across the developmental window.

#### Calgary cohort

All data were acquired on a 3.0 T General Electric scanner with a 32-channel head coil. Structural T1-weighted scans were acquired with the following parameters: TR = 8.23 ms, TE = 3.76 ms, field of view = 230 mm, 0.45 x 0.45 x 0.9 mm^3^ voxels, 210 slices. Diffusion echo planar images were acquired using the following parameters: TR = 6750 ms, TE = 79 ms, flip angle = 90°, field of view = 200 mm, voxel size = 0.78 x 0.78 x 2.2 mm^3^, 50-55 slices, 30 b = 750 s/mm^2^ gradient directions, 5 b = 0 s/mm^2^ non-diffusion-weighted volumes. Diffusion acquisition did not use a slice-acceleration factor and did not include a reverse phase encoding (i.e., posterior-to-anterior) volume.

### 4. Algorithm for combining FreeSurfer and iBEATv2.0 segmentations

Segmentations from each software package were then hybridized using in-house MATLAB code that combined the cortical pial and white matter boundaries labeled by iBEATv2.0 with the FreeSurfer subcortical parcellations (i.e., effectively relabeling cortical gray and white matter in the FreeSurfer segmentation using iBEATv2.0 tissue-class demarcations). Herein, iBEATv2.0 3-class tissue segmentations were first resampled to the space of the aseg file. iBEATv2.0 gray matter and cerebrospinal fluid voxels that overlapped with the aseg gray matter and cerebrospinal fluid labels, respectively, inherited the latter’s labels. iBEATv2.0 white matter voxels overlapping with aseg white *or gray* matter received the aseg white matter label; as iBEATv2.0 does not distinguish hemispheres in its labels and FreeSurfer generally overestimates gray matter in this age group, gray matter information was also used here for relabeling. iBEATv2.0 voxels unlabeled by these procedures were then submitted to a nearest neighbor search to identify aseg labels at minimum Euclidean distance. As FreeSurfer generally performed better compared with iBEATv2.0 on subcortical areas, all subcortical (non-cerebellum areas) were relabeled with subcortical FreeSurfer labels. Finally, hybridized segmentation files were converted to FreeSurfer-style white matter files and submitted to a modified version of the standard FreeSurfer v7.3 “recon” pipeline.

### 5. Preparation of Euler numbers

Similar to Bethlehem and colleagues 2022, for further quality control, we also extracted Euler numbers (10), which quantify topological defects in FreeSurfer’s cortical reconstruction and have been shown to correlate with visual ratings and mark artifactual images (11). Euler numbers from each hemisphere were averaged across observations for each participant (average: (11); sum: (12)) and used in sensitivity analyses.

### 6. Implementations of pyAFQ and pyBabyAFQ

Whole-brain tractography was then submitted for fiber tract segmentation to the open-source instantiation of Automated Fiber Quantification (AFQ; (13, 14). Herein, waypoint regions-of-interest (ROIs) and a probabilistic fiber atlas were mapped from a standard template to the individual brain. Fibers were delineated into separate tracts according to the waypoint ROIs through which they passed and the tract they were most likely to belong based on the probability atlas. Additional fibers were removed (i.e., outlier detection) if they deviated sufficiently from the core of the tract. While all diffusion data underwent the above processes, parameters differed by age group. For brains > 25 months, the standard pyAFQ pipeline was used, in which ROIs and the probability fiber atlas were mapped from an adult MNI template and fibers were removed if they were more than five standard deviations from the core of the tract (14). In contrast, for brains ≤ 24 months, we used pyBabyAFQ (15), an implementation in the pyAFQ suite designed to accommodate the smaller neuroanatomy in infants. Accordingly, ROIs and the probabilistic fiber atlas are mapped from an infant template (16), the ROIs are smaller compared to those used by standard pyAFQ, an additional ROI was used for tracts with acute curves, and the fiber outlier detection threshold was reduced to four standard deviations from the tract core. Importantly, varying the parameters used to segment the tracts by age group preserves the accuracy with which tracts are segmented and reduces age-related bias that would emerge if using standard (i.e., suboptimal) parameters for younger children.

### 7. Quality control procedures for longitudinal trajectory estimation

Prior to modeling, estimates of brain structure and white matter organization, excepting cortical thickness and mean curvature, underwent a final quality control procedure to remove neuroanatomically implausible observations by setting annual change thresholds. For instance, it was unlikely that gray matter volume in any particular brain area changed by more than 10% per year in children over 25 months. Accordingly, for brains > 25 months, observations for brain regions and tracts of interest (please see above) were flagged if they were preceded or followed by ≥ 10% annual change (positive or negative). To improve model convergence in the first sensitivity analysis (with nonlinear mixed effects models), the threshold for mean diffusivity was lowered from 10% to 5%. To mitigate data loss, we next identified which timepoint—the earlier timepoint or later timepoint—was more likely to be inaccurate, using an outlier detection procedure. Herein, we generated average (across participants) estimates for each brain area/tract and each brain measure by timepoint (roughly, grade level) to which to compare the observations flagged in the previous step. Out of the pair of flagged observations, the one farther from the average estimate corresponding to its timepoint was discarded and the other flagged observation was preserved. This process was done iteratively to handle cases in which children over 50 months had MRI observations from more than two timepoints. A similar quality control procedure was used for brains ≤ 24 months, but it needed to account for the rapid brain growth already thoroughly reported (17, 18). Consequently, observations were only flagged when inter-timepoint changes were negative for gray/white matter volume, surface area, or fractional anisotropy, or positive for mean diffusivity. No inter-timepoint threshold was used for cortical thickness or mean curvature for brains ≤ 24 months. All remaining participants after these additional quality control procedures had multiple observations (i.e., longitudinal datasets), including one from ≤ 24 months.

### 8. Sensitivity analyses

To test the reliability of our results, we performed a replication analysis on gray/white matter volume, and surface area with nonlinear mixed effects models using asymptotic functions from the R ‘nlme’ package (19), similar to that described in Alex and colleagues (18); https://github.com/knickmeyer-lab/ORIGINs_ICV-and-Subcortical-volume-development-in-early-childhood). Intercepts and asymptotes were modeled as fixed and random effects; rate constants were modeled as fixed effects. The reason that this model was not used in the main analysis is that it does not use a random slopes term, and a key aspect of the current study is to examine the relation between brain growth and literacy subskills. However, it should be noted that the nonlinear model does provide an indirect examination of the relation between growth and literacy subskills; e.g., if there is no association between the intercept and the literacy subskill, but there is an association between the asymptote and the literacy subskill, then it could be inferred that there is an association between brain growth and the literacy subskill. Although both linear and nonlinear models have different requirements (e.g., linear models require relationships between predictors and outcomes to be linear), the functional forms used on the current dataset in both cases fit the data closely (Supplementary Figure 8), suggesting random parameters were similar or proportional across models.

For the main analysis, we opted to model brain development prior to testing brain-behavior associations because this comports with the temporal order theorized, that brain development effects subsequent behavioral skills. Also, our sample size is larger for longitudinal brain data alone compared with longitudinal brain data plus reading-related outcomes; therefore, longitudinal models of brain development would be improved if not including outcomes in the model. However, in practice, contributions of phonological processing main and age x phonological interaction terms should be analogous to associations between phonological processing and curve intercepts and slopes, respectively. Consequently, we also examined phonological processing main and age x phonological processing interaction terms included in the linear mixed effects models.

In addition, we did not initially control for total intracranial volume (TIV) when modeling longitudinal trajectories of brain structure, consistent with other work examining developmental trajectories of brain structure beginning in infancy (12, 18, 20–23). Further, recent work has shown that TIV correction may be problematic, reducing brain-behavior predictive accuracies for gray matter volume and surface area or potentially generating spurious predictions for cortical thickness (24). Given these concerns and recommendations to report both raw and TIV-corrected results (25), we thought that an appropriate use of TIV would be to recompute brain-behavior associations for volumetric and surface-based measures using semipartial correlations (Pearson) with the random terms from longitudinal modeling with TIV as covariates of no interest. We report results of the linear mixed model run on brain-behavior relations with TIV.

Lastly, for volumetric and surface-based measures, visual ratings of cortical surfaces were used to identify sub-optimal datasets (visual ratings of tract reconstructions were for measures of white matter organization). However, Euler numbers, which quantify the number the topological defects in FreeSurfer’s cortical surface reconstruction (10), have been shown to consistently correlate with quality ratings (11) and to serve as reproducible alternatives to manual quality control, including manual editing (26). Comparable to Rosen and colleagues, visual ratings and Euler numbers were highly correlated after controlling for age and biological sex (r = 0.44, p < 0.001). Therefore, to account for residual variance due to segmentation quality in a data-driven, reproducible manner, we submitted volumetric and surface-based brain-behavior associations to semipartial correlations (Pearson) with average (across timepoint) Euler numbers.

## Supplemental Tables

**Supplementary Table 1.** Participant demographics.

| Supplementary Table 1. Participant demographics |  |  |  |
| --- | --- | --- | --- |
|  |  | Structure | Diffusion |
| *General information* | N. participants | 98 | 128 |
|  | N. observations | 276 | 396 |
|  | N. observations per participant | 3 ± 1 | 3 ± 1 |
|  | Age at literacy-related subskill testing (months) | 63 ± 5.3 | 63 ± 5.1 |
|  | Age at decoding/word reading testing (months) | 82 ± 6.4 | 82 ± 6.4 |
| *Covariates* | Biological sex (F/M) | 46/52 | 67/61 |
|  | Maternal education (years) | 17 ± 2.1 | 17 ± 2.1 |
|  | Cohort ([New England]/Calgary) | 59/39 | 77/51 |
|  | Family history of reading difficulty (+/-) | 30/29 | 39/38 |
|  | Home literacy environment (a.u.) | 0.073 ± 0.40 | 0.041 ± 0.42 |
| *Literacy-related subskills* | Phonological processing standard score | 108 ± 13 | 106 ± 14 |
|  | Oral language standard score | 115 ± 13 | 113 ± 13 |
| *Decoding/ word reading* | Word attack standard score | 115 ± 13 | 112 ± 14 |
|  | Word identification standard score | 113 ± 17 | 109 ± 17 |
| *Cognitive abilities* | Nonverbal general cognitive ability | 109 ± 12 | 106 ± 13 |

**Supplementary Table 2.**
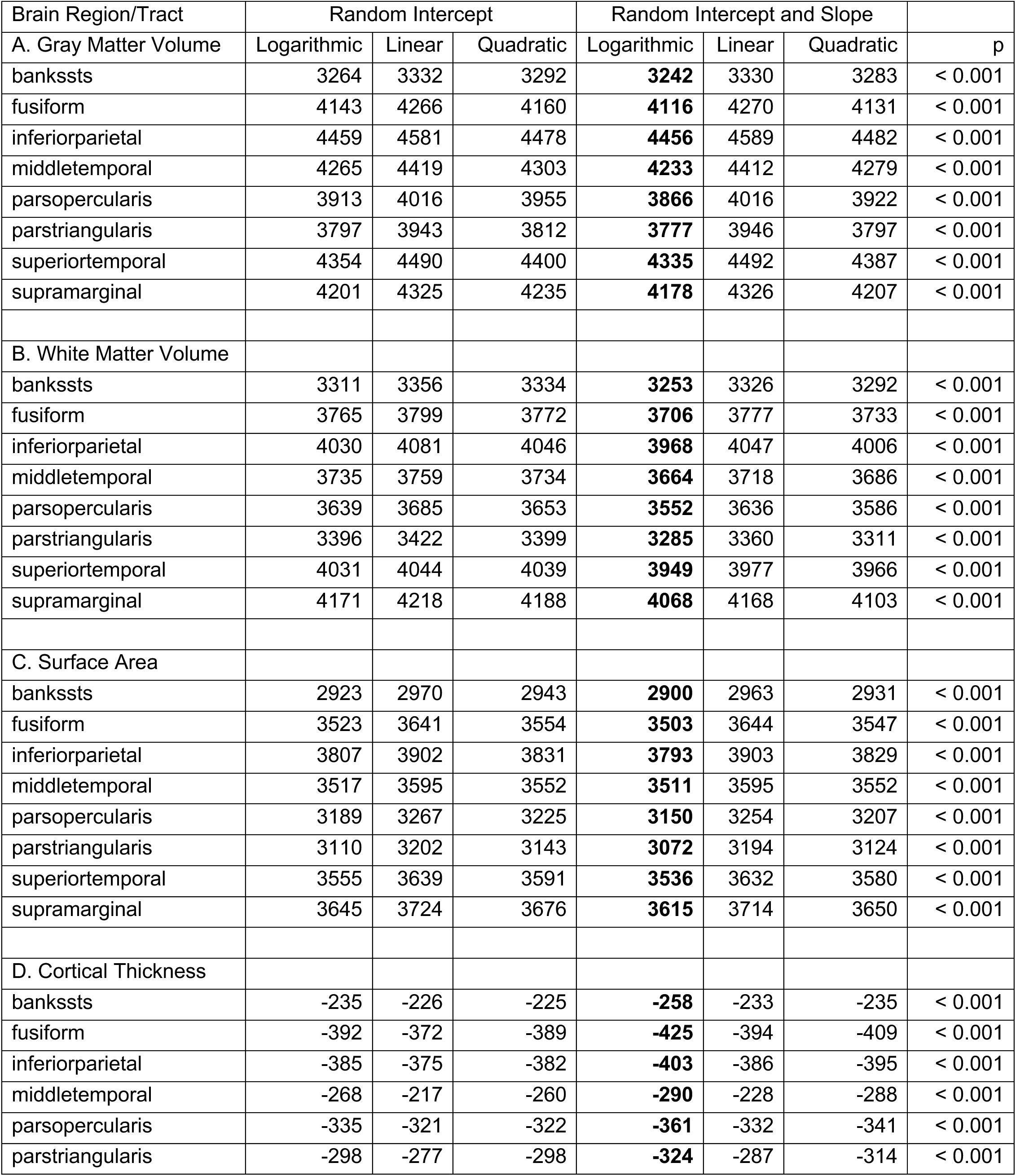
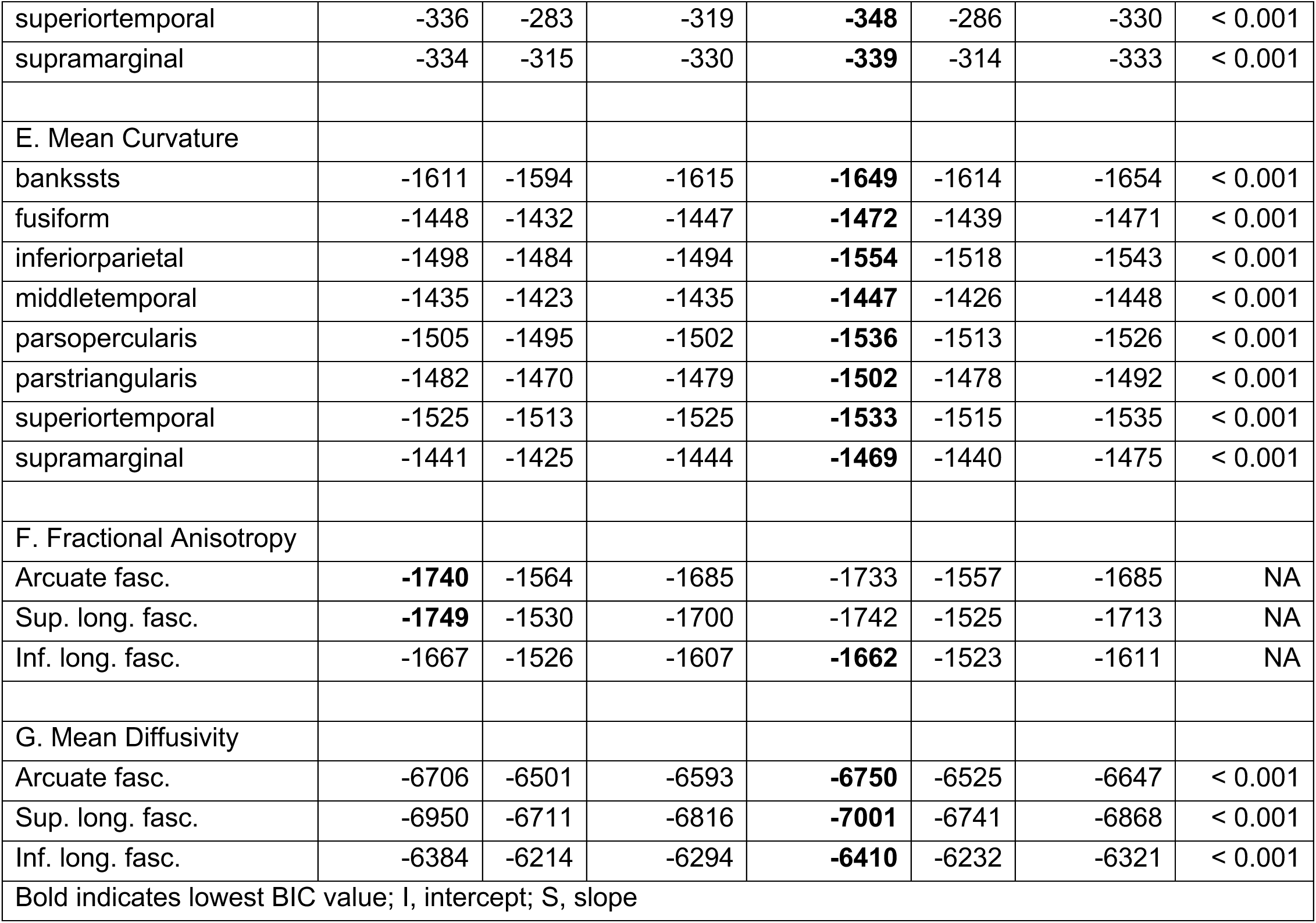
Comparison of Linear Mixed Effects Models for Gray Matter Volume.

**Supplementary Table 3.** Brain-behavior associations with nonlinear mixed effects model.

| Brain Region/Tract | Intercept |  |  | Asymptote |  |  |
| --- | --- | --- | --- | --- | --- | --- |
| A. Gray Matter Volume | r | p | pFDR | r | p | pFDR |
| bankssts | 0.32 | 0.0088 | 0.034 | 0.28 | 0.020 | 0.053 |
| fusiform | 0.02 | 0.86 | 0.86 | 0.08 | 0.48 | 0.48 |
| inferiorparietal | 0.28 | 0.013 | 0.034 | 0.28 | 0.013 | 0.051 |
| middletemporal | 0.20 | 0.082 | 0.11 | 0.20 | 0.082 | 0.11 |
| parsopercularis | 0.17 | 0.14 | 0.16 | 0.22 | 0.059 | 0.11 |
| parstriangularis | 0.29 | 0.0094 | 0.034 | 0.32 | 0.0051 | 0.040 |
| superiortemporal | 0.22 | 0.050 | 0.080 | 0.19 | 0.096 | 0.11 |
| supramarginal | 0.28 | 0.017 | 0.034 | 0.20 | 0.095 | 0.11 |
| B. White Matter Volume |  |  |  |  |  |  |
| bankssts | 0.26 | 0.026 | 0.066 | 0.26 | 0.027 | 0.027 |
| fusiform | 0.11 | 0.34 | 0.34 | 0.30 | 0.0080 | 0.013 |
| inferiorparietal | 0.30 | 0.010 | 0.050 | 0.43 | < 0.001 | < 0.001 |
| middletemporal | NA | NA | NA | NA | NA | NA |
| parsopercularis | 0.12 | 0.30 | 0.34 | 0.29 | 0.011 | 0.014 |
| parstriangularis | 0.23 | 0.048 | 0.080 | 0.40 | < 0.001 | 0.0012 |
| superiortemporal | NA | NA | NA | NA | NA | NA |
| supramarginal | NA | NA | NA | NA | NA | NA |
| C. Surface Area |  |  |  |  |  |  |
| bankssts | 0.30 | 0.014 | 0.027 | 0.29 | 0.015 | 0.029 |
| fusiform | 0.14 | 0.24 | 0.24 | 0.20 | 0.084 | 0.096 |
| inferiorparietal | 0.35 | 0.0018 | 0.0072 | 0.39 | < 0.001 | 0.0023 |
| middletemporal | 0.22 | 0.062 | 0.083 | 0.23 | 0.050 | 0.081 |
| parsopercularis | 0.21 | 0.078 | 0.089 | 0.21 | 0.072 | 0.096 |
| parstriangularis | 0.39 | < 0.001 | 0.0046 | 0.39 | < 0.001 | 0.0023 |
| superiortemporal | 0.30 | 0.011 | 0.027 | 0.30 | 0.010 | 0.027 |
| supramarginal | 0.26 | 0.030 | 0.049 | 0.20 | 0.097 | 0.097 |
| D. Mean Diffusivity |  |  |  |  |  |  |
| Arcuate fasc.* | -0.22 | 0.033 | NA | 0.03 | 0.65 | NA |
| *averaged across p<sub>FWE</sub> < 0.05 significant nodes |  |  |  |  |  |  |

**Supplementary Table 4.** Contributions of outcomes to linear mixed effects model.

| Brain Region/Tract | Main Effect |  |  | Interaction |  |  |
| --- | --- | --- | --- | --- | --- | --- |
| A. Gray Matter Volume | Effect size | p | pFDR | Effect size | p | pFDR |
| bankssts | $1.8 \times 10^{+01}$ | < 0.001 | 0.0044 | $-1.1 \times 10^{+00}$ | 0.52 | 0.90 |
| fusiform | $1.1 \times 10^{+01}$ | 0.52 | 0.52 | $-7.4 \times 10^{-01}$ | 0.87 | 0.90 |
| inferiorparietal | $4.0 \times 10^{+01}$ | 0.073 | 0.14 | $3.1 \times 10^{+00}$ | 0.58 | 0.90 |
| middletemporal | $2.9 \times 10^{+01}$ | 0.066 | 0.14 | $-6.7 \times 10^{-01}$ | 0.90 | 0.90 |
| parsopercularis | $1.7 \times 10^{+01}$ | 0.14 | 0.19 | $-7.1 \times 10^{-01}$ | 0.82 | 0.90 |
| parstriangularis | $8.3 \times 10^{+00}$ | 0.21 | 0.24 | $2.4 \times 10^{+00}$ | 0.18 | 0.90 |
| superiortemporal | $3.4 \times 10^{+01}$ | 0.087 | 0.14 | $-3.2 \times 10^{+00}$ | 0.62 | 0.90 |
| supramarginal | $4.5 \times 10^{+01}$ | 0.014 | 0.055 | $-2.3 \times 10^{+00}$ | 0.70 | 0.90 |
| B. White Matter Volume |  |  |  |  |  |  |
| bankssts | $3.3 \times 10^{+00}$ | 0.33 | 0.37 | $1.9 \times 10^{+00}$ | 0.18 | 0.18 |
| fusiform | $-1.0 \times 10^{+01}$ | 0.35 | 0.37 | $7.3 \times 10^{+00}$ | 0.015 | 0.017 |
| inferiorparietal | $-2.9 \times 10^{+01}$ | 0.027 | 0.13 | $2.0 \times 10^{+01}$ | < 0.001 | < 0.001 |
| middletemporal | $-1.8 \times 10^{+01}$ | 0.092 | 0.17 | $9.9 \times 10^{+00}$ | 0.0017 | 0.0035 |
| parsopercularis | $-9.8 \times 10^{+00}$ | 0.050 | 0.13 | $5.0 \times 10^{+00}$ | 0.0054 | 0.0072 |
| parstriangularis | $-7.2 \times 10^{+00}$ | 0.11 | 0.17 | $5.5 \times 10^{+00}$ | < 0.001 | < 0.001 |
| superiortemporal | $-2.3 \times 10^{+01}$ | 0.033 | 0.13 | $1.3 \times 10^{+01}$ | < 0.001 | < 0.001 |
| supramarginal | $-1.1 \times 10^{+01}$ | 0.37 | 0.37 | $1.2 \times 10^{+01}$ | 0.0049 | 0.0072 |
| C. Surface Area |  |  |  |  |  |  |
| bankssts | $4.9 \times 10^{+00}$ | 0.0078 | 0.062 | $-2.1 \times 10^{-02}$ | 0.97 | 0.97 |
| fusiform | $1.4 \times 10^{+00}$ | 0.73 | 0.73 | $1.0 \times 10^{+00}$ | 0.35 | 0.56 |
| inferiorparietal | $9.4 \times 10^{+00}$ | 0.21 | 0.42 | $3.2 \times 10^{+00}$ | 0.11 | 0.43 |
| middletemporal | $3.0 \times 10^{+00}$ | 0.55 | 0.62 | $1.4 \times 10^{+00}$ | 0.29 | 0.56 |
| parsopercularis | $1.6 \times 10^{+00}$ | 0.45 | 0.60 | $4.8 \times 10^{-01}$ | 0.53 | 0.70 |
| parstriangularis | $1.9 \times 10^{+00}$ | 0.20 | 0.42 | $1.1 \times 10^{+00}$ | 0.026 | 0.21 |
| superiortemporal | $4.5 \times 10^{+00}$ | 0.35 | 0.56 | $1.8 \times 10^{+00}$ | 0.22 | 0.56 |
| supramarginal | $1.4 \times 10^{+01}$ | 0.019 | 0.076 | $-7.2 \times 10^{-01}$ | 0.71 | 0.81 |
| D. Mean Diffusivity |  |  |  |  |  |  |
| Arcuate fasc.* | $-2.4 \times 10^{-06}$ | 0.0092 | NA | < 0.001 | 0.0069 | NA |
| *averaged across p<sub>FWE</sub> < 0.05 significant nodes |  |  |  |  |  |  |

**Supplementary Table 5.** Brain-behavior associations controlling for TIV.

|  |  |  |  |  |
| --- | --- | --- | --- | --- |
| Brain Region | Measure-Curve Feature | r | p | pFDR |
| bankssts | GMV-intercept | 0.37 | 0.0021 | 0.017 |
| fusiform | WMV-slope | 0.21 | 0.068 | 0.11 |
| inferiorparietal | WMV-slope | 0.41 | < 0.001 | 0.0022 |
| middletemporal | WMV-slope | 0.23 | 0.050 | 0.10 |
| parsopercularis | WMV-slope | 0.19 | 0.11 | 0.14 |
| parstriangularis | WMV-slope | 0.35 | 0.0030 | 0.012 |
| superiortemporal | WMV-slope | 0.23 | 0.046 | 0.10 |
| supramarginal | WMV-slope | 0.15 | 0.19 | 0.22 |
| inferiorparietal | SA-intercept | 0.31 | 0.0067 | 0.027 |
| parstriangularis | SA-intercept | 0.35 | 0.0024 | 0.019 |
| superiortemporal | SA-intercept | 0.26 | 0.029 | 0.075 |
| inferiorparietal | SA-slope | 0.20 | 0.092 | 0.37 |
| parstriangularis | SA-slope | 0.24 | 0.043 | 0.34 |
| superiortemporal | SA-slope | 0.01 | 0.96 | 0.99 |
| TIV, total intracranial volume |  |  |  |  |

**Supplementary Table 6.** Brain-behavior associations controlling for Euler numbers.

|  |  |  |  |  |
| --- | --- | --- | --- | --- |
| Brain Region | Measure-Curve Feature | r | p | pFDR |
| bankssts | GMV-intercept | 0.38 | 0.0013 | 0.011 |
| fusiform | WMV-slope | 0.33 | 0.0037 | 0.0049 |
| inferiorparietal | WMV-slope | 0.50 | < 0.001 | < 0.001 |
| middletemporal | WMV-slope | 0.36 | 0.0016 | 0.0031 |
| parsopercularis | WMV-slope | 0.30 | 0.0089 | 0.010 |
| parstriangularis | WMV-slope | 0.46 | < 0.001 | < 0.001 |
| superiortemporal | WMV-slope | 0.38 | < 0.001 | 0.0016 |
| supramarginal | WMV-slope | 0.33 | 0.0034 | 0.0049 |
| inferiorparietal | SA-intercept | 0.32 | 0.0053 | 0.021 |
| parstriangularis | SA-intercept | 0.36 | 0.0017 | 0.014 |
| superiortemporal | SA-intercept | 0.28 | 0.016 | 0.042 |
| inferiorparietal | SA-slope | 0.33 | 0.0031 | 0.013 |
| parstriangularis | SA-slope | 0.37 | 0.0012 | 0.0097 |
| superiortemporal | SA-slope | 0.27 | 0.019 | 0.050 |

**Supplementary Table 7.**
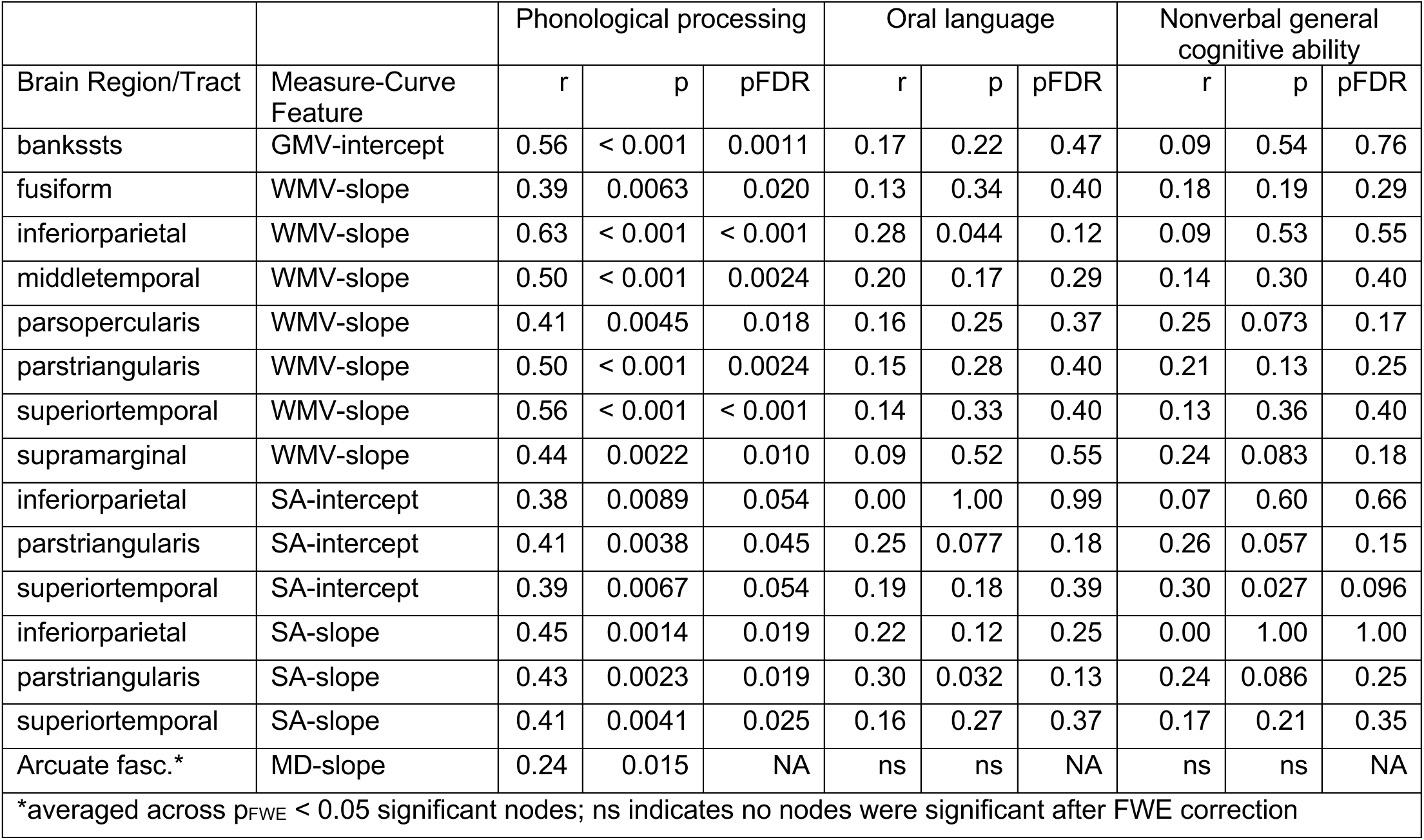
Associations between brain measures and literacy-related and cognitive (sub)skills.

**Supplementary Table 8.** Contributions of literacy-related covariates to growth curves.

|  |  |  |  |  |  |
| --- | --- | --- | --- | --- | --- |
|  |  | FHD |  | HLE |  |
| Brain Region/Tract | Measure | Effect size | p | Effect size | p |
| bankssts | GMV | $6.8 \times 10^{+01}$ | 0.59 | $2.2 \times 10^{+02}$ | 0.17 |
| fusiform | WMV | $1.0 \times 10^{+02}$ | 0.51 | $1.4 \times 10^{+02}$ | 0.48 |
| inferiorparietal | WMV | $-1.1 \times 10^{+02}$ | 0.67 | $3.9 \times 10^{+02}$ | 0.25 |
| middletemporal | WMV | $-8.4 \times 10^{+01}$ | 0.65 | $5.5 \times 10^{+02}$ | 0.021 |
| parsopercularis | WMV | $7.0 \times 10^{+01}$ | 0.42 | $9.6 \times 10^{+00}$ | 0.93 |
| parstriangularis | WMV | $1.9 \times 10^{+02}$ | 0.025 | $-3.0 \times 10^{+01}$ | 0.79 |
| superiortemporal | WMV | $3.4 \times 10^{+01}$ | 0.86 | $1.8 \times 10^{+02}$ | 0.48 |
| supramarginal | WMV | $-2.7 \times 10^{+02}$ | 0.32 | $3.5 \times 10^{+02}$ | 0.32 |
| inferiorparietal | SA | $1.0 \times 10^{+01}$ | 0.95 | $2.2 \times 10^{+02}$ | 0.28 |
| parstriangularis | SA | $8.2 \times 10^{+01}$ | 0.031 | $5.3 \times 10^{+00}$ | 0.92 |
| superiortemporal | SA | $6.5 \times 10^{+01}$ | 0.51 | $1.0 \times 10^{+02}$ | 0.42 |
| Arcuate fasc. | MD | ns | ns | ns | ns |
| FHD, family history of reading difficulty |  |  |  |  |  |
| HLE, home literacy environment |  |  |  |  |  |
| ns indicates no nodes were significant after FWE correction |  |  |  |  |  |

**Supplementary Table 9.** Comparison of Linear Mixed Effects Models with versus without Literacy-Related Covariates.

|  |  |  |  |  |
| --- | --- | --- | --- | --- |
| Brain Region/Tract | Measure | without covariate | with covariate | p |
| **A. Family history of reading difficulty** |  |  |  |  |
| bankssts | GMV | **2328.2** | 2332.5 | 0.37 |
| fusiform | WMV | **2505.3** | 2509.9 | 0.51 |
| inferiorparietal | WMV | **2670.1** | 2675.0 | 0.67 |
| middletemporal | WMV | **2496.7** | 2501.6 | 0.66 |
| parsopercularis | WMV | **2311.7** | 2315.9 | 0.34 |
| parstriangularis | WMV | **2274.7** | 2275.0 | 0.028 |
| superiortemporal | WMV | **2530.3** | 2535.3 | 0.86 |
| supramarginal | WMV | **2647.6** | 2651.3 | 0.24 |
| inferiorparietal | SA | **2451.3** | 2456.4 | 0.97 |
| parstriangularis | SA | **2012.9** | 2013.6 | 0.036 |
| superiortemporal | SA | **2354.4** | 2359.0 | 0.50 |
| Arcuate fasc. | MD | **-3358.7** | -3354.7 | 0.26 |
| **B. Home literacy Environment** |  |  |  |  |
| bankssts | GMV | **2328.2** | 2331.4 | 0.17 |
| fusiform | WMV | **2505.3** | 2509.9 | 0.49 |
| inferiorparietal | WMV | **2670.1** | 2673.9 | 0.25 |
| middletemporal | WMV | 2496.7 | **2496.4** | 0.021 |
| parsopercularis | WMV | **2311.7** | 2316.8 | 0.93 |
| parstriangularis | WMV | **2274.7** | 2279.7 | 0.79 |
| superiortemporal | WMV | **2530.3** | 2534.8 | 0.48 |
| supramarginal | WMV | **2647.6** | 2651.8 | 0.32 |
| inferiorparietal | SA | **2451.3** | 2455.3 | 0.29 |
| parstriangularis | SA | **2012.9** | 2018.0 | 0.92 |
| superiortemporal | SA | **2354.4** | 2358.8 | 0.42 |
| Arcuate fasc. | MD | **-3358.7** | -3353.4 | 1.00 |
| Bold indicates lower BIC value |  |  |  |  |

**Supplementary Table 10.** Indirect effects between brain structure and decoding via phonological processing.

|  |  |  |  |  |  |
| --- | --- | --- | --- | --- | --- |
| Brain Region/Tract | Measure-Curve Feature | Effect size | Lower CI | Upper CI | p |
| bankssts | GMV-intercept | $1.7 \times 10^{-02}$ | $3.4 \times 10^{-03}$ | $3.8 \times 10^{-02}$ | 0.011 |
| fusiform | WMV-slope | $6.8 \times 10^{-03}$ | $-1.0 \times 10^{-03}$ | $1.9 \times 10^{-02}$ | 0.094 |
| inferiorparietal | WMV-slope | $9.1 \times 10^{-03}$ | $5.5 \times 10^{-04}$ | $2.2 \times 10^{-02}$ | 0.032 |
| middletemporal | WMV-slope | $9.4 \times 10^{-03}$ | $6.1 \times 10^{-04}$ | $2.3 \times 10^{-02}$ | 0.034 |
| parsopercularis | WMV-slope | $2.0 \times 10^{-02}$ | $3.0 \times 10^{-03}$ | $4.1 \times 10^{-02}$ | 0.016 |
| parstriangularis | WMV-slope | $2.2 \times 10^{-02}$ | $2.8 \times 10^{-03}$ | $4.9 \times 10^{-02}$ | 0.016 |
| superiortemporal | WMV-slope | $8.3 \times 10^{-03}$ | $7.8 \times 10^{-04}$ | $2.2 \times 10^{-02}$ | 0.020 |
| supramarginal | WMV-slope | $5.0 \times 10^{-03}$ | $-1.3 \times 10^{-03}$ | $1.3 \times 10^{-02}$ | 0.12 |
| inferiorparietal | SA-intercept | $8.5 \times 10^{-03}$ | $-9.7 \times 10^{-04}$ | $1.5 \times 10^{-02}$ | 0.088 |
| parstriangularis | SA-intercept | $4.6 \times 10^{-02}$ | $2.3 \times 10^{-03}$ | $1.0 \times 10^{-01}$ | 0.036 |
| superiortemporal | SA-intercept | $1.6 \times 10^{-02}$ | $-1.9 \times 10^{-03}$ | $3.6 \times 10^{-02}$ | 0.083 |
| inferiorparietal | SA-slope | $2.3 \times 10^{-02}$ | $2.0 \times 10^{-03}$ | $5.9 \times 10^{-02}$ | 0.024 |
| parstriangularis | SA-slope | $7.4 \times 10^{-02}$ | $3.7 \times 10^{-03}$ | $1.8 \times 10^{-01}$ | 0.034 |
| superiortemporal | SA-slope | $1.8 \times 10^{-02}$ | $-3.8 \times 10^{-03}$ | $4.6 \times 10^{-02}$ | 0.11 |
| Arcuate fasc.* | MD-slope | $2.6 \times 10^{+05}$ | $6.8 \times 10^{+04}$ | $6.2 \times 10^{+05}$ | 0.014 |
| *averaged across $p_{FWE} < 0.05$ significant nodes | | | | | |

**Supplementary Table 11.** Indirect effects between brain structure and word reading via phonological processing.

| Brain Region/Tract | Measure-Curve Feature | Effect size | Lower CI | Upper CI | p |
| --- | --- | --- | --- | --- | --- |
| bankssts | GMV-intercept | $2.6 \times 10^{-02}$ | $9.2 \times 10^{-03}$ | $5.4 \times 10^{-02}$ | < 0.001 |
| fusiform | WMV-slope | $9.7 \times 10^{-03}$ | $-1.3 \times 10^{-03}$ | $2.5 \times 10^{-02}$ | 0.083 |
| inferiorparietal | WMV-slope | $1.4 \times 10^{-02}$ | $2.7 \times 10^{-03}$ | $3.2 \times 10^{-02}$ | 0.0070 |
| middletemporal | WMV-slope | $1.4 \times 10^{-02}$ | $1.6 \times 10^{-03}$ | $3.3 \times 10^{-02}$ | 0.026 |
| parsopercularis | WMV-slope | $2.9 \times 10^{-02}$ | $6.2 \times 10^{-03}$ | $5.8 \times 10^{-02}$ | 0.011 |
| parstriangularis | WMV-slope | $3.4 \times 10^{-02}$ | $6.9 \times 10^{-03}$ | $7.2 \times 10^{-02}$ | 0.013 |
| superiortemporal | WMV-slope | $1.3 \times 10^{-02}$ | $3.2 \times 10^{-03}$ | $3.2 \times 10^{-02}$ | 0.0042 |
| supramarginal | WMV-slope | $7.1 \times 10^{-03}$ | $-1.6 \times 10^{-03}$ | $1.8 \times 10^{-02}$ | 0.10 |
| inferiorparietal | SA-intercept | $1.2 \times 10^{-02}$ | $3.8 \times 10^{-04}$ | $2.1 \times 10^{-02}$ | 0.045 |
| parstriangularis | SA-intercept | $6.6 \times 10^{-02}$ | $3.3 \times 10^{-03}$ | $1.4 \times 10^{-01}$ | 0.041 |
| superiortemporal | SA-intercept | $2.6 \times 10^{-02}$ | $3.1 \times 10^{-03}$ | $4.9 \times 10^{-02}$ | 0.026 |
| inferiorparietal | SA-slope | $3.7 \times 10^{-02}$ | $7.1 \times 10^{-03}$ | $8.6 \times 10^{-02}$ | 0.013 |
| parstriangularis | SA-slope | $1.1 \times 10^{-01}$ | $5.3 \times 10^{-03}$ | $2.5 \times 10^{-01}$ | 0.039 |
| superiortemporal | SA-slope | $3.1 \times 10^{-02}$ | $-3.0 \times 10^{-04}$ | $6.6 \times 10^{-02}$ | 0.052 |
| Arcuate fasc.* | MD-slope | $3.1 \times 10^{+05}$ | $7.0 \times 10^{+04}$ | $6.8 \times 10^{+05}$ | 0.020 |

\*averaged across $p_{\text{FWE}} < 0.05$ significant nodes

**Supplementary Table 12.** Indirect effects between brain and decoding via phonological processing controlling for TIV.

|  |  |  |  |  |  |
| --- | --- | --- | --- | --- | --- |
| Brain Region | Measure-Curve Feature | Effect size | Lower CI | Upper CI | p |
| bankssts | GMV-intercept | $1.7 \times 10^{-02}$ | $3.2 \times 10^{-03}$ | $3.8 \times 10^{-02}$ | 0.010 |
| fusiform | WMV-slope | $3.5 \times 10^{-03}$ | $-4.6 \times 10^{-03}$ | $1.6 \times 10^{-02}$ | 0.37 |
| inferiorparietal | WMV-slope | $8.5 \times 10^{-03}$ | $-1.5 \times 10^{-05}$ | $2.5 \times 10^{-02}$ | 0.051 |
| middletemporal | WMV-slope | $7.1 \times 10^{-03}$ | $-3.1 \times 10^{-03}$ | $2.3 \times 10^{-02}$ | 0.16 |
| parsopercularis | WMV-slope | $1.4 \times 10^{-02}$ | $-5.3 \times 10^{-04}$ | $3.7 \times 10^{-02}$ | 0.065 |
| parstriangularis | WMV-slope | $1.7 \times 10^{-02}$ | $-1.7 \times 10^{-03}$ | $4.6 \times 10^{-02}$ | 0.085 |
| superiortemporal | WMV-slope | $7.7 \times 10^{-03}$ | $-5.2 \times 10^{-04}$ | $2.2 \times 10^{-02}$ | 0.079 |
| supramarginal | WMV-slope | $2.2 \times 10^{-03}$ | $-5.9 \times 10^{-03}$ | $1.1 \times 10^{-02}$ | 0.56 |
| inferiorparietal | SA-intercept | $8.4 \times 10^{-03}$ | $-1.2 \times 10^{-03}$ | $1.5 \times 10^{-02}$ | 0.10 |
| parstriangularis | SA-intercept | $4.5 \times 10^{-02}$ | $1.6 \times 10^{-03}$ | $1.1 \times 10^{-01}$ | 0.039 |
| superiortemporal | SA-intercept | $1.6 \times 10^{-02}$ | $-3.4 \times 10^{-03}$ | $3.7 \times 10^{-02}$ | 0.10 |
| inferiorparietal | SA-slope | $1.8 \times 10^{-02}$ | $-1.4 \times 10^{-03}$ | $5.9 \times 10^{-02}$ | 0.078 |
| parstriangularis | SA-slope | $5.2 \times 10^{-02}$ | $-1.2 \times 10^{-02}$ | $1.7 \times 10^{-01}$ | 0.14 |
| superiortemporal | SA-slope | $8.3 \times 10^{-03}$ | $-3.1 \times 10^{-02}$ | $4.3 \times 10^{-02}$ | 0.67 |

**Supplementary Table 13.** Indirect effects between brain and word reading via phonological processing controlling for TIV.

|  |  |  |  |  |  |
| --- | --- | --- | --- | --- | --- |
| Brain Region | Measure-Curve Feature | Effect size | Lower CI | Upper CI | p |
| bankssts | GMV-intercept | $2.6 \times 10^{-02}$ | $8.4 \times 10^{-03}$ | $5.3 \times 10^{-02}$ | 0.0017 |
| fusiform | WMV-slope | $5.0 \times 10^{-03}$ | $-6.6 \times 10^{-03}$ | $2.0 \times 10^{-02}$ | 0.38 |
| inferiorparietal | WMV-slope | $1.3 \times 10^{-02}$ | $1.4 \times 10^{-03}$ | $3.3 \times 10^{-02}$ | 0.020 |
| middletemporal | WMV-slope | $1.0 \times 10^{-02}$ | $-4.9 \times 10^{-03}$ | $3.2 \times 10^{-02}$ | 0.15 |
| parsopercularis | WMV-slope | $2.1 \times 10^{-02}$ | $-8.0 \times 10^{-04}$ | $4.9 \times 10^{-02}$ | 0.060 |
| parstriangularis | WMV-slope | $2.6 \times 10^{-02}$ | $-3.8 \times 10^{-03}$ | $6.3 \times 10^{-02}$ | 0.093 |
| superiortemporal | WMV-slope | $1.2 \times 10^{-02}$ | $-2.4 \times 10^{-04}$ | $3.2 \times 10^{-02}$ | 0.056 |
| supramarginal | WMV-slope | $2.9 \times 10^{-03}$ | $-8.7 \times 10^{-03}$ | $1.4 \times 10^{-02}$ | 0.57 |
| inferiorparietal | SA-intercept | $1.2 \times 10^{-02}$ | $-6.5 \times 10^{-05}$ | $2.1 \times 10^{-02}$ | 0.051 |
| parstriangularis | SA-intercept | $6.6 \times 10^{-02}$ | $2.5 \times 10^{-03}$ | $1.4 \times 10^{-01}$ | 0.042 |
| superiortemporal | SA-intercept | $2.6 \times 10^{-02}$ | $5.2 \times 10^{-04}$ | $5.1 \times 10^{-02}$ | 0.047 |
| inferiorparietal | SA-slope | $2.8 \times 10^{-02}$ | $-1.3 \times 10^{-03}$ | $8.5 \times 10^{-02}$ | 0.061 |
| parstriangularis | SA-slope | $7.2 \times 10^{-02}$ | $-2.6 \times 10^{-02}$ | $2.2 \times 10^{-01}$ | 0.17 |
| superiortemporal | SA-slope | $1.4 \times 10^{-02}$ | $-4.3 \times 10^{-02}$ | $6.4 \times 10^{-02}$ | 0.60 |

## Supplementary Figures

**Supplementary Figure 1.**
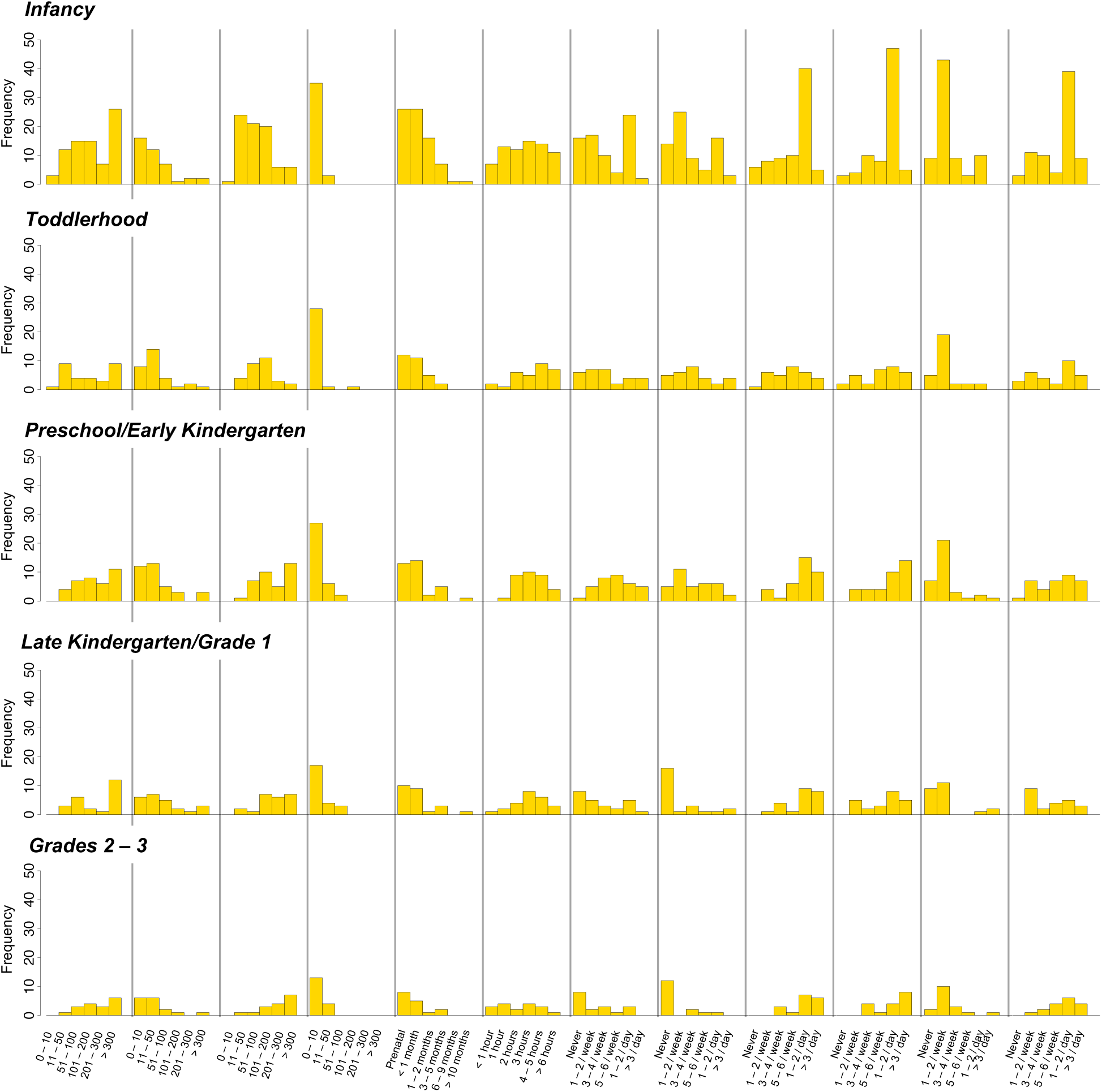
Distribution of home literacy environment variables. Histograms depict the distribution of home literacy environment questionnaire responses across five developmental timepoints. New England data only, as these data were not collected in the Calgary dataset.

**Supplementary Figure 2.**
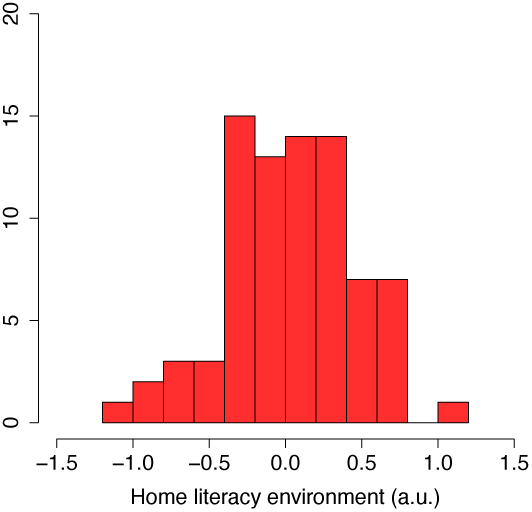
Distribution of home literacy environment scores. Histogram depicts the distribution of home literacy environment scores normalized from the responses shown in Supplementary Figure 1. New England data only, as these data were not collected in the Calgary dataset.

**Supplementary Figure 3.**
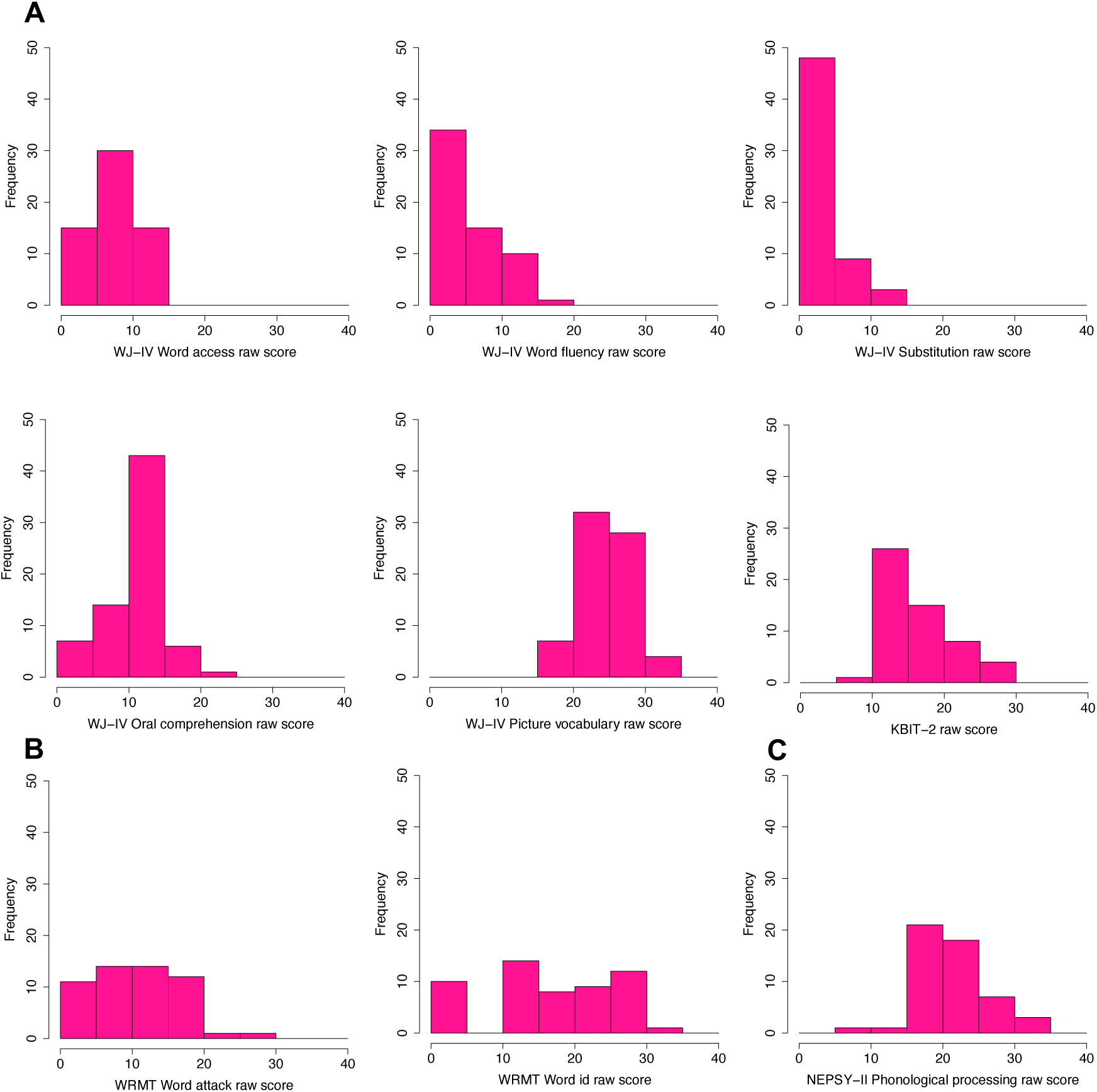
Distribution of raw literacy-related outcomes. Histograms depict the distributions of New England data for (A) subtests constituting phonological processing and oral language composite scores and KBIT-2, and (B) word ID and word attack assessments. (C) Histogram for raw phonological processing scores from the Calgary dataset. WJ-IV, Woodcock-Johnson edition IV; KBIT-2, Kaufman Brief Intelligence Test edition 2; WRMT, Woodcock Reading Mastery Tests; NEPSY-II, Neuropsychological Assessment.

**Supplementary Figure 4.**
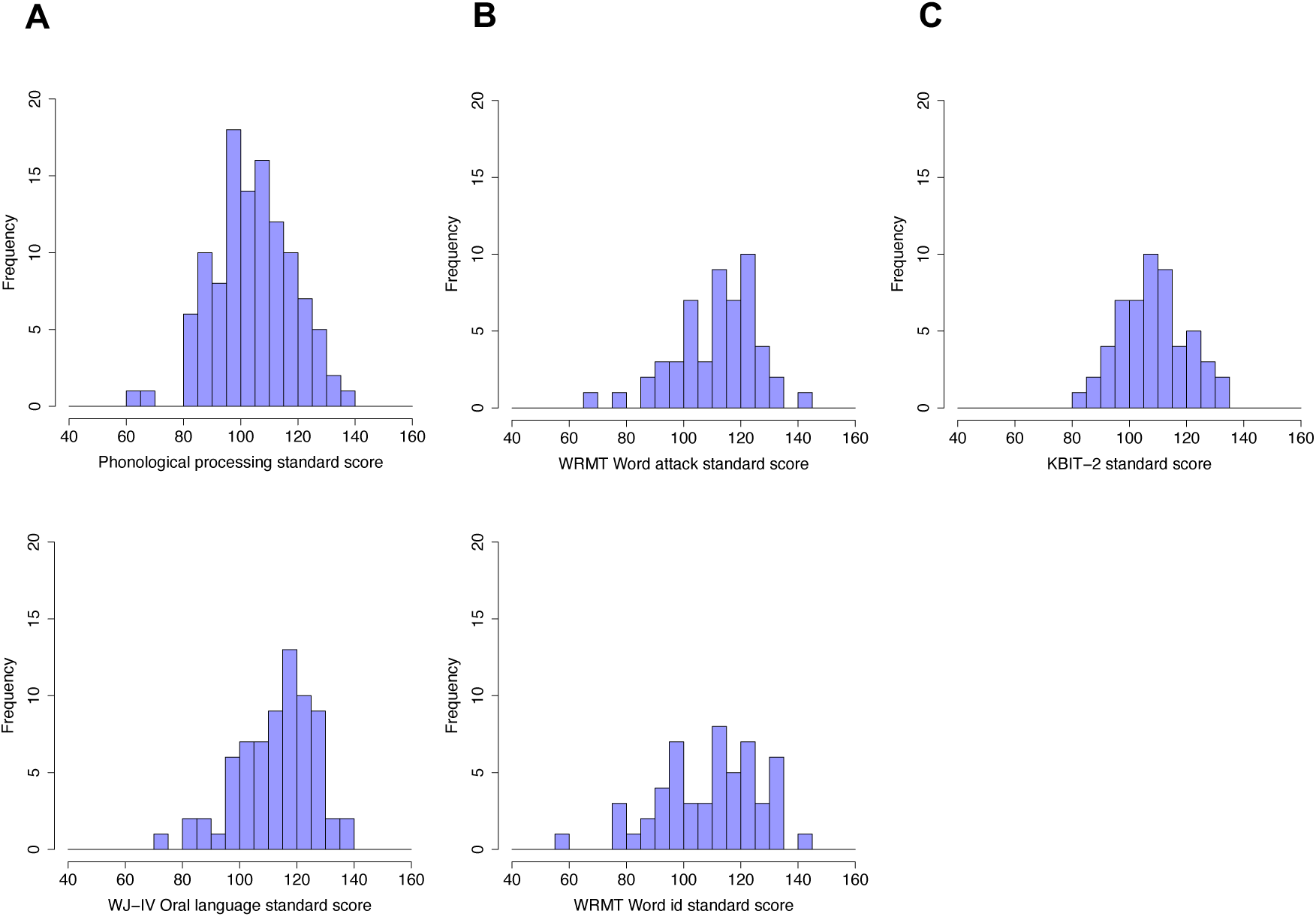
Distribution of standardized literacy-related outcomes. Histograms depict the distributions of for (A) literacy-related subskills, (B) decoding/word reading assessments and (C) nonverbal general cognitive abilities. Phonological processing scores sum WJ-IV estimates from the New England dataset and NEPSY-II estimates from the Calgary dataset. WJ-IV, Woodcock-Johnson edition IV; NEPSY-II, Neuropsychological Assessment; WRMT, Woodcock Reading Mastery Tests; KBIT-2, Kaufman Brief Intelligence Test edition 2.

**Supplementary Figure 5.**
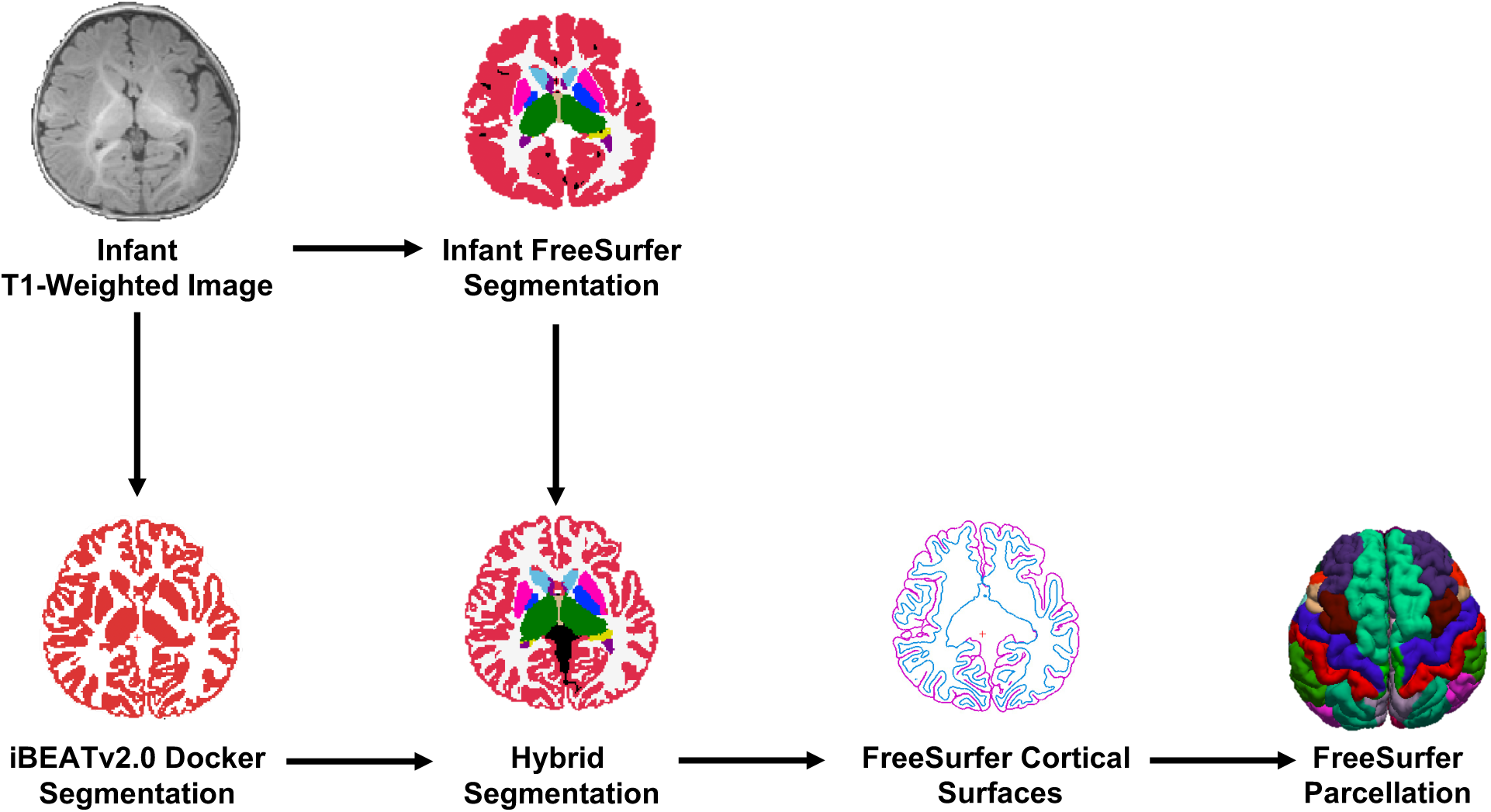
Infant Structural Processing Pipeline Overview. Raw images without visual artifacts were processed using a combination of iBEATv2.0 Docker, Infant FreeSurfer, FreeSurfer, and in-house scripts. Cortical surfaces were visually inspected for tissue classification accuracy (please see Methods section for details).

**Supplementary Figure 6.**
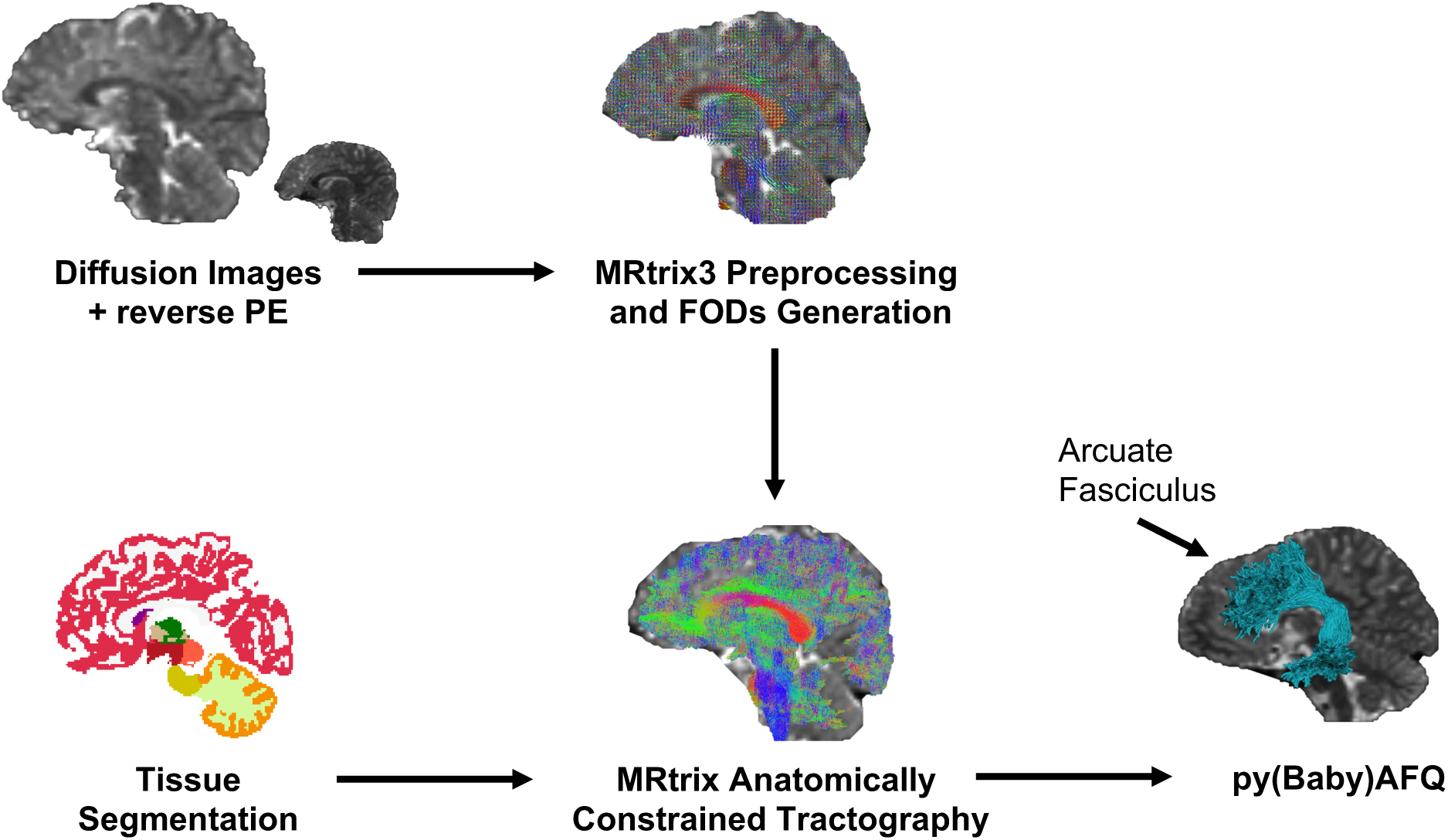
Diffusion Processing Pipeline Overview. Diffusion data were denoised and then corrected for susceptibility distortions, eddy currents, head motion, and intensity inhomogeneity using MRtrix with FSL and ANTs implementations. Fiber orientation densities (FODs) were generated using constrained spherical deconvolution. Fibers were tracked with Anatomically Constrained Tractography, which leveraged the tissue segmentations from the structural processing pipeline (Supplementary Figure 1): the hybrid segmentation for brains < 50 months and the standard FreeSurfer segmentation for brains > 50 months. Fibers were segmented into tracts using pyBabyAFQ for brains ≤ 24 months and pyAFQ for brains > 50 months (the left arcuate fasciculus is depicted as an example). PE, phase encoding.

**Supplementary Figure 7.**
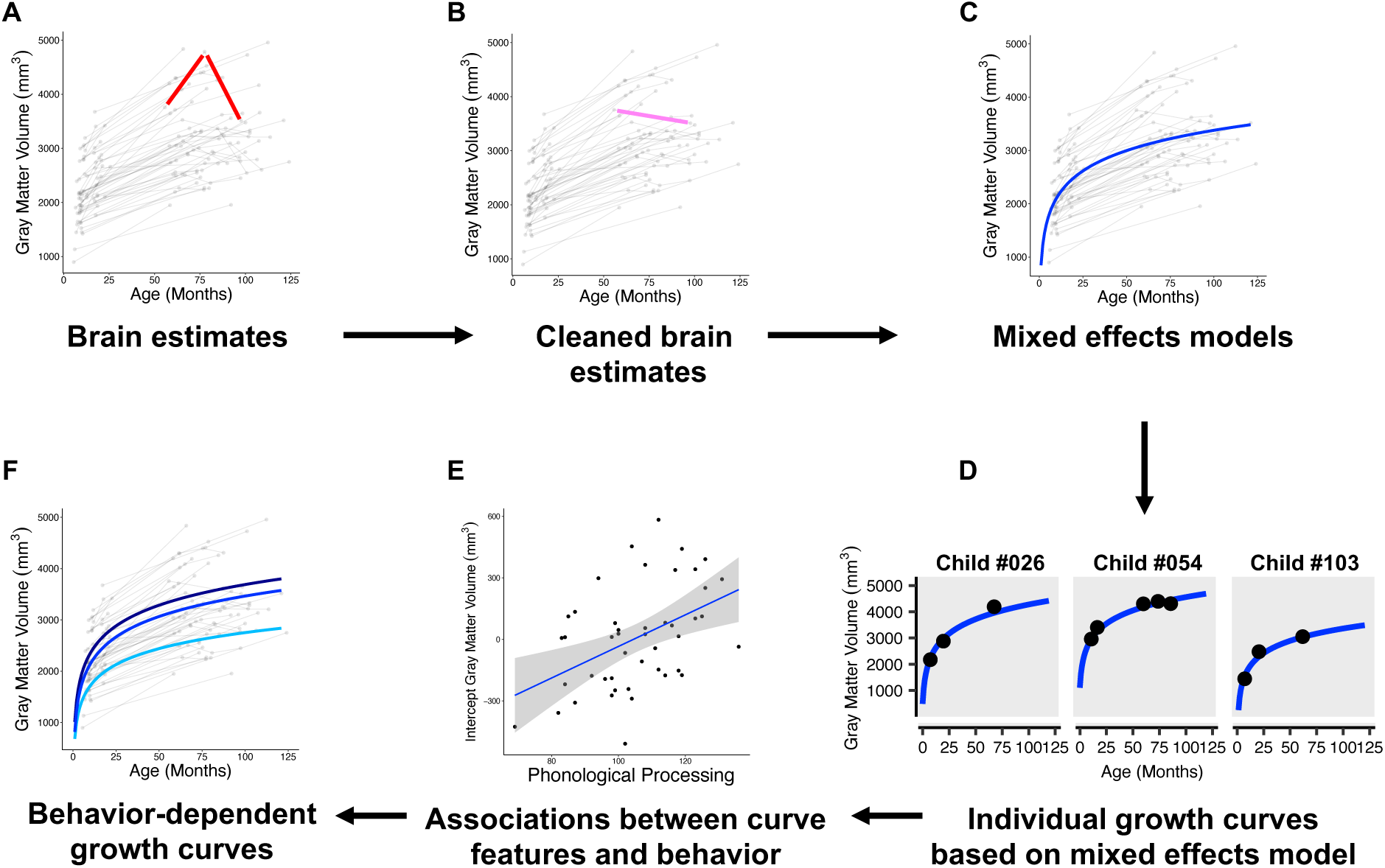
Statistical Analysis Overview. (A, B) Structural and diffusion brain estimates for regions and tracts of interest are first cleaned to remove observations with annual changes that are not neurophysiologically plausible (example inter-observation segments in red); remaining observations are then bridged (pink segment). Please see Methods for determining whether to remove the estimate preceding or following the problematic annual change. (C) Mixed effects models, here using logarithmic functions and random intercept and slope terms, were used to generate (D) individual growth curves. (E) Brain-behavior associations were tested using curve features (i.e., random intercepts and slopes). (F) Individual growth curves for measures and regions/tracts with significant brain-behavior associations were divided into low (< 85, dotted line), average (85 – 115, solid line), and high (> 115, dashed line) behavioral performance, meaned within groups, and plotted.

**Supplementary Figure 8.**
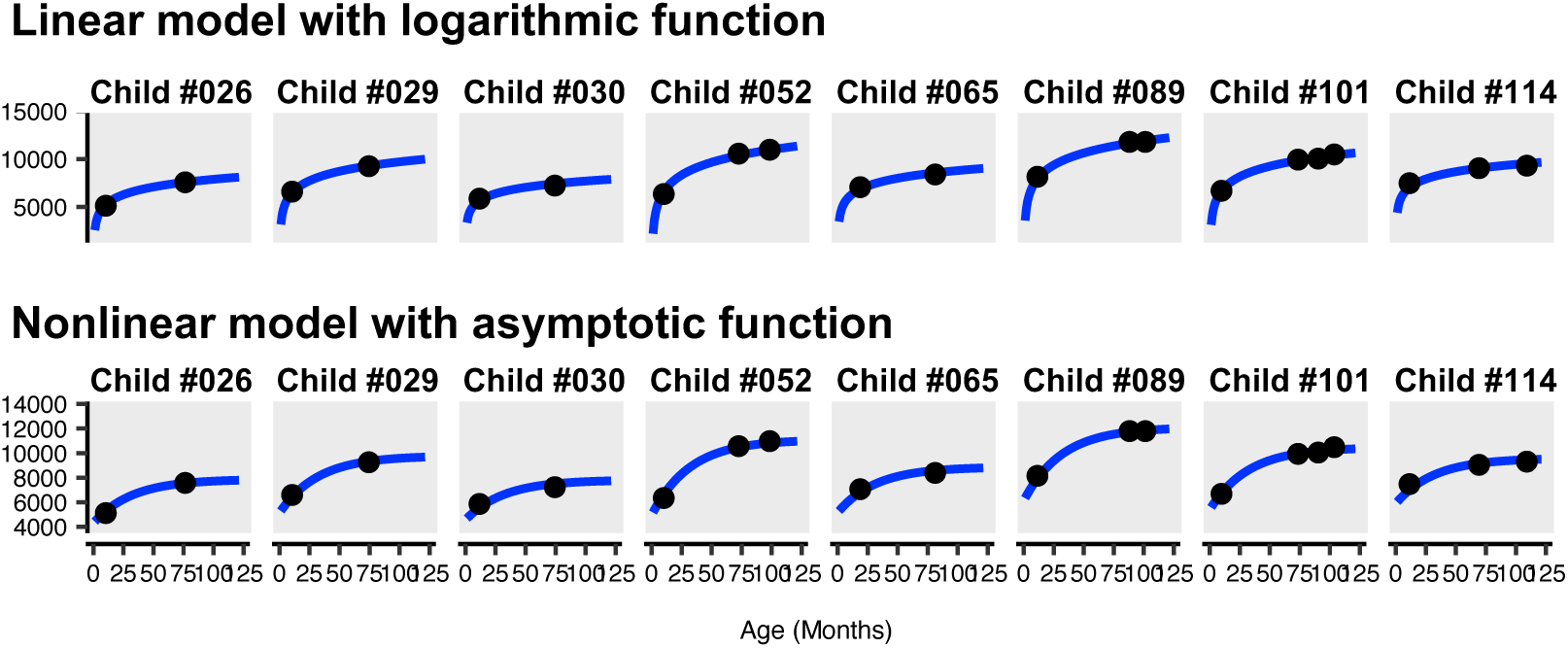
Individual growth curves for inferior parietal white matter in a subset of participants. Projected longitudinal trajectories (blue lines) show close fits to raw brain estimates (black dots) for both linear and nonlinear models.

**Supplementary Figure 9.**
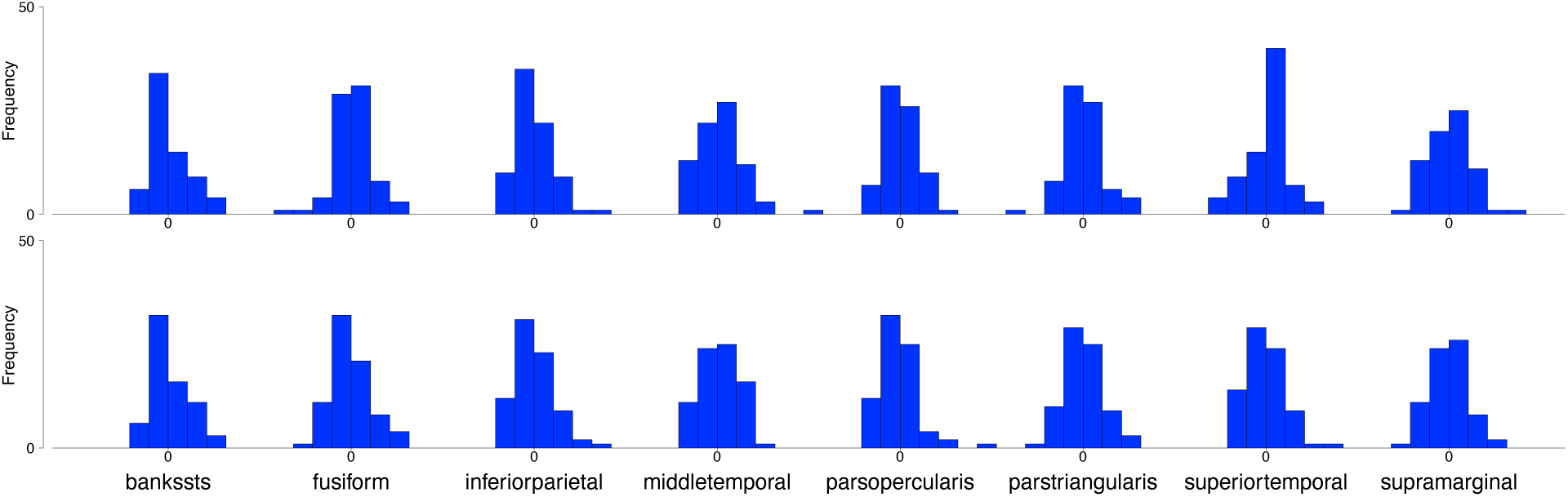
Distribution of curve features of gray matter volume. Top row depicts histograms for curve intercepts. Bottom row depicts histograms for curve slopes. Brain estimates were scaled for visualization purposes, but skewness was not altered.

**Supplementary Figure 10.**
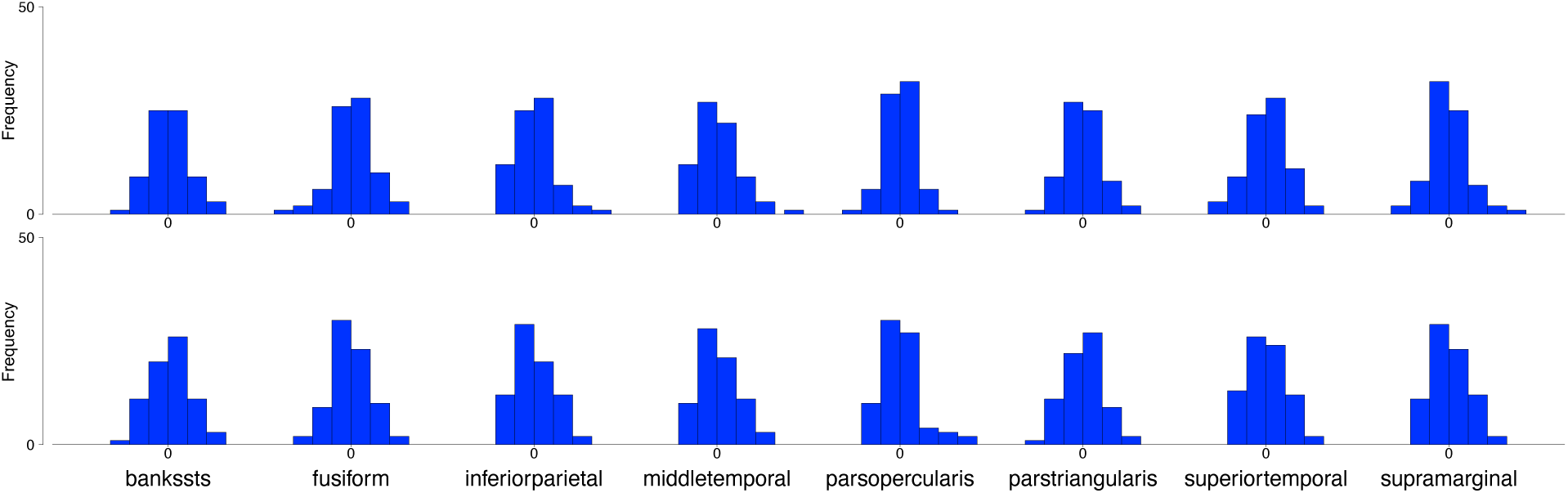
Distribution of curve features of white matter volume. Top row depicts histograms for curve intercepts. Bottom row depicts histograms for curve slopes. Brain estimates were scaled for visualization purposes, but skewness was not altered.

**Supplementary Figure 11.**
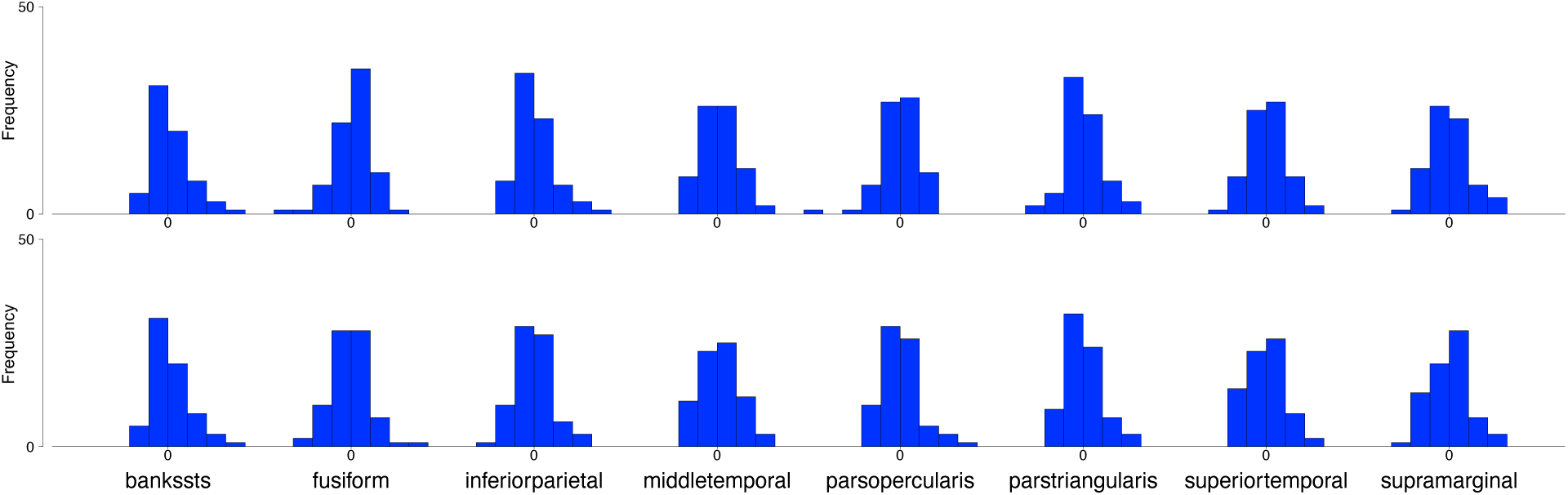
Distribution of curve features of surface area. Top row depicts histograms for curve intercepts. Bottom row depicts histograms for curve slopes. Brain estimates were scaled for visualization purposes, but skewness was not altered.

**Supplementary Figure 12.**
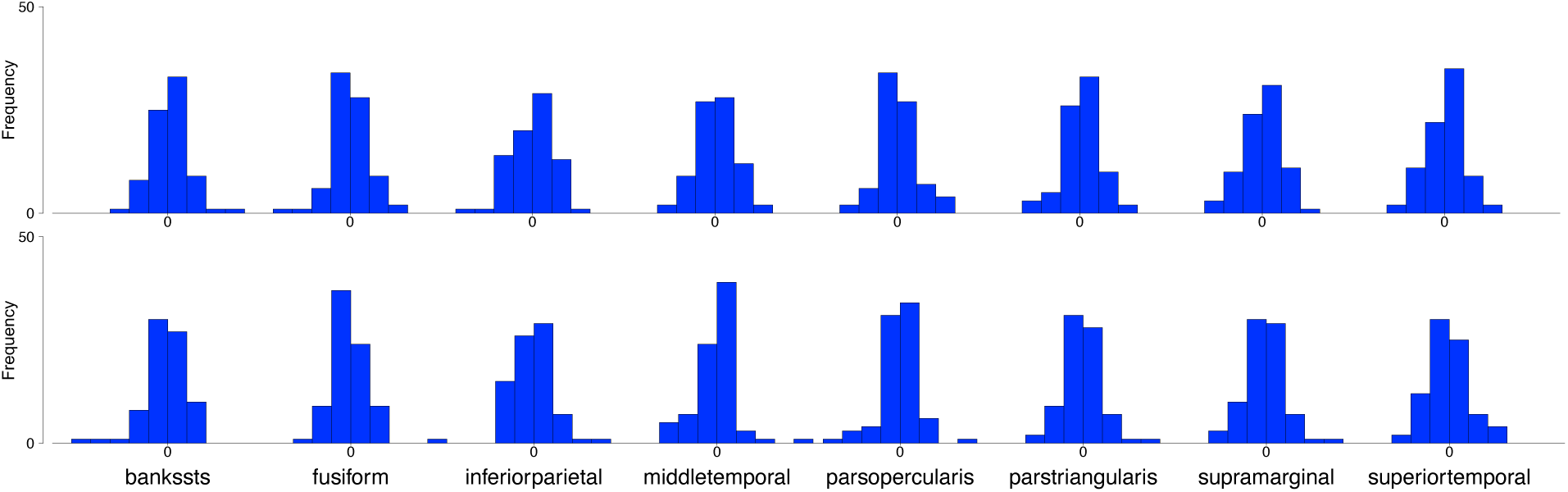
Distribution of curve features of cortical thickness. Top row depicts histograms for curve intercepts. Bottom row depicts histograms for curve slopes. Brain estimates were scaled for visualization purposes, but skewness was not altered.

**Supplementary Figure 13.**
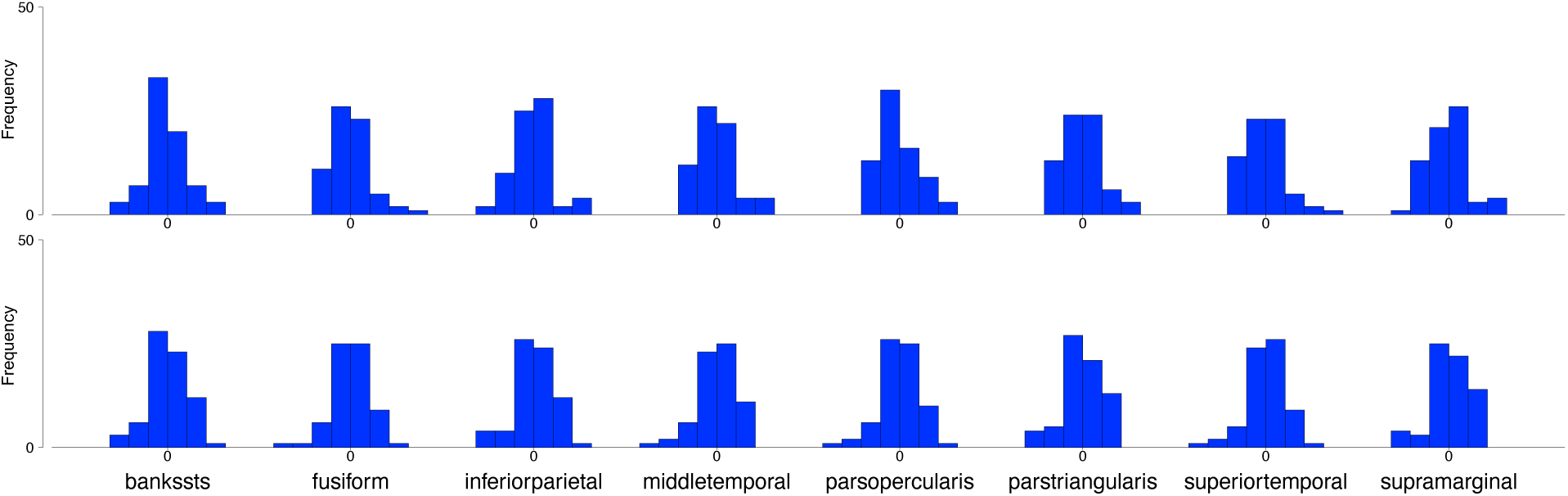
Distribution of curve features of mean curvature. Top row depicts histograms for curve intercepts. Bottom row depicts histograms for curve slopes. Brain estimates were scaled for visualization purposes, but skewness was not altered.

**Supplementary Figure 14.**
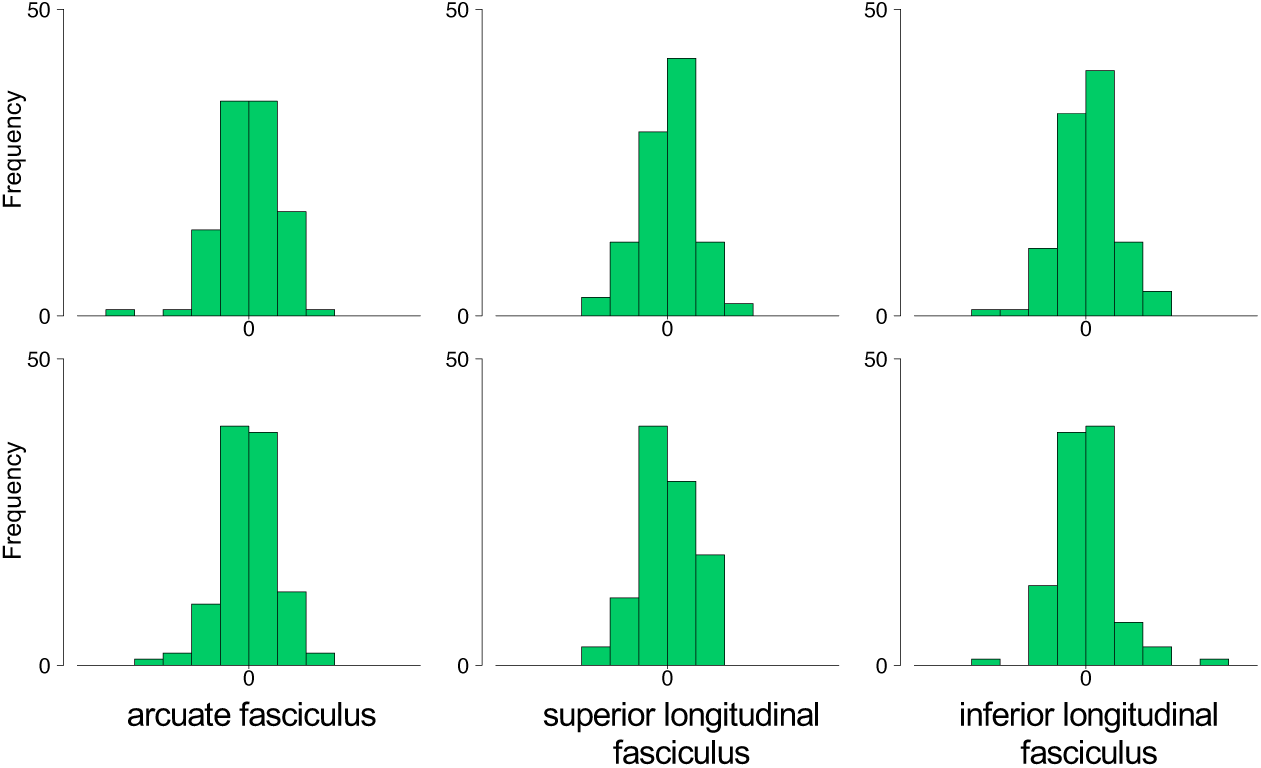
Distribution of curve features of fractional anisotropy. Top row depicts histograms for curve intercepts. Bottom row depicts histograms for curve slopes. Brain estimates were scaled for visualization purposes, but skewness was not altered.

**Supplementary Figure 15.**
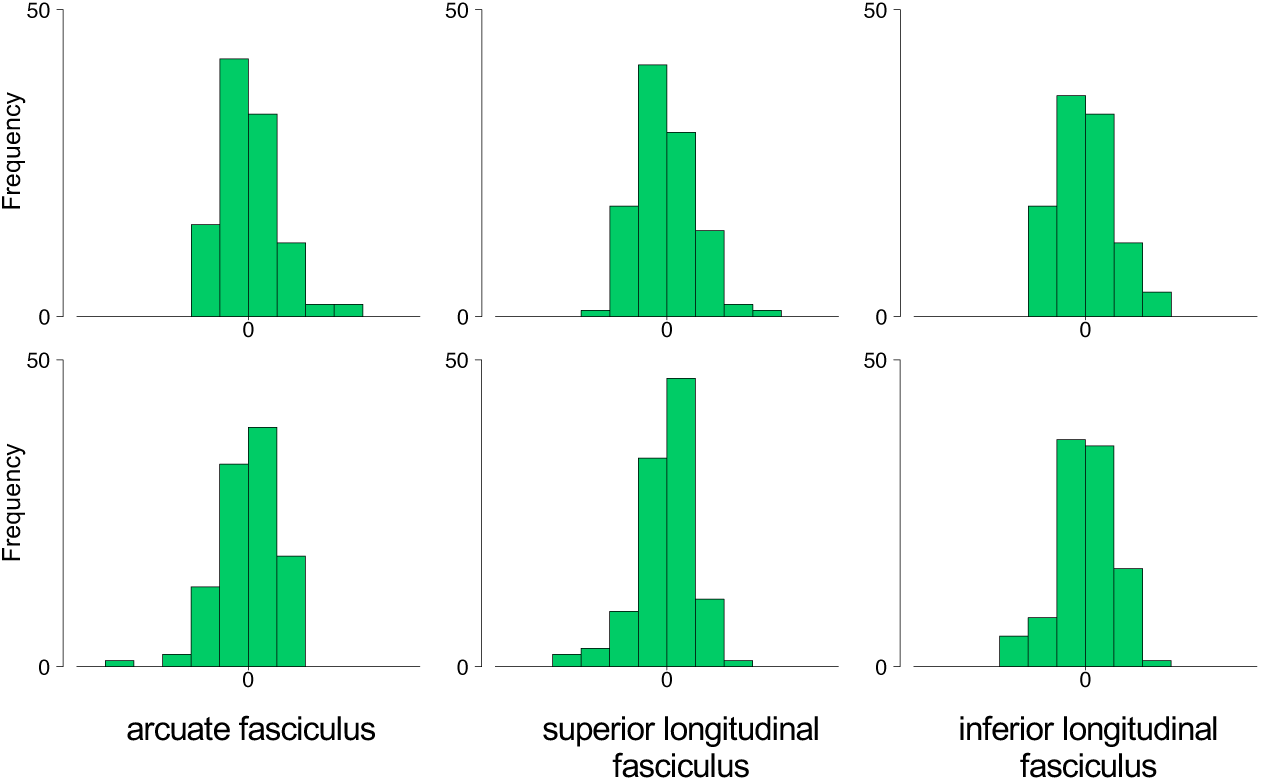
Distribution of curve features of mean diffusivity. Top row depicts histograms for curve intercepts. Bottom row depicts histograms for curve slopes. Brain estimates were scaled for visualization purposes, but skewness was not altered.

**Supplementary Figure 16.**
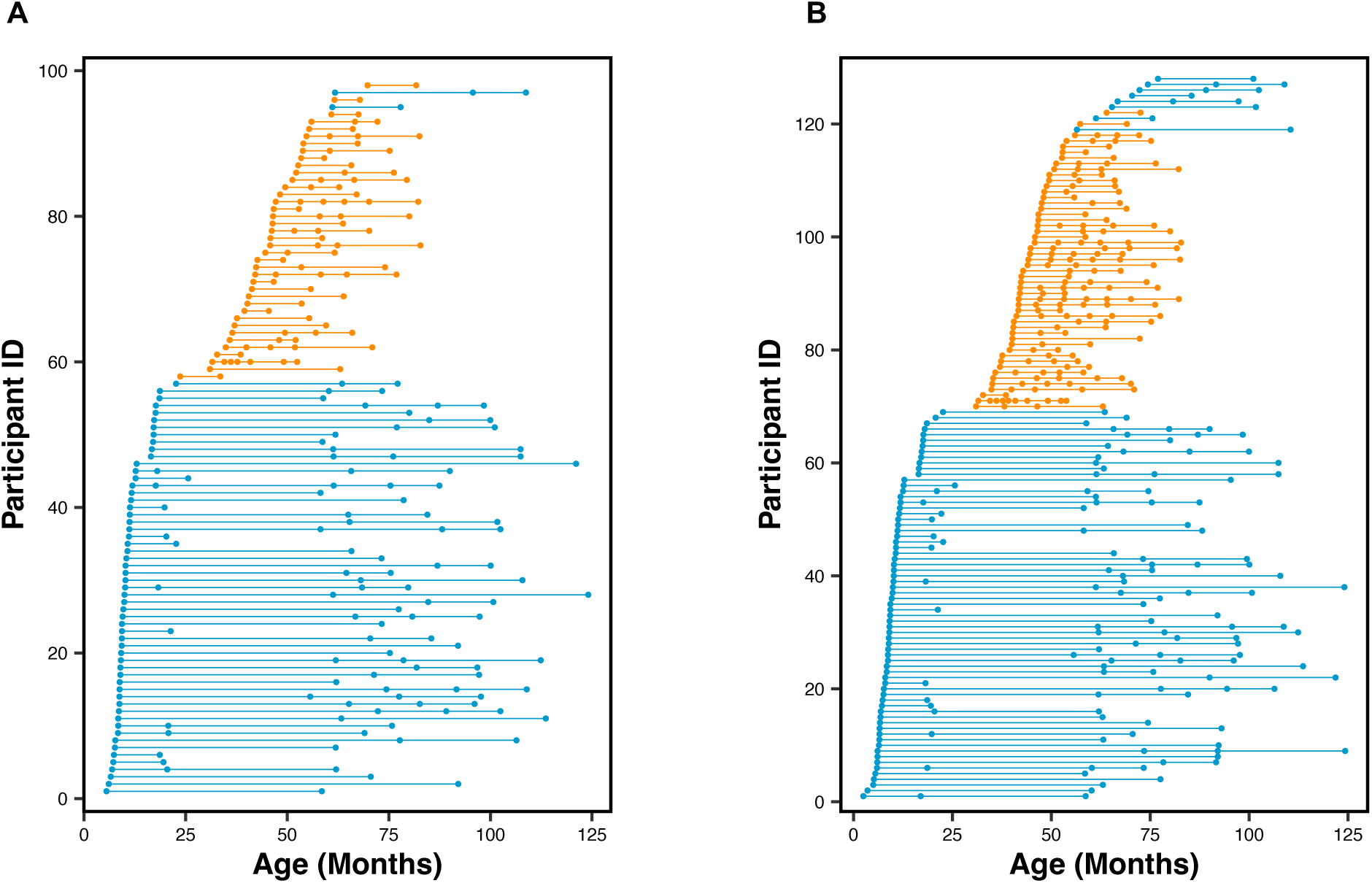
Age distributions by modality of longitudinal datasets from infancy to late childhood. All children had (A) structural and/or (B) diffusion MRI data from at least two observations (dots). Blue, New England cohort; orange, Calgary cohort.

**Supplementary Figure 17.**
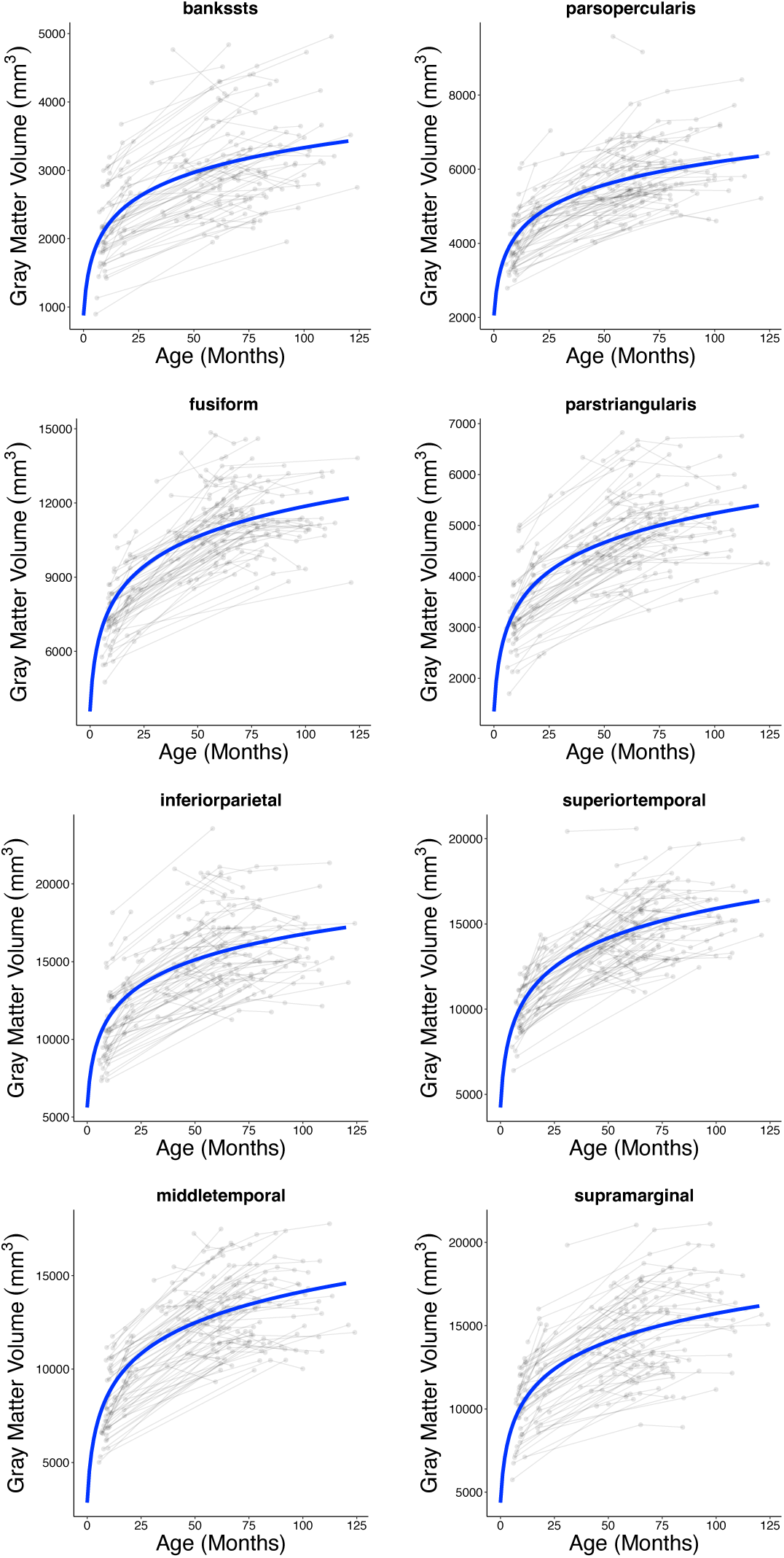
Longitudinal trajectories of gray matter volume between infancy and

**Supplementary Figure 18.**
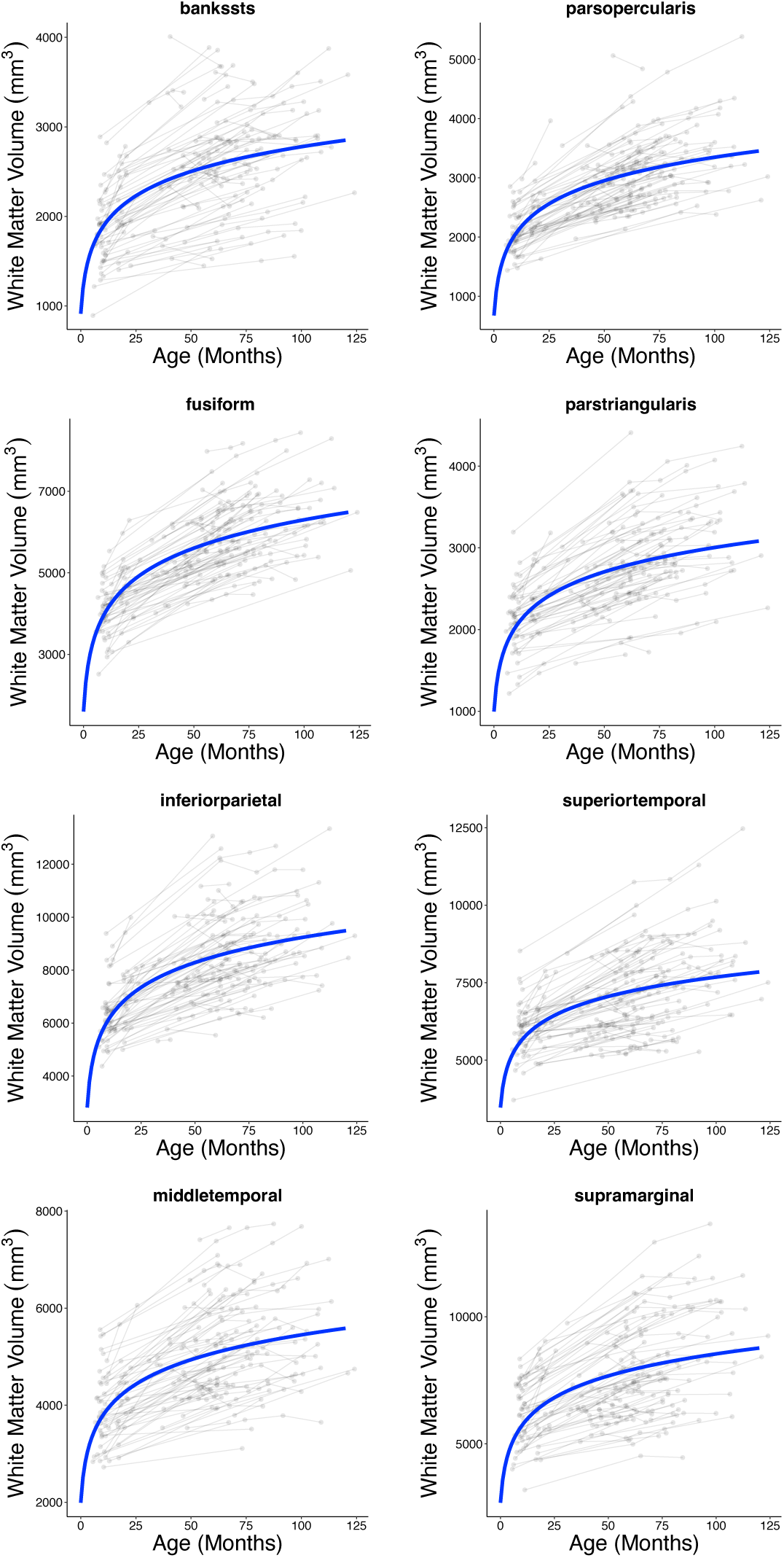
Longitudinal trajectories of white matter volume between infancy and

**Supplementary Figure 19.**
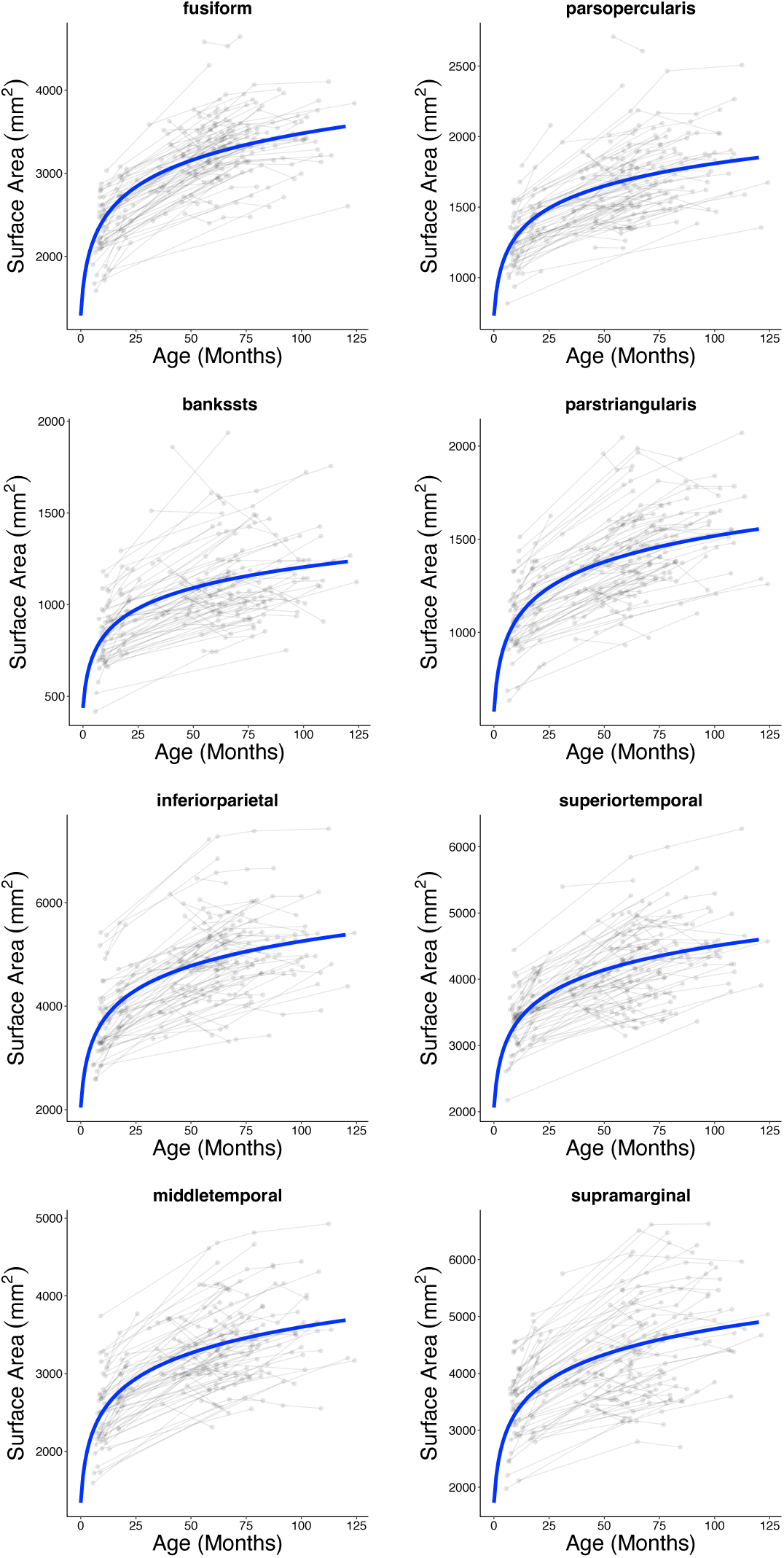
Longitudinal trajectories of surface area between infancy and late childhood. Raw estimates of surface area (gray spaghetti plot backdrop) were submitted to linear mixed effects models using a logarithmic function. Individual growth curves predicted by this

**Supplementary Figure 20.**
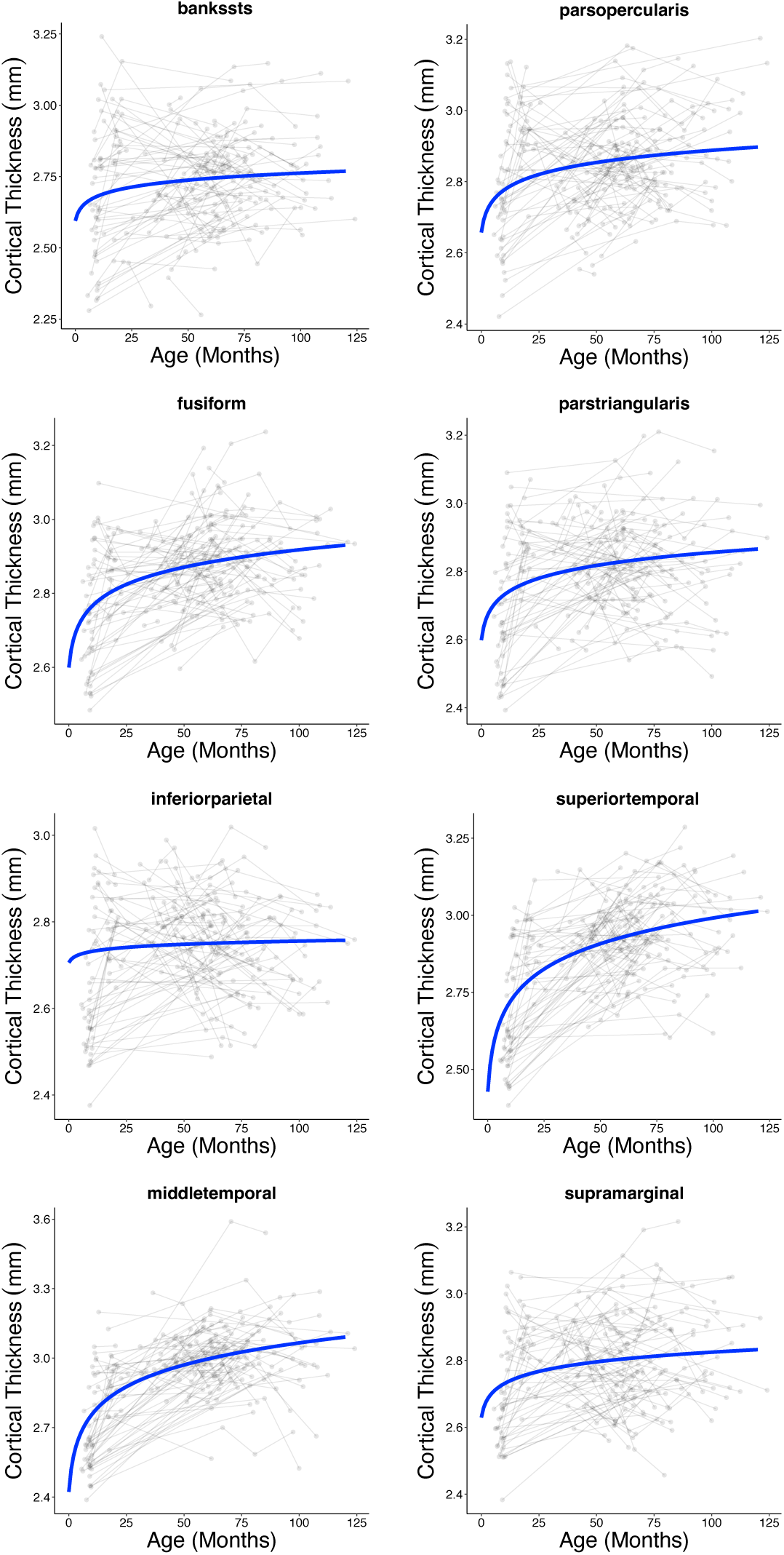
Longitudinal trajectories of cortical thickness between infancy and late childhood. Raw estimates of cortical thickness (gray spaghetti plot backdrop) were submitted

**Supplementary Figure 21.**
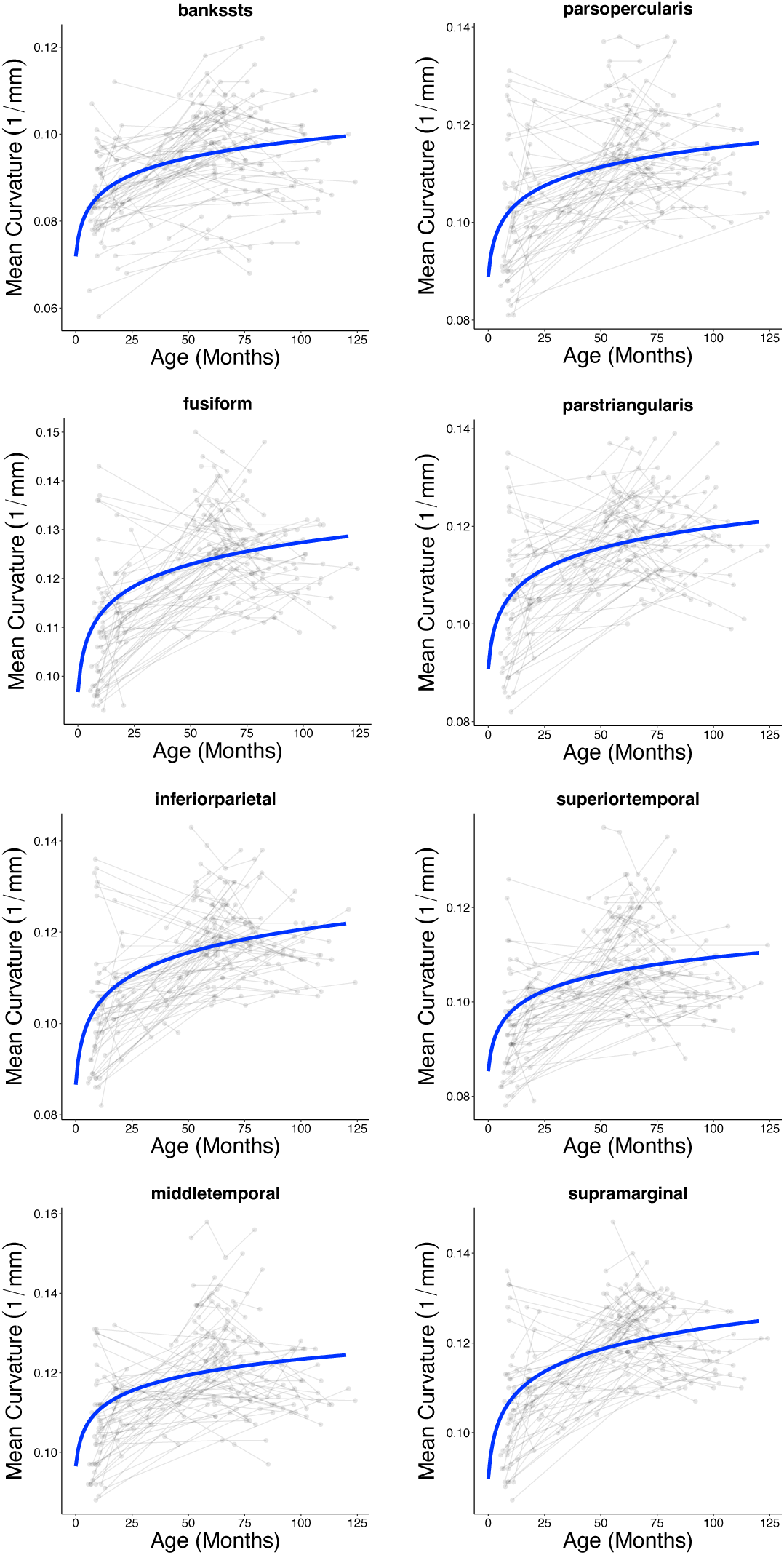
Longitudinal trajectories of mean curvature between infancy and late childhood. Raw estimates of mean curvature (gray spaghetti plot backdrop) were submitted to

**Supplementary Figure 22.**
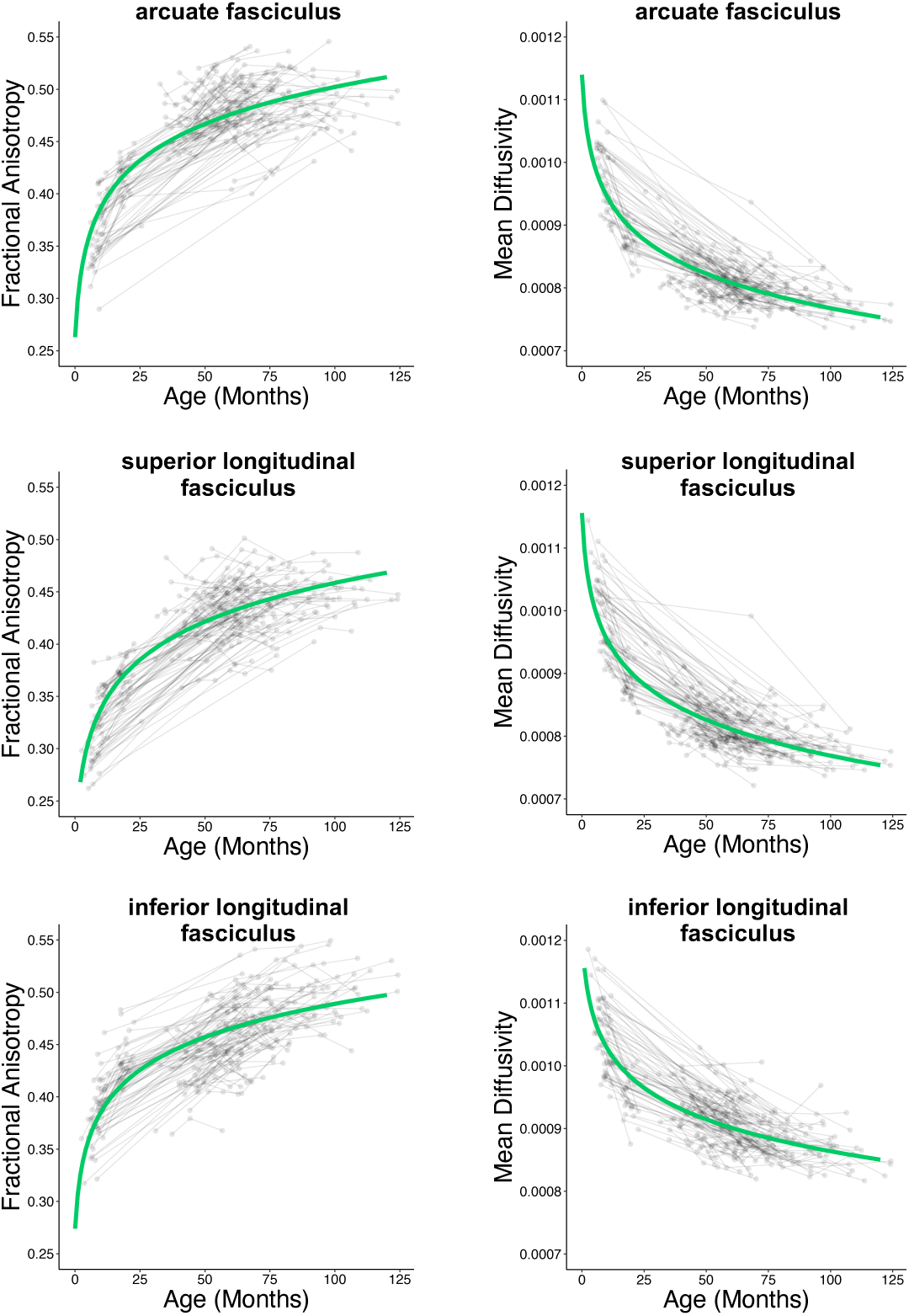
Longitudinal trajectories of fractional anisotropy and mean diffusivity between infancy and late childhood. Raw diffusion estimates (gray spaghetti plot backdrop) were submitted to linear mixed effects models using a logarithmic function. Individual growth curves predicted by this model were averaged to show the overall longitudinal trajectory of the sample (green line).

**Supplementary Figure 23.**
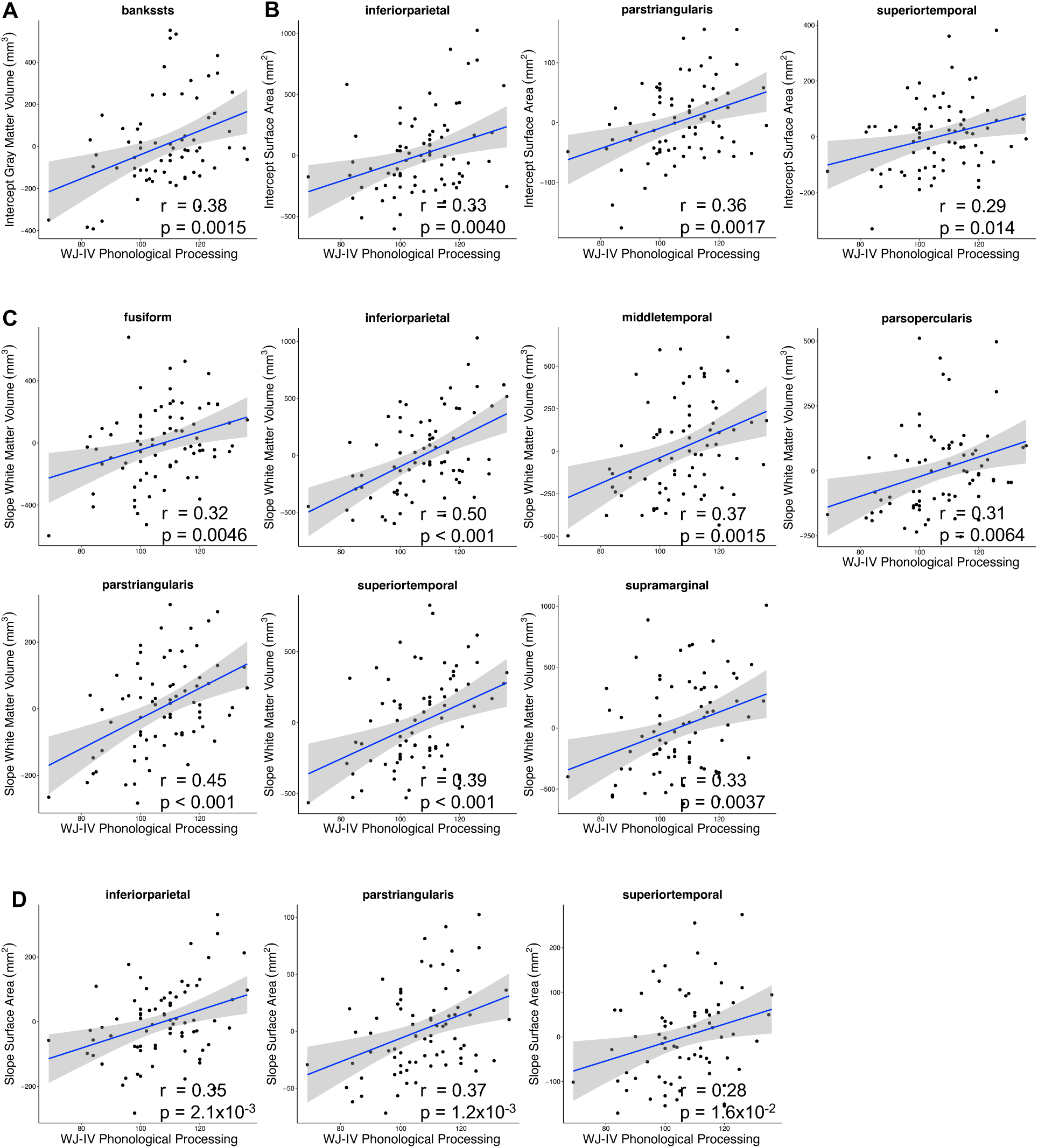
Statically significant associations between growth curve features of brain volume and surface area and preschool/early kindergarten phonological processing. (A) Gray matter volume in the left banks of the superior temporal sulcus exhibited an association between growth curve intercepts and phonological processing, whereas (C) all associations with white matter volume were between growth curve slopes and phonological processing. Both surface area intercepts (B) and slopes (D) exhibited associations with phonological phonological processing. All associations pass p_FDR_ < 0.05. WJ-IV, Woodcock-Johnson IV.

**Supplementary Figure 24.**
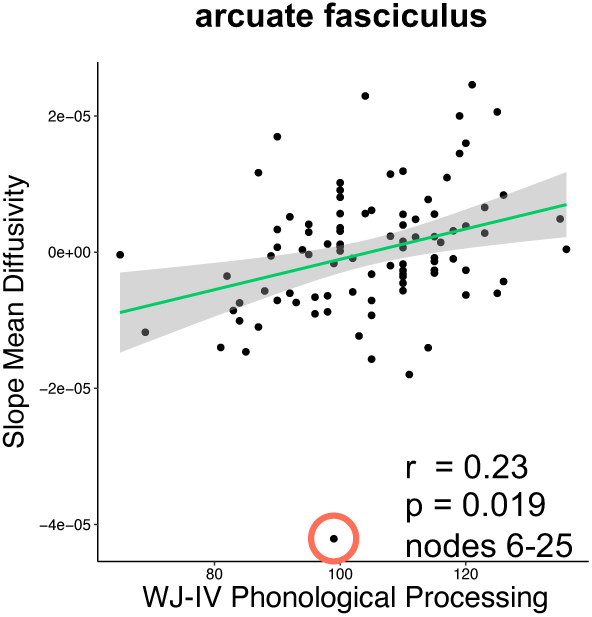
Statistically significant associations between growth curve features of white matter organization and preschool/early kindergarten phonological processing skill. Mean diffusivity in the left arcuate fasciculus exhibited associations between growth curve slopes and phonological processing. Association is cluster-level corrected at p_FWE_ < 0.05. Note: removal of outlier (orange circle) did not abolish cluster-level significance. WJ-IV, Woodcock-Johnson IV.

**Supplementary Figure 25.**
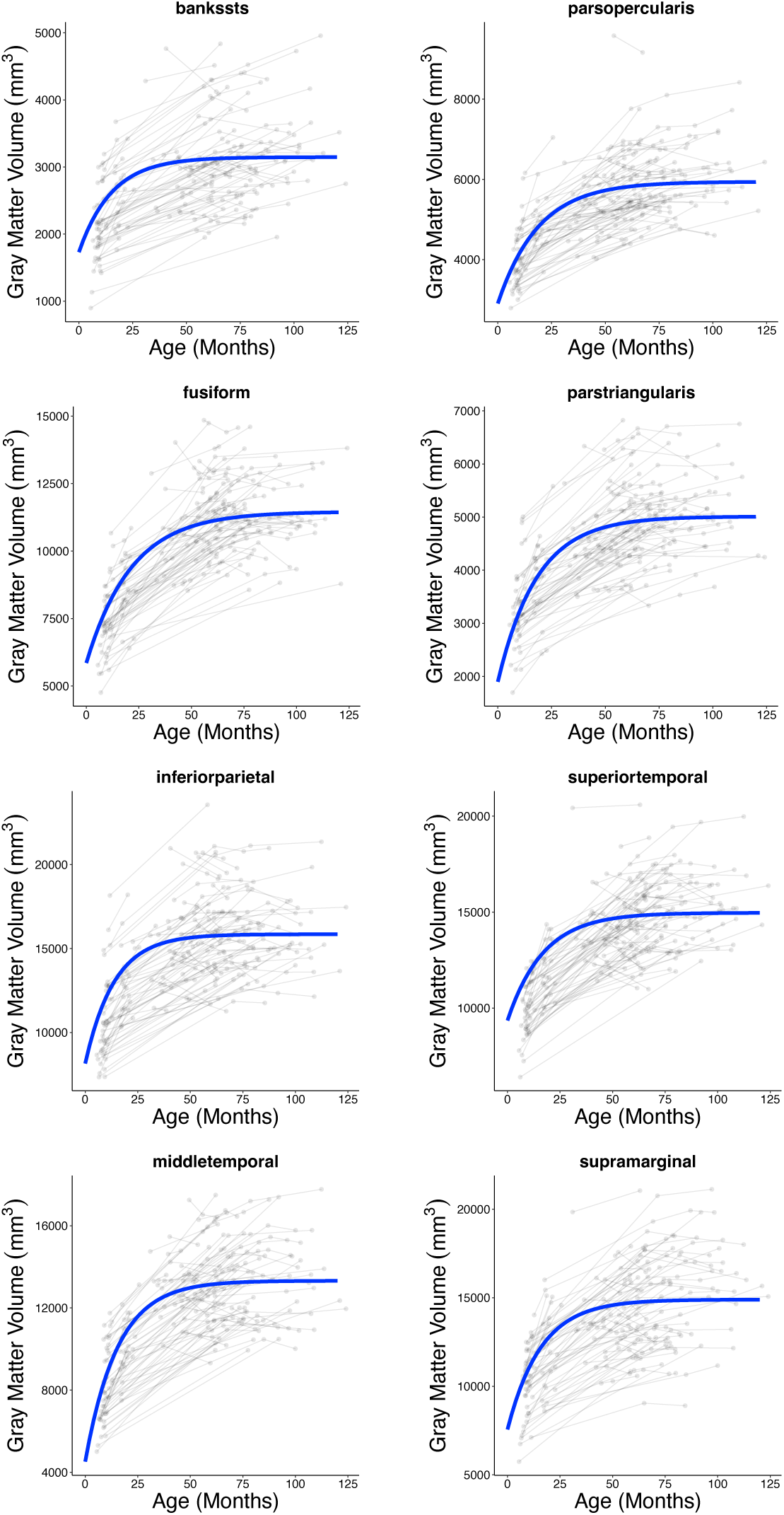
Longitudinal trajectories of gray matter volume between infancy and late childhood generated for sensitivity analyses. Raw estimates of gray matter volume (gray spaghetti plot backdrop) were submitted to nonlinear mixed effects models using an asymptotic

**Supplementary Figure 26.**
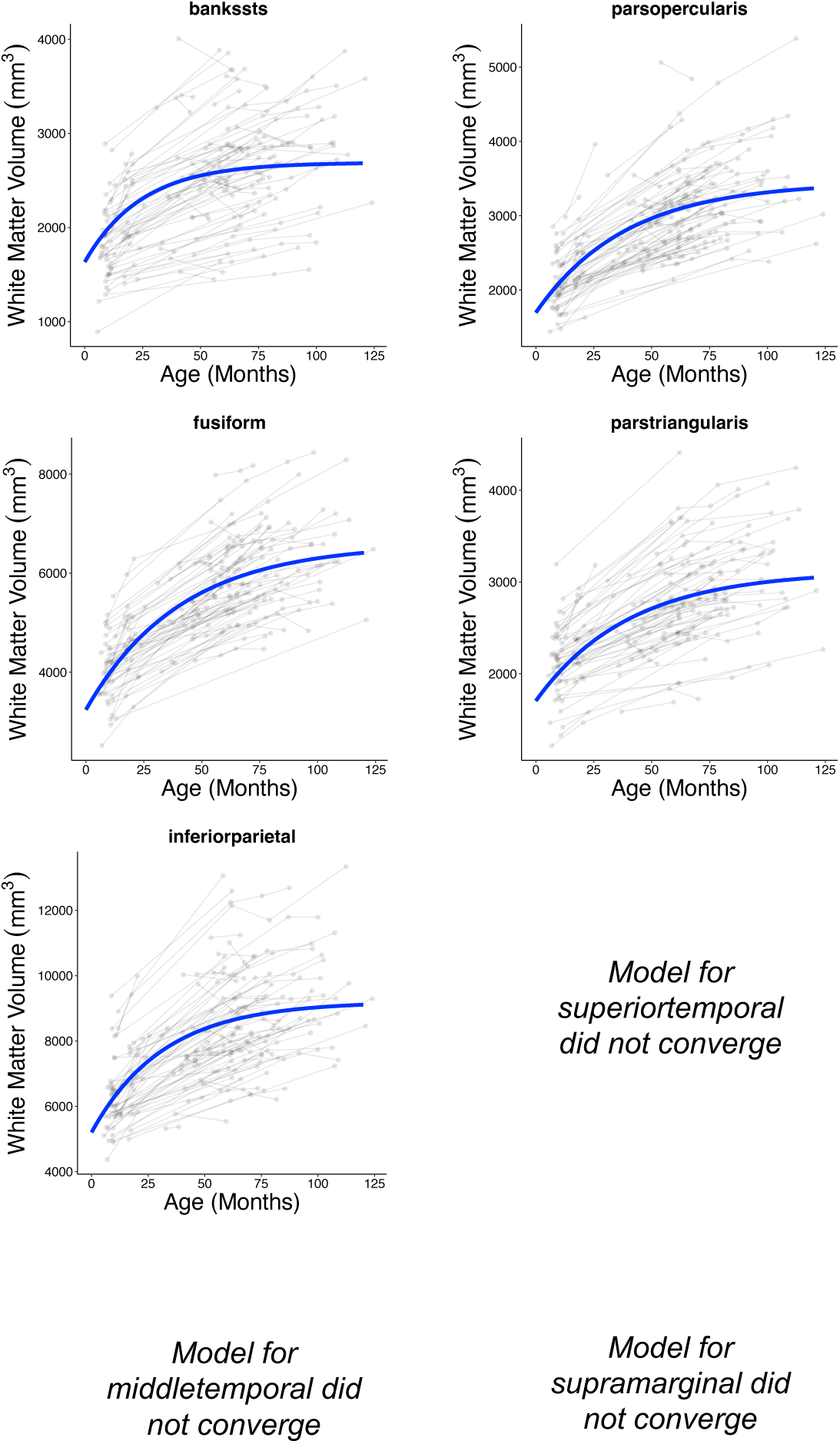
Longitudinal trajectories of white matter volume between infancy and late childhood generated for sensitivity analyses. Raw estimates of white matter volume (gray spaghetti plot backdrop) were submitted to nonlinear mixed effects models using an asymptotic

**Supplementary Figure 27.**
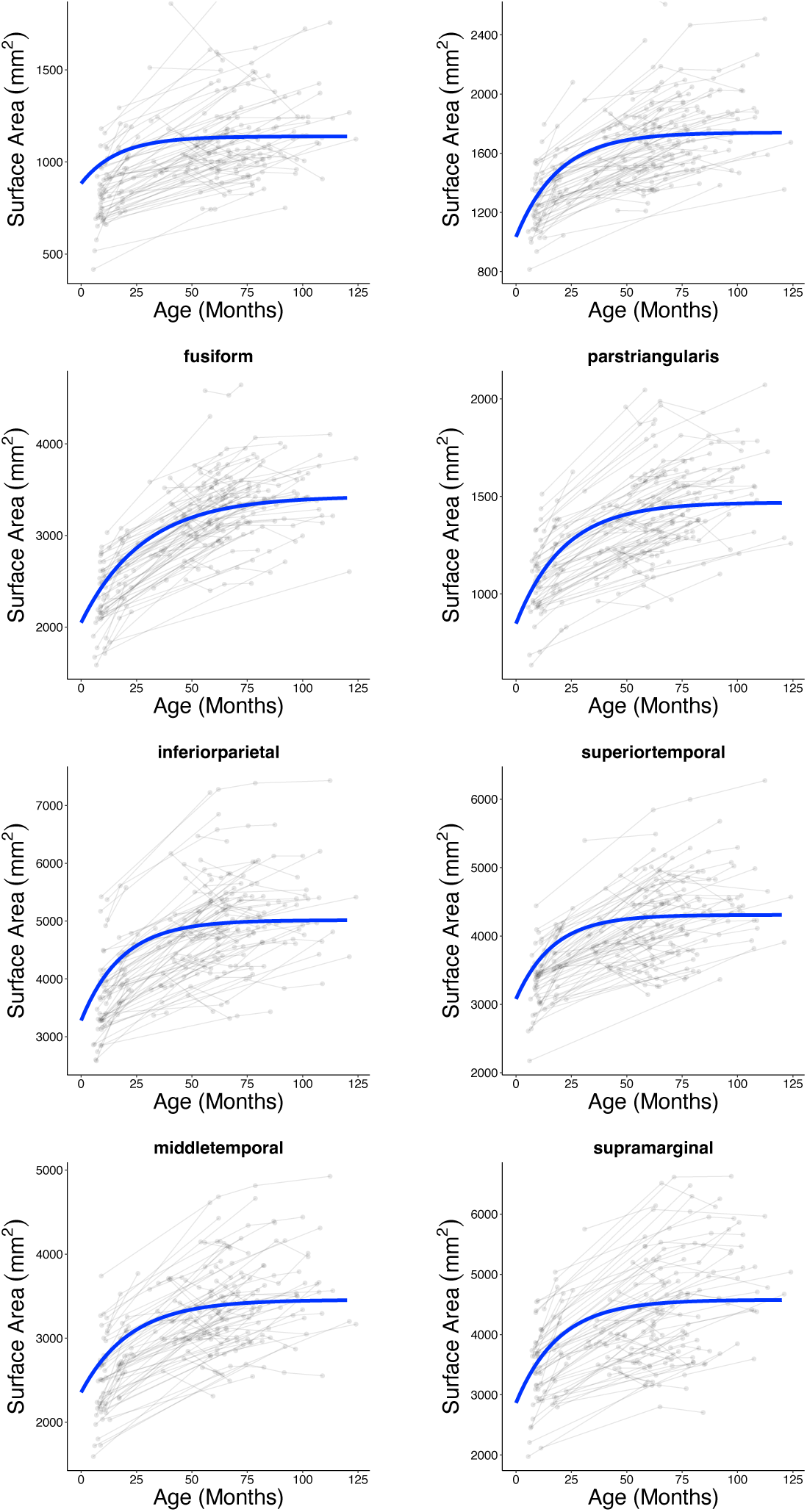
Longitudinal trajectories of surface area between infancy and late childhood generated for sensitivity analyses. Raw estimates of surface area (gray spaghetti plot backdrop) were submitted to nonlinear mixed effects models using an asymptotic function.

**Supplementary Figure 28.**
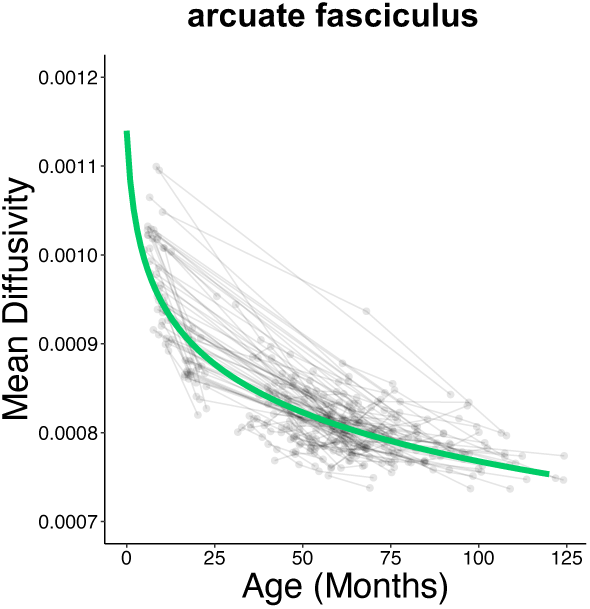
Longitudinal trajectories of mean diffusivity between infancy and late childhood generated for sensitivity analyses. Raw estimates of mean diffusivity (gray spaghetti plot backdrop) were submitted to a nonlinear mixed effects model using an asymptotic function. Individual growth curves predicted by this model were averaged to show the overall longitudinal trajectory of the sample (green lines).

**Supplementary Figure 29.**
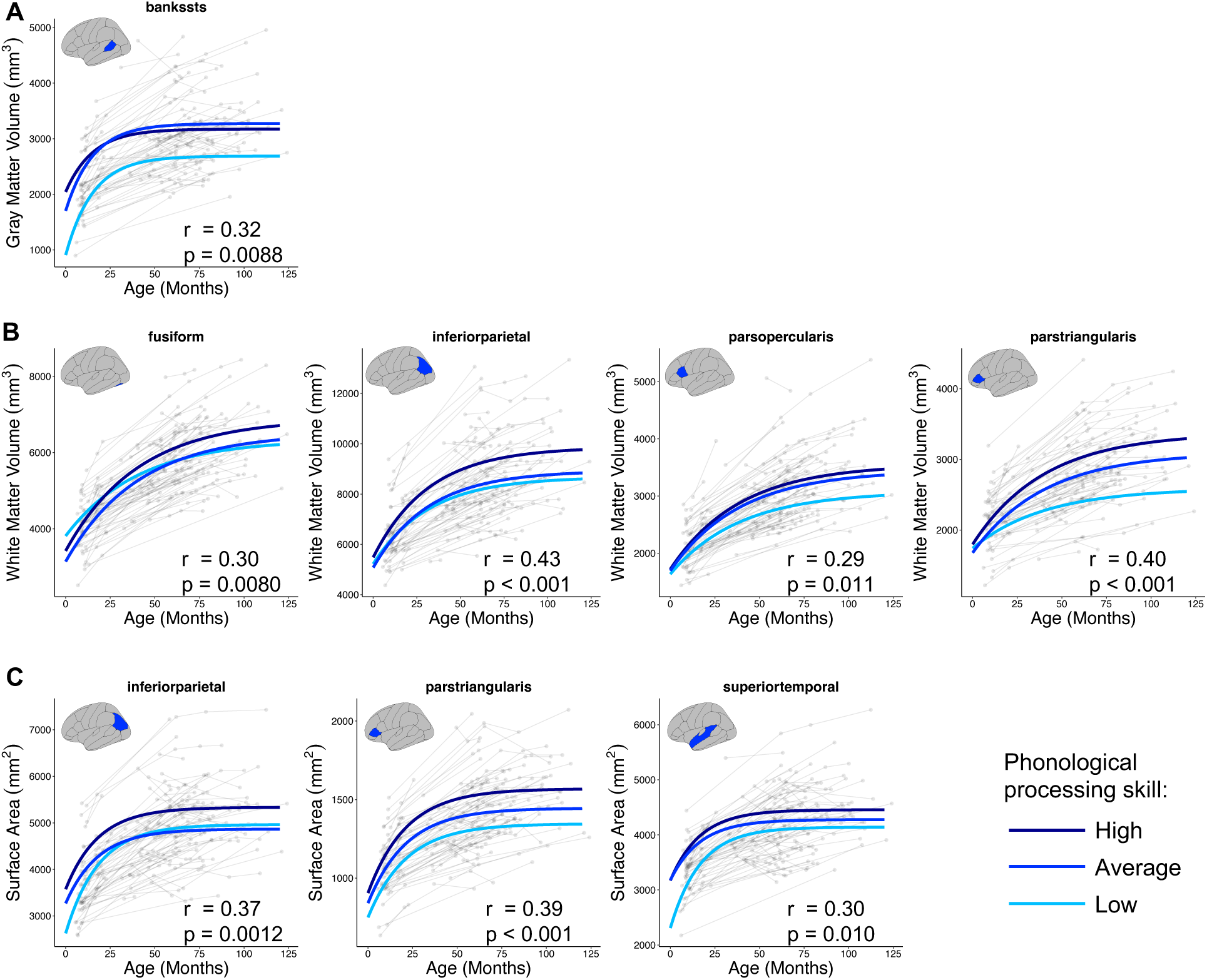
Longitudinal trajectories of brain structure from infancy to late childhood according to phonological processing skill in preschool/early kindergarten for sensitivity analyses. Graphs depict average trajectories for children with low (< 85), average (85–115), and high (> 115) standardized phonological processing scores for measures and regions whose (A) intercepts, (B) slopes, or (C) both intercepts and asymptotes correlated with phonological processing (p_FDR_ < 0.05). Correlation statistics are reported adjacent to their corresponding plots; intercept and asymptote statistics for surface area averaged here for visualization purposes but reported separately in Supplemental Table 3.

**Supplementary Figure 30.**
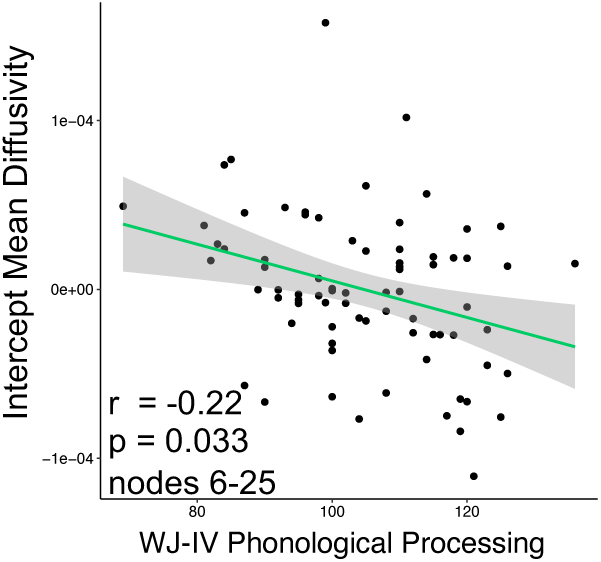
Statistically significant associations between growth curve features of white matter organization and preschool/early kindergarten phonological processing for sensitivity analyses. Mean diffusivity in the left arcuate fasciculus exhibited a negative association between growth curve intercepts and phonological processing. Association is cluster-level corrected at p_FWE_ < 0.05. WJ-IV, Woodcock-Johnson IV.

**Supplementary Figure 31.**
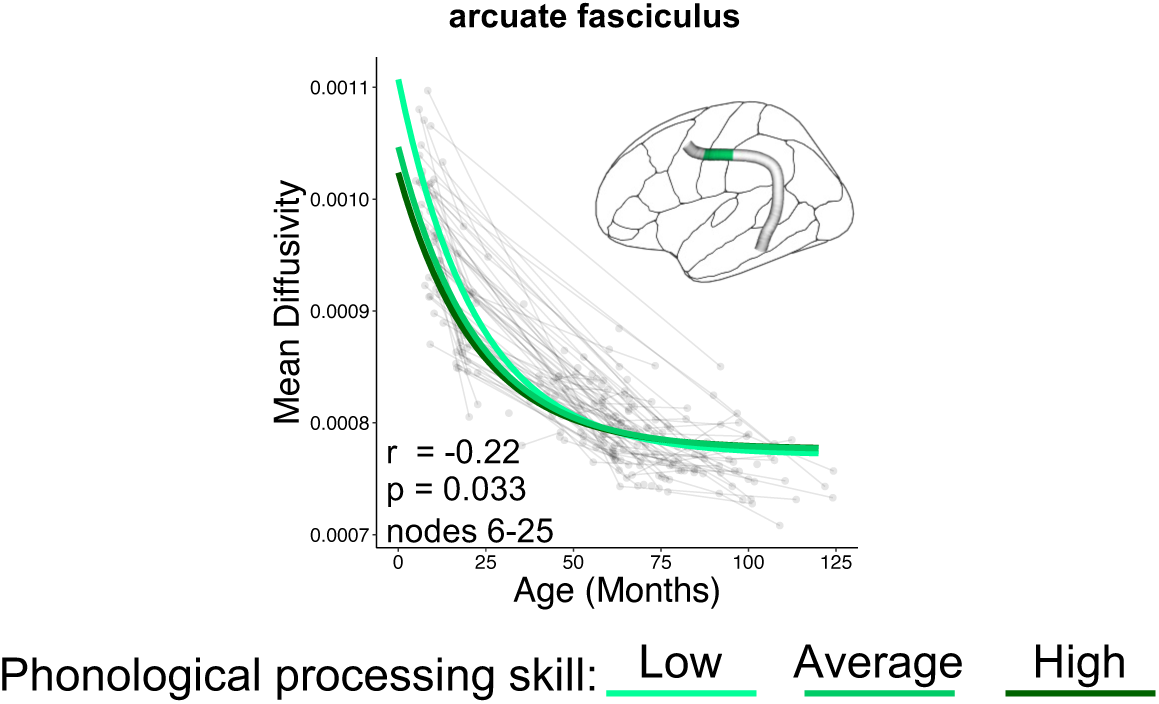
Longitudinal trajectories of mean diffusivity from infancy to late childhood according to phonological processing skill in preschool/early kindergarten for sensitivity analyses. Graph depicts average trajectories for children with low (< 85), average (85–115), and high (> 115) standardized phonological processing scores for the left arcuate fasciculus nodes whose intercepts correlated negatively with phonological processing (p_FWE_ cluster-level < 0.05).

**Supplementary Figure 32.**
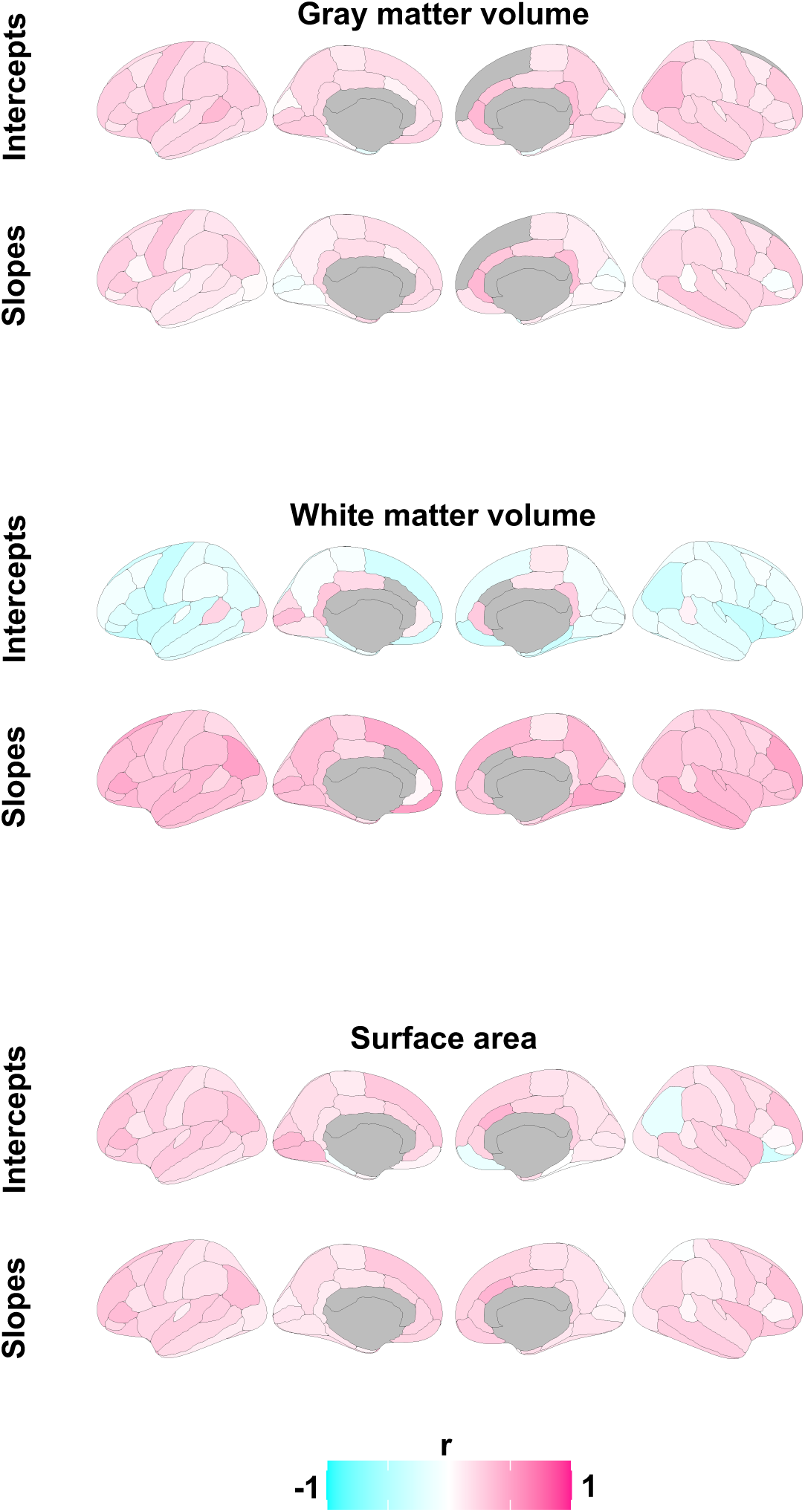
Maps of brain-behavior associations. Brain maps show associations between curve features (intercepts and slopes) and preschool/early kindergarten phonological processing across brain regions in the Desikan-Killiany atlas for gray matter volume, white matter volume, and surface area. Note: no model convergence for superior frontal volume and no white matter volume measures were available for anterior cingulate cortex.

